# Concerted expansion and contraction of immune receptor gene repertoires in plant genomes

**DOI:** 10.1101/2022.01.01.474684

**Authors:** Bruno Pok Man Ngou, Robert Heal, Michele Wyler, Marc W Schmid, Jonathan DG Jones

## Abstract

Recent reports suggest that cell-surface and intracellular immune receptors function synergistically to activate robust defence against pathogens, but whether or not they co-evolve is unclear. Here we determined the copy numbers of cell-surface and intracellular immune receptors in 208 species. Surprisingly, these receptor gene families contract and/or expand together in plant genomes, suggesting the mutual potentiation of immunity initiated by cell-surface and intracellular receptors is reflected in the concerted co-evolution of the size of their repertoires across plant species.

## Introduction

Plants have evolved a two-tier immune system that recognises and activates defence against pathogens^1^. Cell-surface pattern-recognition receptors (PRRs) recognise apoplastic and usually conserved pathogen-associated molecular patterns (PAMPs) and activate pattern-triggered immunity (PTI). Virulent pathogens secrete effector molecules into plant cells that suppress PTI and promote infection. Intracellular nucleotide-binding leucine-rich repeat (NLR) receptors recognise effectors and activate effector-triggered immunity (ETI). Although PTI and ETI were envisaged as two independent immune systems^1^, emerging evidence suggests they are inter-dependent and share multiple signalling components^2–5^. Thus, PTI and ETI function synergistically to provide robust immunity against pathogens. While PRRs and NLRs are functionally inter-dependent, co-evolution of these receptor gene families has not previously been investigated.

Plant PRR proteins are structurally diverse but are usually receptor-like kinases (RLKs) or receptor-like proteins (RLPs). RLKs carry extracellular ectodomains and cytosolic kinase domains, while RLPs lack cytosolic kinase domains. RLKs carry multiple types of extracellular domains, such as leucine-rich repeats (LRRs), lectins and lysM motifs (LysMs)^6^. LRR domain-containing RLKs (LRR-RLKs) and RLPs (LRR-RLPs) are the largest RLK- and RLP-gene families in plants^7^. Intracellular nucleotide-binding LRR (NLR) immune receptors carry NB-ARC domains with C-terminal LRR domains and N-terminal domains, usually comprising either coiled-coil (CC-), Toll/Interleukin-1 receptor/Resistance protein (TIR-) or RPW8-like coiled-coil (RPW8-) domains (hence, CC-NLRs (CNLs), TIR-NLRs (TNLs) and RPW8-NLRs (RNLs)^8^).

### Identification of cell-surface and intracellular immune receptors from plant genomes

To assess possible concerted expansion or contraction of PRR- and NLR-genes, we determined the size of these gene families in annotated proteomes from 208 publicly available genomes. These genomes include 25 algal species, 5 bryophyte species, 3 fern species and 171 angiosperms (3 basal angiosperms, 47 monocots and 118 eudicots) (Extended data figure 1a; Supplementary table 1). Genome sizes of these organisms range from 13 Mb - 14 Gb, with annotated protein count range from ∼5k - ∼300k (Supplementary table 1). To ensure consistency, we used the same pipeline to identify LRR-RLKs, LRR-RLPs, NB-ARCs and NLRs (with both NB-ARC and LRR domains) in each of these annotated proteomes (Supplementary figure 1; also see methods).

In total, we identified 59,809 LRR-RLKs, 13,942 LRR-RLPs, 61,023 NB-ARCs and 28,461 NLRs from 208 annotated proteomes (Supplementary figure 2; Supplementary table 2). As expected, the number of receptors varies enormously across 171 angiosperms, with LRR-RLKs ranging from 17 - 1567, LRR-RLPs ranging from 1 - 474, NB-ARCs ranging from 8 - 3542 and NLRs ranging from 0 - 827. To account for the effect of genome duplication and variable proteome sizes, we normalised these data using percentages (%) of LRR-RLKs, LRR-RLPs, NB-ARCs, NLRs from each proteome (number of identified genes divided by number of searched genes) (Supplementary figure 2; Supplementary table 3). After adjustment, LRR-RLKs range from 0.146 % - 2.56 %, LRR-RLPs range from 0.0111% - 1.10 %, NB-ARCs range from 0.0565 % - 4.58 % and NLRs range from 0% - 2.70% in 171 angiosperms (Figure 1; Supplementary table 3).

**Figure 1.**
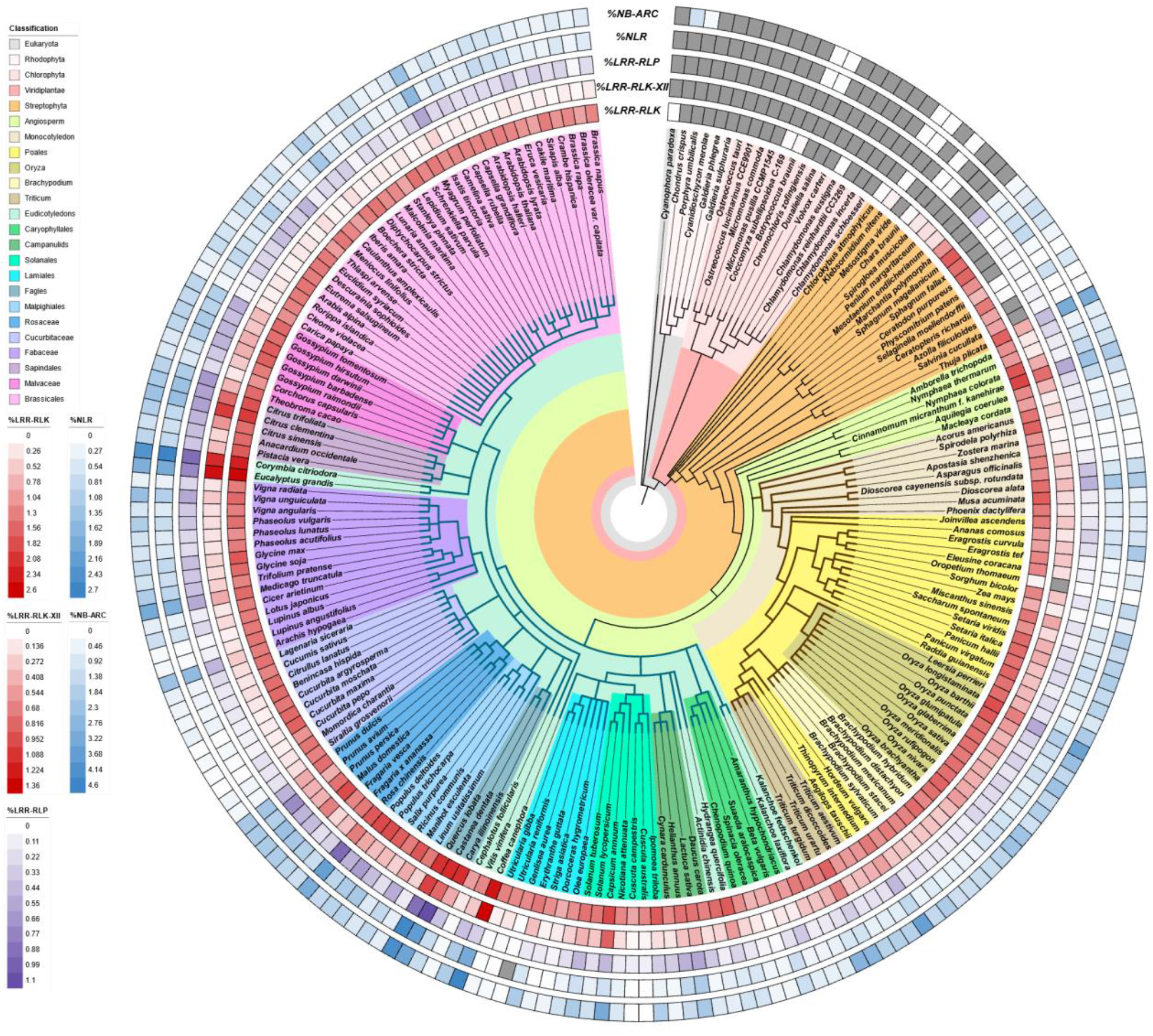
Immune receptor gene families in 208 plant genomes. Phylogenetic tree of 208 plant species, with heatmaps representing the percentages (%) of LRR-RLKs, LRR-RLK-XIIs, LRR-RLPs, NLRs and NB-ARCs in their corresponding annotated proteomes. Grey boxes in heatmaps indicate null values where no receptors were identified. Brown branches indicate monocots and green branches represent eudicots.

### Positive correlation between sizes of immune receptor families in plant genomes

Next, we determined the correlation between the percentages of PRRs (% LRR-RLKs and % LRR-RLPs) and NB-ARCs in angiosperms. Surprisingly, % NB-ARC and % LRR-RLPs show a strong positive linear correlation (Pearson’s r = 0.715), suggesting that NB-ARC and LRR-RLP gene families expand together (Extended data figure 3 and 4; supplementary data figure 7; supplementary table 4). On the other hand, % NB-ARC and % LRR-RLKs show a positive but weaker linear correlation (Pearson’s r = 0.598). We propose that PRRs involved in pathogen recognition are more likely to co-expand with NB-ARC gene families. This is consistent with the observation that characterized LRR-RLPs are usually involved in pathogen recognition, while LRR-RLKs can be involved not only in immunity but also in development and reproduction^9^.

To test if the NB-ARC gene family co-expands with PRRs specifically involved in pathogen recognition, we further classified LRR-RLKs into subgroups according to their kinase domains. LRR-RLKs can be classified into 20 subgroups, with each subgroup involved in different biological processes^10^ (Extended data figure 2). For example, BAK1 (BRI1-ASSOCIATED RECEPTOR KINASE) and other SOMATIC EMBRYOGENESIS RECEPTOR-LIKE KINASES (SERKs) function as PRR co-receptors and belong to LRR-RLK-II^11^. BRASSINOSTEROID INSENSITIVE 1 (BRI1) is a receptor for brassinolide and belongs to LRR-RLK Xb^12^. Members of LRR-RLK-XI are involved in recognition of self-peptides, such as PEP1 RECEPTOR 1 (PEPR1), CLAVATA 1 (CLV1) and RGF1 INSENSITIVE 1 (RGFR1)^9^. Members of LRR-RLK-XII, such as FLAGELLIN-SENSITIVE 2 (FLS2), EF-TU RECEPTOR (EFR) and Xa21^6^, are involved in detecting pathogen-derived molecules (Extended data figure 2; Supplementary figure 3). Across 208 species, LRR-RLK-XII form the largest LRR-RLK subgroup, followed by LRR-RLK-III and LRR-RLK-XI (Supplementary figure 4 and 5).

Next, we determined the correlation between % LRR-RLK from different subgroups and % NB-ARC in angiosperms (Extended data figure 3 and 4; Supplementary table 4). Strikingly, only 3 out of 20 LRR-RLK subgroups show significant and positive linear correlation with % NB-ARCs (LRR-RLK-VIII_1, LRR-RLK-VIII_2 and LRR-RLK-XII). Furthermore, LRR-RLK-XII forms much stronger positive correlation with % NB-ARCs (Pearson’s r = 0.826) compared to LRR-RLK-VIII_1 or LRR-RLK-VIII_2 (Pearson’s r = 0.329 and 0.443, respectively) (Figure 2a; Supplementary figure 7; Supplementary table 4). While LRR-RLKs involved in pathogen recognition are predominantly in subgroup XII, some members from LRR-RLK-VIII are also involved in immunity and pathogen recognition, such as CANNOT RESPOND TO DMBQ 1 (CARD1/HPCA1), HDS-ASSOCIATED RLK1 (HAK1) and LysM RLK1-interacting kinase 1 (LIK1)^13–16^ (Extended data figure 2). Since LRR-RLK-XII forms the largest LRR-RLKs subgroup, we tested if the positive correlation between % LRR-RLK (total) and % NB-ARC is predominantly caused by subgroup XII. Indeed, % LRR-RLK (all subgroups excluding XII) does not show any correlation with % NB-ARC (Pearson’s r = -0.0546). On the other hand, % LRR-RLP combined with % LRR-RLK-XII show strong positive correlation with % NB-ARC (Pearson’s r = 0.857) (Figure 2). Thus, PRR gene families specifically involved in pathogen recognition co-expand or co-contract with NB-ARC gene families.

**Figure 2.**
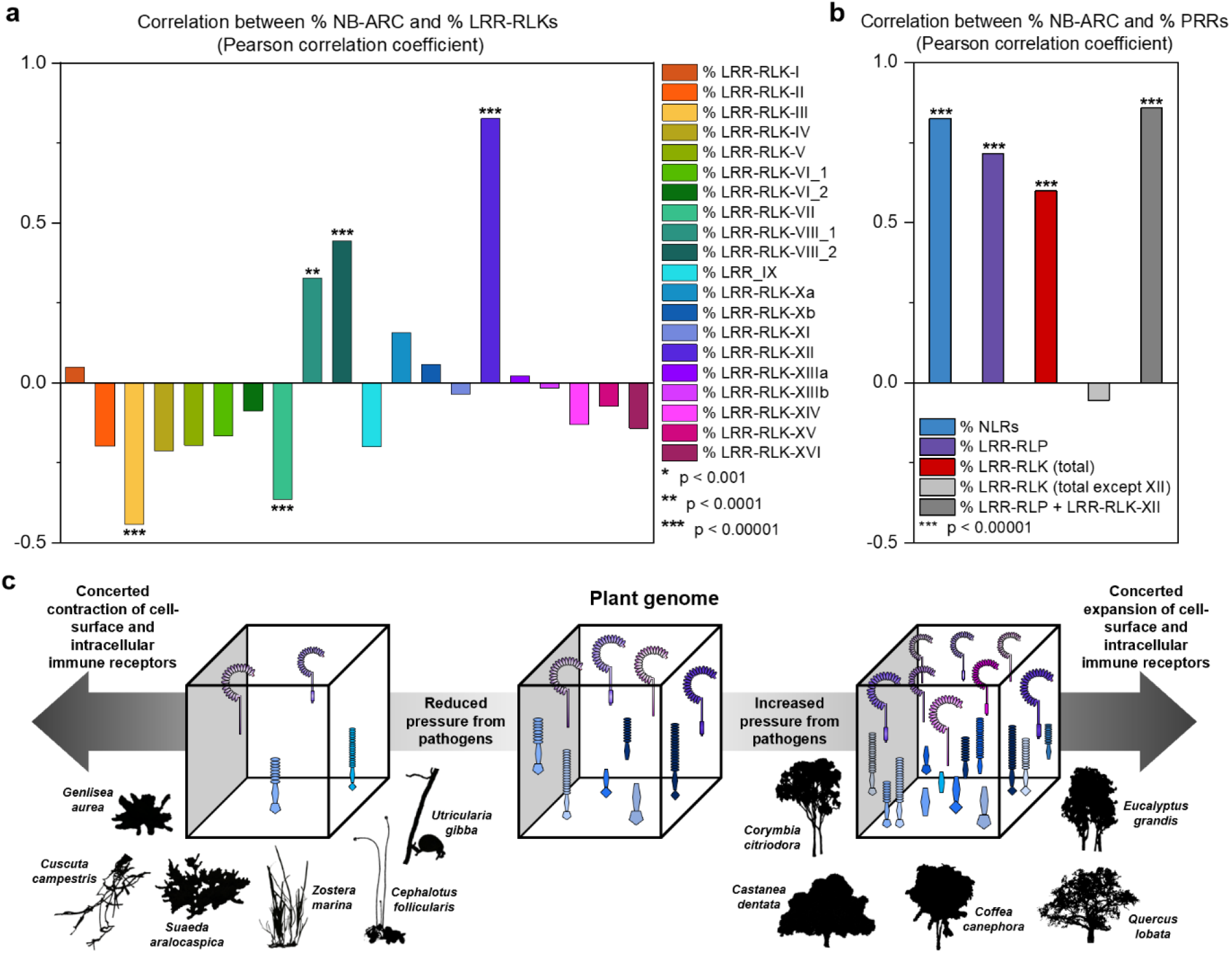
Concerted expansion and contraction of cell-surface and intracellular immune receptor genes in plant genomes. **a-b**, Correlation between % NB-ARC, % LRR-RLKs from different subgroups and % PRRs of different combinations. Bar chart represents the Pearson correlation coefficient, with significant values indicated with *. **c**, Schematic illustration of the co-expansion and co-contraction of immune receptors in plant genomes. We hypothesise that under increased pressure from pathogens, both PRR- and NLR-gene families expand and with reduced pressure from pathogens, PRR- and NLR-gene families contract together (see text).

### Expansion and contraction of PRR- and NLR-gene families in plants adapted to particular ecological niches

We observed a strong linear correlation between % NB-ARC and % LRR-RLK-XII and % LRR-RLP. Next, we checked if NLR gene family contraction coincides with PRR gene family contraction in organisms adapted to specific lifestyles, such as parasitism and carnivorism. The *Alismatales* and *Lentibulariaceae* lineages show a reduction in the size of NLR gene repertoires^17^ and species from these lineages also have low % PRRs (% LRR-RLP and % LRR-RLK-XII). These include *Genislea aurea, Zostera marina, Utricularia gibba, Utricularia reniformis* and *Spirodela polyrhiza* (0.0143 - 0.187 %; median in 171 angiosperms: 0.459 %) (Extended data figure 5; supplementary table 3; supplementary figure 8). We can infer that the % NB-ARC and % LRR-RLK correlation is not just due to co-expansion, but also co-contraction.

Carnivorous, aquatic and parasitic plant genomes carry few NLRs^17,18^. We tested if the number of cell-surface immune receptors is also reduced in these plants. Compared to monocots and core eudicots, species with these lifestyles have lower % NB-ARC, % LRR-RLK-XII and % LRR-RLP (Extended data figure 5c; supplementary figure 8). Notably, % LRR-RLK (total) in these groups is similar to monocots and core eudicots, as are most other LRR-RLK subgroups (Supplementary figure 8). It has been proposed that the reduced root system in parasitic and carnivorous plants result in fewer interactions or entry routes for pathogens^18^. Similarly, partial or complete submersion of aquatic species results in reduced exposure to airborne pathogenic spores, removing an interface for interaction with pathogens. Whilst still showing low receptor number, *Nymphaea thermarum, Nymphaea colorata* and *Spirodela polyrhiza* have greater % NB-ARC, % LRR-RLK-XII and % LRR-RLP values compared to three other submerging aquatic species (*Genlisea aurea, Utricularia gibba* and *Zostera marina*) (Extended data figure 5b-c). The larger immune receptor families in these floating aquatic species might be due to greater potential exposure to rapidly evolving airborne pathogens compared to submerged aquatic species.

Some other species and genera also show lower % NB-ARC, % LRR-RLK and % LRR-RLP. For example, the *Cucurbitaceae* show far fewer immune receptors than the neighbouring *Fabaceae* or *Rosaceae* clades (Figure 1; supplementary figure 6; supplementary table 3). Remarkably, in the monocot species *Oropetium thomaceum*, we observed only 0.0688 % NB-ARC containing proteins and no LRR-RLK-XII. This contrasts the other members of the *Poales*, where high % PRRs and % NB-ARCs are more frequent (Figure 1, supplementary table 3). *O. thomaeum* is an atypical member in the *Poales*. This drought-tolerant resurrection grass has the smallest known grass genome (245 Mb) and can survive losing 95 % of cellular water^19^. Despite its small genome, *O. thomaeum* has similar number of predicted proteins as other *Poales* species such as *Ananas comosus, Oryza longistaminata* and *Triticum urartu*, suggesting that the small immune receptor families could be independent of the observed genome reduction (Extended data figure 1).

On the other hand, some plant groups show much larger immune receptor families. Many species of the order *Poales* show high % LRR-RLK, % LRR-RLP and % NB-ARC, most notably in the *Oryza* and *Triticum* genera (Figure 1; supplementary figure 6). Expansion and diversification of immune receptors in these species may be driven by high pathogen pressure. Grasses are typically grown in high densities and are frequently challenged by rust and blast species which produce numerous, wind-dispersed spores with high genetic diversity. Genetic exchange by sexual reproduction and somatic hybridisation drives the emergence of new virulent strains^20^ such as the *Ug99* strain of the wheat stem rust pathogen *Puccinia graminis f. sp. tritici*^21^. The expanded repertoire of the immune receptors and increased heterogeneity could be a result of high pressure from these pathogens. In addition, many tree species also show a high proportion of PRR and NB-ARC proteins in their proteomes. These include *Eucalyptus grandis, Castanea dentata, Corymbia citriodora, Quercus lobata, Coffea canephora, Cinnamomum kanehirae, Prunus avium, Malus domestica, Theobroma cacao* and *Citrus* species (Extended data figure 5b-c; supplementary table 3). Thus, some plant lifestyles might also correlate with expansion of immune receptor gene families. Annual plants are subject to shorter periods of pathogen pressure before reproduction, whereas biennial or perennial plants, especially trees, must survive for much longer. Conceivably, this long-term pathogen pressure drives the expansion of immune receptor gene families.

### Positive correlation between the sizes of immune receptor families is not due to genomic clustering

Previously, analyses of the *Solanum lycopersicum* and *Arabidopsis thaliana* genome have suggested that NLRs, RLPs and RLKs might form genomic clusters^22,23^. Genomic clustering could mean that expansion/contraction of a gene family could result in genes in close proximity indirectly expanding in tandem. To determine if concerted expansion/contraction of immune receptor families is due to genomic clustering, we investigated the *S. lycopersicum, S. tuberosum* and *O. sativa* genomes. In all three genomes, many LRR-RLK-XII and LRR-RLP loci overlap with NB-ARC encoding loci (Supplementary figure 9). To quantify this, we calculated the average distance of LRR-RLKs and LRR-RLPs to the closest NB-ARC encoding genes and compared to a distribution of randomly selected genes (see methods for details). Both LRR-RLK-XIIs and LRR-RLPs are more closely linked to NB-ARC genes than randomly selected genes. However, LRR-RLK-III and LRR-RLK-XI genes are also closely linked to NB-ARC genes (Extended data figure 6-7). Since % LRR-RLK-III and % LRR-RLK-XI does not show positive correlation with % NB-ARC, we conclude that whilst NB-ARC-encoding genes can form genomic clusters adjacent to LRR-RLK-XIIs, the co-expansion/contraction of these immune receptors is likely to be caused by mechanisms other than genomic clustering.

## Discussion

Previously it was shown that cell-surface and intracellular immune systems exhibit mutual potentiation and inter-dependency^2–5^. Here, we show that in addition to their functional relationship, there is also an evolutionary correlation between numbers of cell-surface and intracellular immune receptors. Expansion and/or contraction of intracellular NLRs coincides with expansion and/or contraction of cell-surface PRRs involved in pathogen recognition (Figure 2c). We propose that pathogen pressure shapes the immune receptor diversity and repertoire, which, as a result, is determined by plant lifestyles and their ecological niches. Since parasite pressure drives the retention of sexual reproduction that reshuffles immune receptor alleles each generation^24^, inbreeding species may require an increased number of immune receptors compared to their outbreeding ancestors, an outcome that can also result from polyploidy. Since the concerted expansion and contraction of immune receptors in plant genomes are not due to genomic clustering, further study is needed to understand the mechanism(s) underpinning these observations. As functionally inter-dependent genes often co-expand/contract together, it is likely that the functional relationship between cell-surface and intracellular immune receptors is conserved across plant species.

## Methods

### LRR-RLK identification

Protein sequences from all 208 plant proteomes were first filtered for the longest protein isoforms and unique protein sequences by removing shorter sequences that were perfectly identical to another longer sequence with usearch^25^ (version v10.0.240_i86linux32, options -sort length -id 1.0 -cluster_fast - centroids,). Sequences shorter than 300 amino acids were removed as they are unlikely LRR-RLKs. The remaining proteins were searched for the presence of a protein kinase domain (PFAM PF00069.26) and an LRR domain (PFAM PF18805.2, PF18831.2, PF18837.2, PF00560.34, PF07723.14, PF07725.13, PF12799.8, PF13306.7, PF13516.7, PF13855.7, PF14580.7, PF01463.25, PF08263.13, and PF01462.19) with hmmer^25^ (version 3.1b2, options -E 1e-10 for the kinase domain and -E 10e-3 for the LRR domains). The Arabidopsis sequences which were previously classified into 20 LRR-RLK subgroups^10^ were filtered likewise for the presence of LRR and kinase domains. 19 Sequences were thereby removed. To classify all candidate sequences according to the Arabidopsis subgroups, the highest scoring kinase domain region of each candidate was extracted and aligned to the Arabidopsis reference sequences using diamond^26^ (version 0.9.26, options -e 1e-10 -k 300).

### Phylogeny

The phylogeny of each subgroup was inferred using the kinase domains. Sequences were aligned with FAMSA^27^. Alignments were not trimmed^28^ and phylogenetic trees were inferred with FastTree^29^ (version 2.1.11 SSE3, option -lg). Trees were rooted with gotree^30^ (v0.4.2) using the sequences belonging to the most basal species as outgroup (according to the taxonomic tree).

### NLR identification

NLR receptors were identified using the same set of proteins as for LRR-RLK identification. Proteins were searched for the presence of LRR domains and the absence of a kinase domain (hmmer options as above), as well as the presence of NB-ARC (PF00931.23) domains (hmmer option -E 1e-10 for NB-ARC).

### LRR-RLP identification

LRR-RLPs were identified similarly but filtering for proteins of a minimal length of 250 bp first. Proteins were then searched for the presence of LRR domains and the absence of a kinase domain (hmmer options as above), as well as the presence of a C3F domain (hmmer option -E 1e-10 and a minimal alignment length of 140). The hmmer-profile for the C3F domain was obtained from a multiple alignment^31^) of Arabidopsis LRR-RLPs^32^ (Steidele and Stam (2021)). The domain was trimmed manually, starting from the conserved Y in the C2 domain (Figure 6B in Fritz-Laylin et al^33^). Candidates were finally filtered for the presence of a transmembrane domain using tmhmm^34^ (version 2.0).

### Test for co-occurrence of NB-ARC, LRR-RLKs and LRR-RLPs

To test whether two gene groups are closer to each other than expected by chance, we used a test based on random sampling. For example, group A (LRR-RLK-XII) with n and group B (NLRs) with m genes. The observed distance was calculated as the average closest distance between genes in group A and genes in group B. A distribution for the expected distance was obtained by random sampling m genes and calculating the average closest distance of genes in group A to the genes in the random set (1,000 times). Genes were sampled from the list of genes that was used to search for the genes in group B (see Supplementary Figure 1).

### Taxonomic tree

The taxonomic tree was obtained from NCBI (https://www.ncbi.nlm.nih.gov/Taxonomy/CommonTree/wwwcmt.cgi). Phylogenetic tree of the 208 species are generated by phyloT (https://phylot.biobyte.de/) based on NCBI taxonomy database. Phylogenetics trees are visualised and figures are generated by iTOL^35^.

### Correlation analysis

Pearson’s correlation analysis were performed with OriginPro (https://www.originlab.com/).

## Supporting information

Supplementary Table 1. List of species included in this study

Supplementary Table 2. Number of receptor genes in 208 species

Supplementary Table 3. Percentages of receptor gene families in 208 species

Supplementary Table 4. Correlation analysis

## Acknowledgements

We thank Sebastian Fairhead, Detlef Weigel, Anna-Liisa Laine and Matthew Moscou for discussions and suggestions; and the Gatsby Foundation for funding to the J.D.G.J. laboratory. B.P.M.N was supported by the Norwich Research Park Biosciences Doctoral Training Partnership from the Biotechnology and Biological Sciences Research Council (BBSRC) (grant agreement BB/M011216/1).

## Author contributions

B.P.M.N., R.H. and J.D.G.J. conceived and conceptualized the study; M.W.S. and M.W. performed the bioinformatic analyses; B.P.M.N, M.W. and M.W.S. performed the statistical analyses; B.P.M.N. and R.H. wrote the original draft; and B.P.M.N., R.H. and J.D.G.J. reviewed and edited the manuscript.

## Competing interests

The authors declare no competing interests.

## Supplementary information

Supplementary figures 1-9 and supplementary tables 1-4 are included with the manuscript.

**Extended data figure 1.**
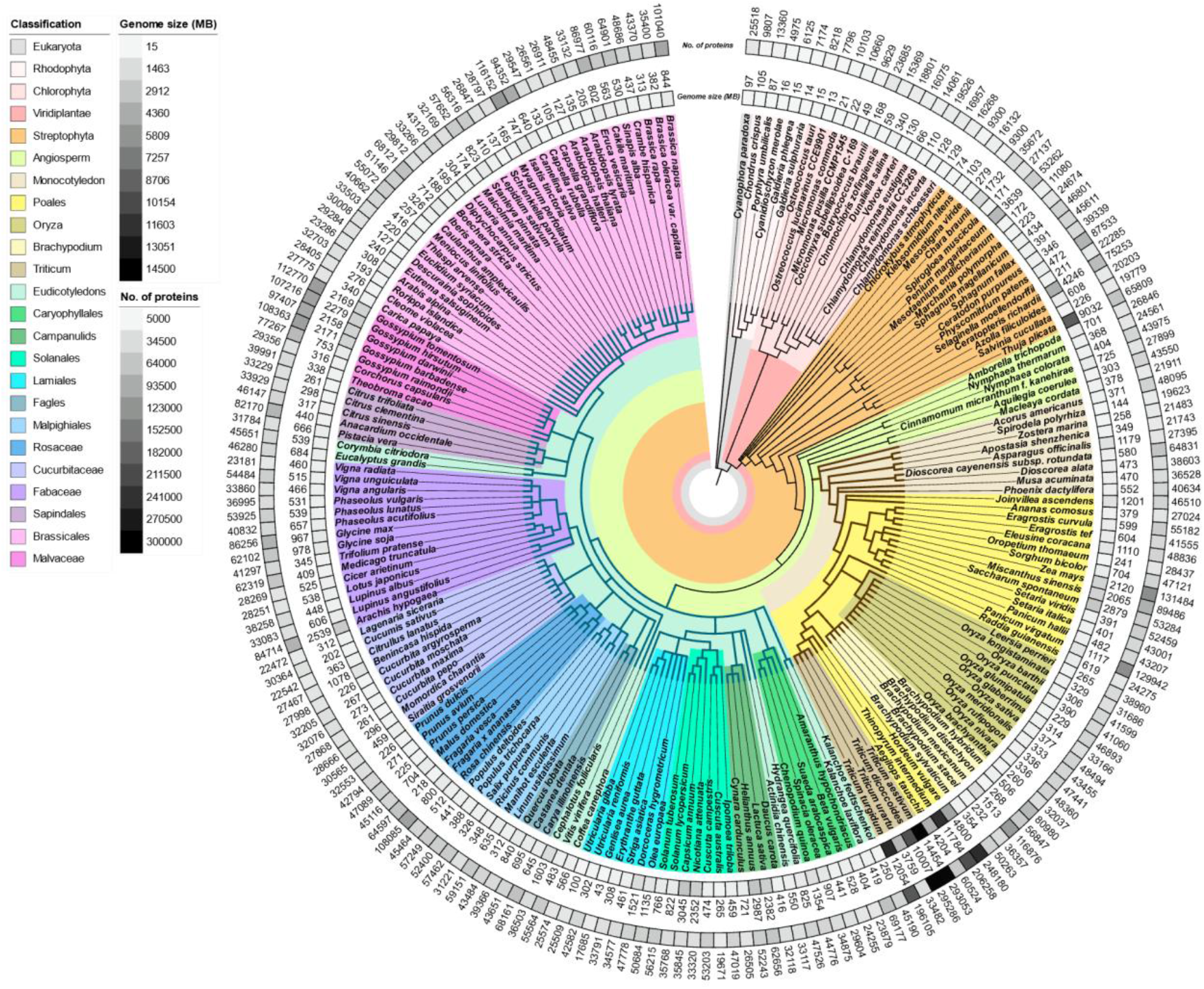
208 plant genomes used in this study. Phylogenetic tree of 208 plant species, with heatmaps representing the assembled genome size and number of annotated proteins. Brown branches indicates monocots and green branches represent eudicots.

**Extended data figure 2.**
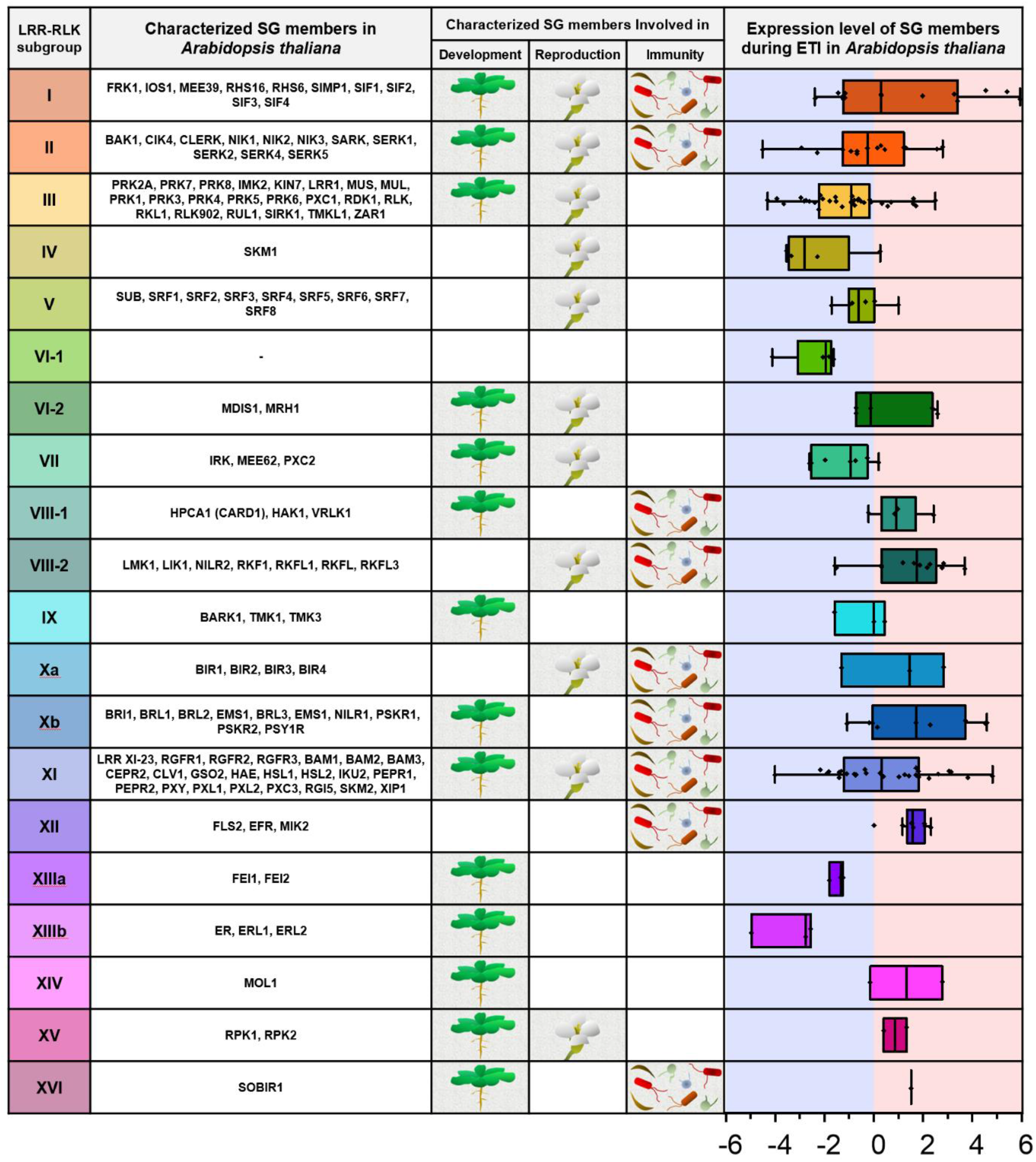
LRR-RLK subgroups in plants. Table representing the characterized subgroup members in *Arabidopsis thaliana*; the biological processes of which the characterized members are involved in, and expression of subgroup members during effector-triggered immunity (ETI). Red shade represents increased expression and blue shade represents decreased expression during ETI. X-axis values represents log_2_(fold change during ETI relative to untreated samples). RNA-seq data analysed here were reported previously^2^. ETI is activated by estradiol-induced expression of avrRps4 in *Arabidopsis thaliana* for 4 hours.

**Extended data figure 3.**
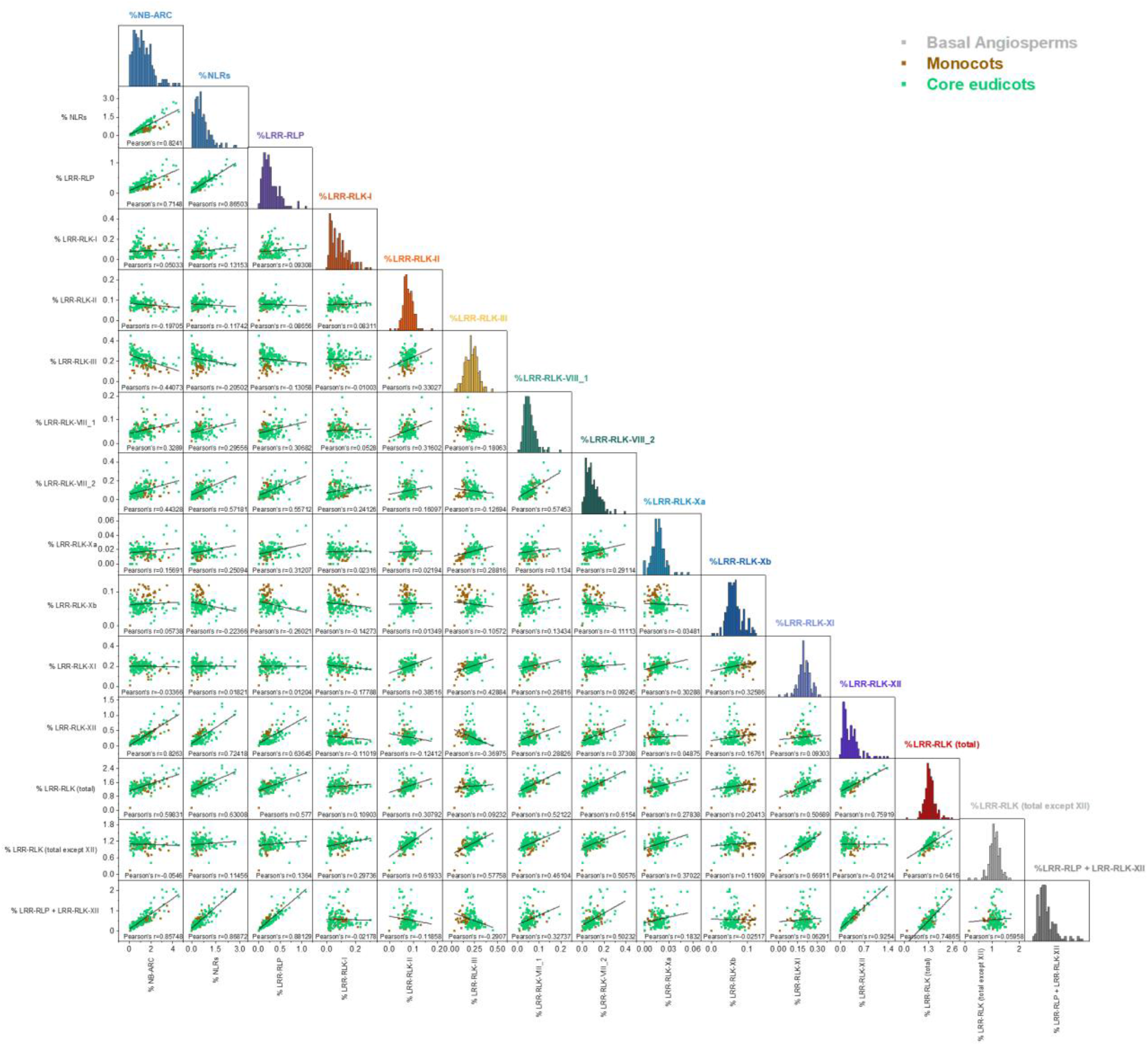
Scatter plot of Pearson correlation analysis between % NB-ARCs, % NLRs, % LRR-RLPs and % LRR-RLKs. Bottom left boxes include scatter plot between the corresponding % receptor-gene families in 171 angiosperms. Grey dots represent basal angiosperms (n=2), brown dots represent monocots (n=47) and green dots represent eudiots (n=118). The Pearson correlation coefficient (Pearson’s r) is indicated below each scatter plot. The diagonal boxes include the distribution of % receptor-gene families in 171 angiosperms.

**Extended data figure 4.**
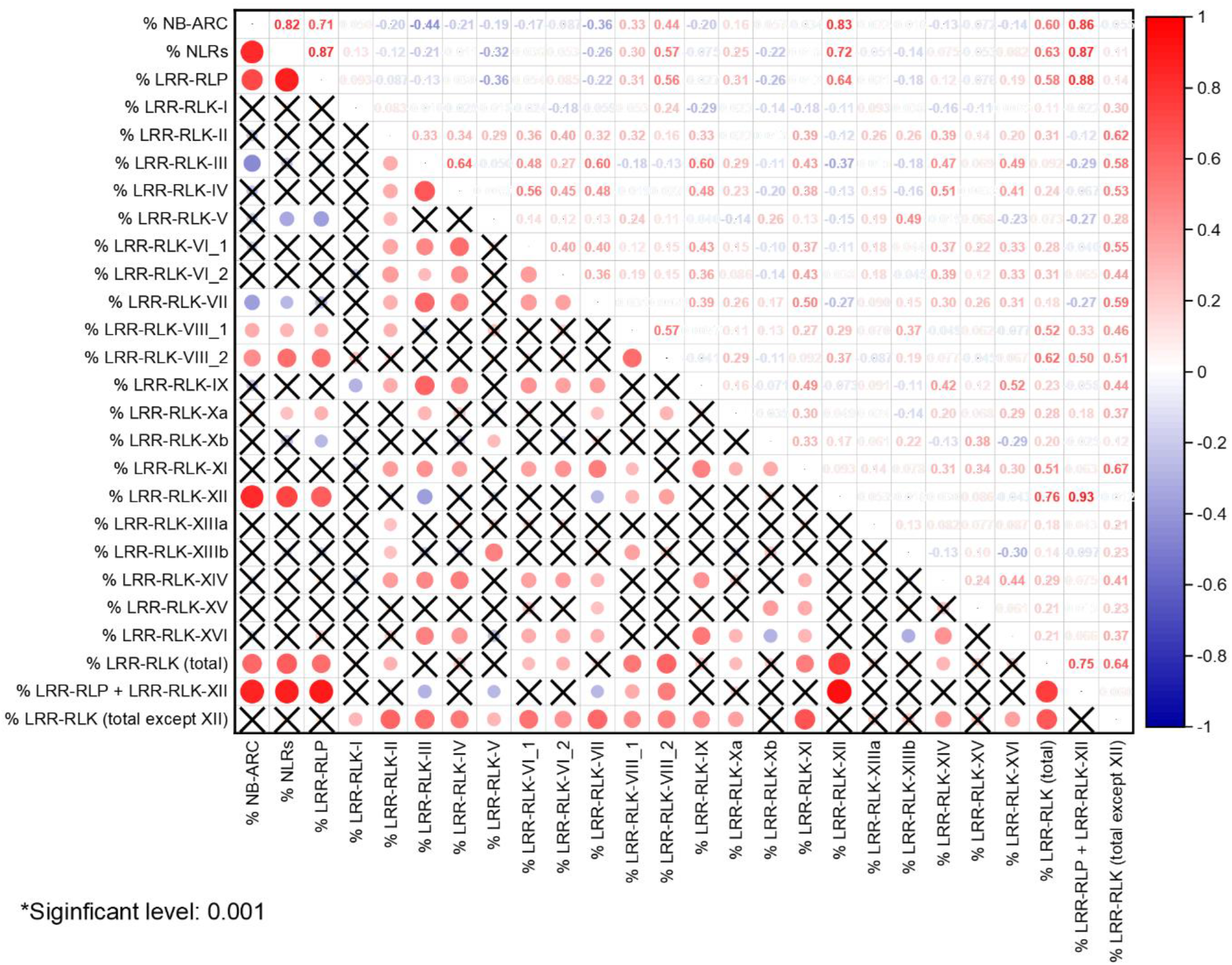
Pearson correlation plot of % NB-ARCs, % NLRs, % LRR-RLPs and % LRR-RLKs. Bottom left squares indicate the strength of correlation between the corresponding % receptor-gene families in 171 angiosperms. Red circles indicate significant and positive linear correlations, blue circles indicate significant and negative linear correlations and crosses (X) indicate insignificant correlations. The sizes of circles represent the strength of the correlations. P-values < 0.001 are considered as significant. Top right squares indicate the values of Pearson correlation coefficient between the corresponding % receptor-gene families in 171 angiosperms. Red values represent positive correlations and blue values represent negative correlations.

**Extended data figure 5.**
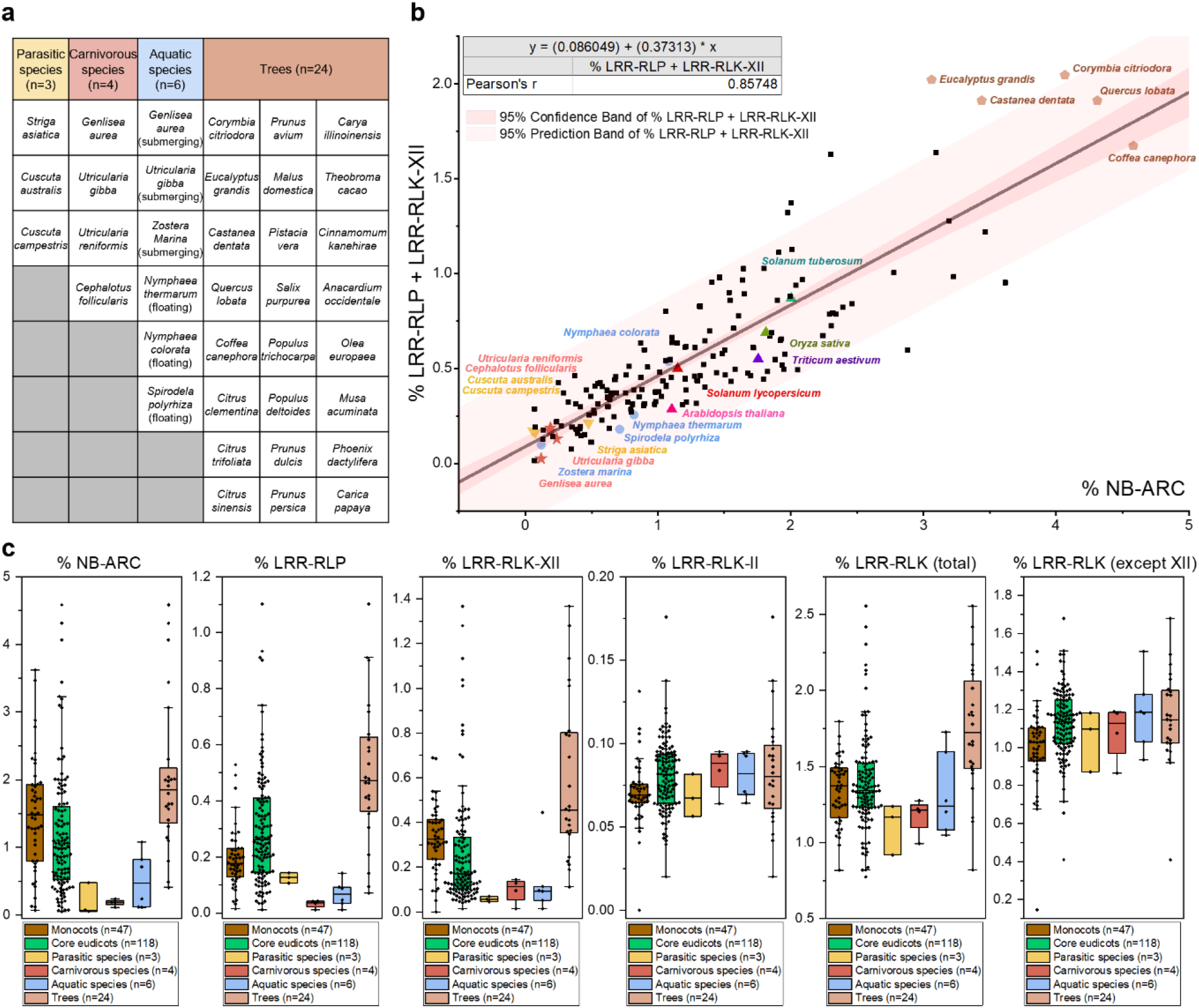
Expansion and contraction of PRR- and NLR-gene families in plants adapted to particular lifestyles and ecological niches. **a**. List of parasitic species, carnivorous species, aquatic species and trees used in this study. **b**. Scatter plot of % LRR-RLP + LRR-RLK-XII against % NB-ARC. % LRR-RLP + LRR-RLK-XII and % NB-ARC forms a strong and positive linear correlation. Parasitic species, carnivorous species, aquatic species and trees are indicated as yellow inverted triangles, orange stars, blue circles and brown pentagons. Model organisms are also indicated as triangles of different colours. **c**. Boxplot of % NB-ARC, LRR-RLP, LRR-RLK-XII, LRR-RLK-II, LRR-RLK (total) and LRR-RLK (total except XII) in monocots, eudicots, parasitic species, carnivorous species, aquatic species and trees.

**Extended data figure 6.**
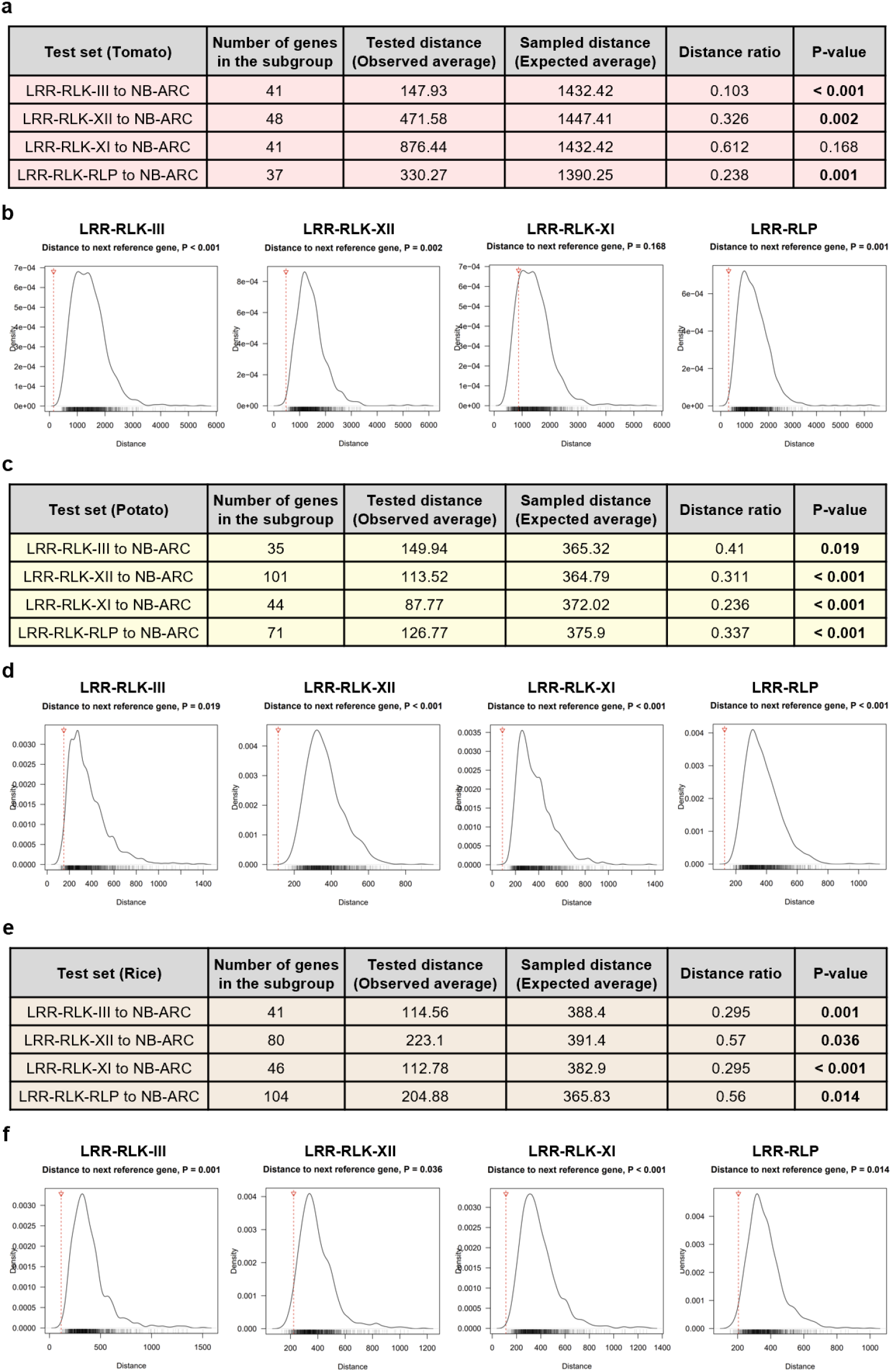
Genomic clustering of LRR-RLKs, LRR-RLPs and NB-ARCs in tomato, potato and rice. **a, c, e**. Table summarizing the statistical analysis of genomic clustering between PRRs and NLRs in *Solanum lycopersicum* (a), *Solanum tuberosum* (c) and *Oryza sativa* (e). The average distance between PRR gene family members and the next closest NB-ARC genes were calculated. This is then compared to a distribution (n=1000) of average distances between randomly-sampled genes and the next closest NB-ARC genes. Significant values are indicated in bold (p-value < 0.05 is considered as significant). **b, d, f**. Distribution (n=1000) of shortest average distances between randomly-sampled genes and the next closest NB-ARC genes in Solanum lycopersicum (b), Solanum tuberosum (d) and Oryza sativa (f). Red lines indicate the average distance between the corresponding PRR gene family members and the next closest NB-ARC genes.

**Extended data figure 7.**
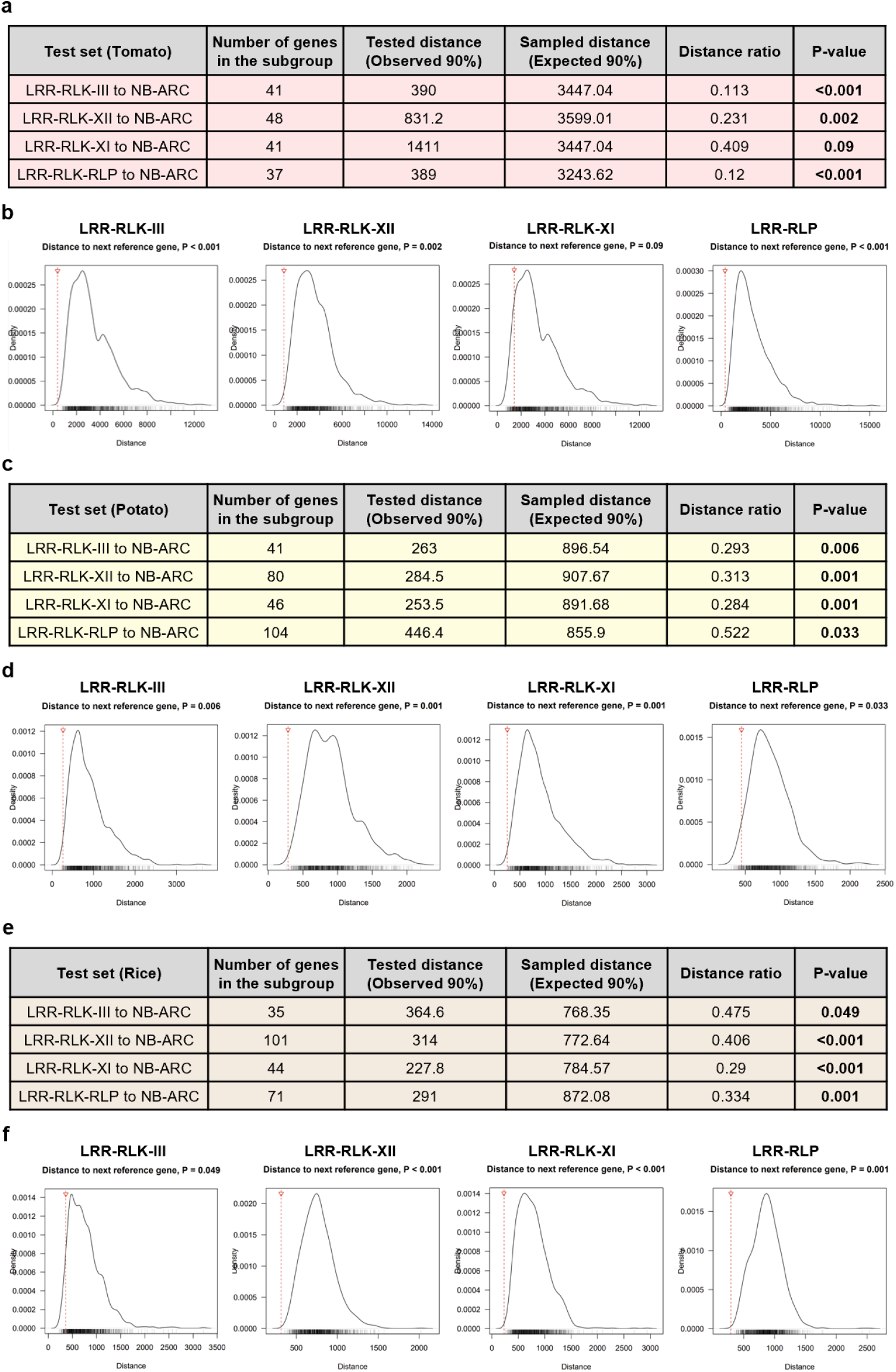
Genomic clustering of LRR-RLKs, LRR-RLPs and NB-ARCs in tomato, potato and rice. **a, c, e**. Table summarizing the statistical analysis of genomic clustering between PRRs and NLRs in *Solanum lycopersicum* (a), *Solanum tuberosum* (c) and *Oryza sativa* (e). The 90-percentile distance between PRR gene family members and the next closest NB-ARC genes were calculated. This is then compared to a distribution (n=1000) of 90-percentile distances between randomly-sampled genes and the next closest NB-ARC genes. Significant values are indicated in bold (p-value < 0.05 is considered as significant). **b, d, f**. Distribution (n=1000) of 90-percentile distances between randomly-sampled genes and the next closest NB-ARC genes in *Solanum lycopersicum* (b), *Solanum tuberosum* (d) and *Oryza sativa* (f). Red lines indicate the 90-percentile distance between the corresponding PRR gene family members and the next closest NB-ARC genes.

## Notes

### Competing Interest Statement

The authors have declared no competing interest.

