## Supplementary figure 1-9 for "Concerted expansion and contraction of immune receptor gene repertoires in plant genomes"

### Slide 1
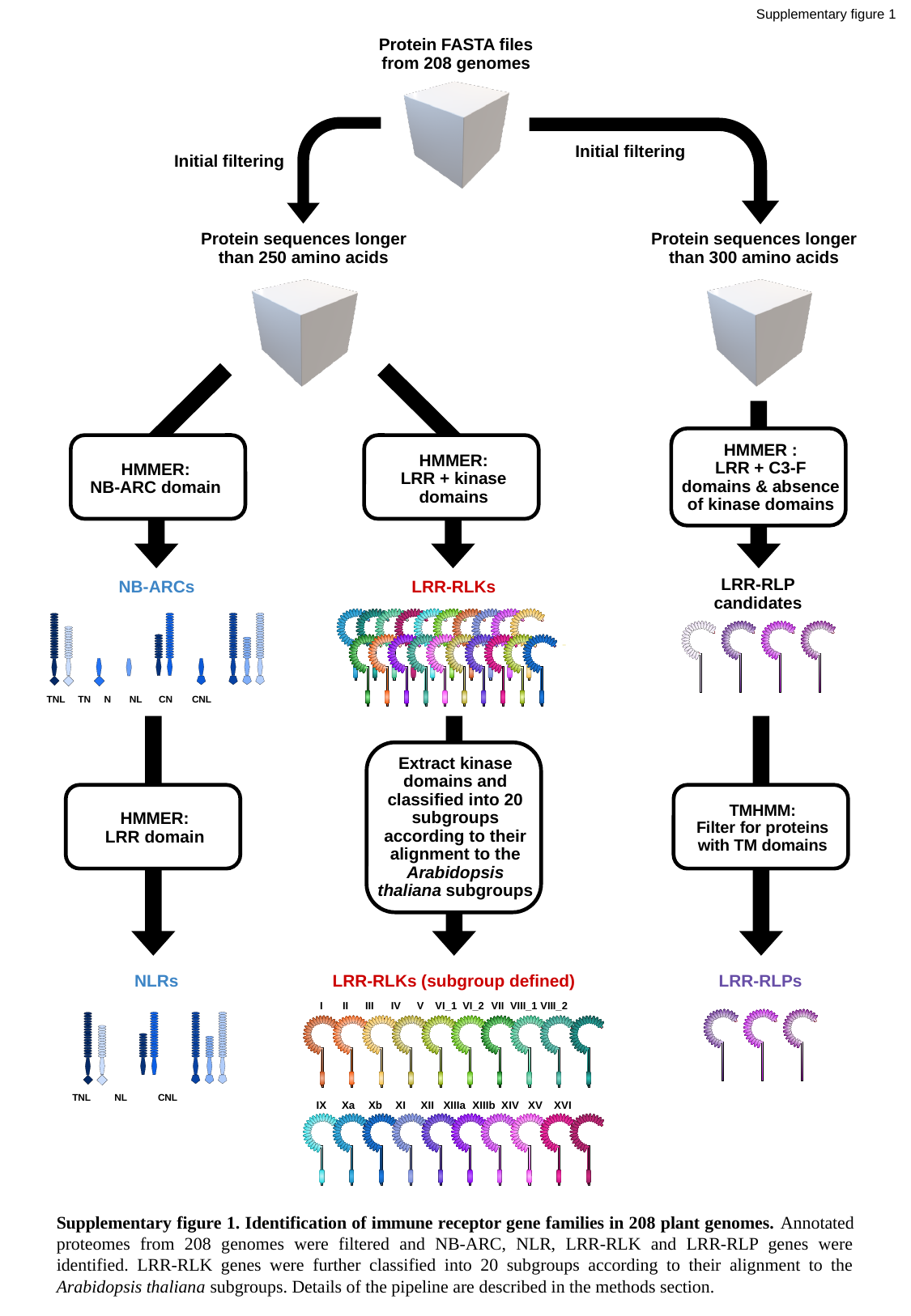

Supplementary figure 1
Protein FASTA files from 208 genomes
Initial filtering
Initial filtering
Protein sequences longer than 250 amino acids
Protein sequences longer than 300 amino acids
HMMER :
LRR + C3-F domains & absence of kinase domains
HMMER:
NB-ARC domain
HMMER:
LRR + kinase domains
NB-ARCs
LRR-RLKs
LRR-RLP candidates
 TNL TN N NL CN CNL
HMMER:
LRR domain
Extract kinase domains and classified into 20 subgroups according to their alignment to the Arabidopsis thaliana subgroups
TMHMM:
Filter for proteins with TM domains
NLRs
LRR-RLKs (subgroup defined)
LRR-RLPs
I II III IV V VI_1 VI_2 VII VIII_1 VIII_2
 TNL NL CNL
IX Xa Xb XI XII XIIIa XIIIb XIV XV XVI
Supplementary figure 1. Identification of immune receptor gene families in 208 plant genomes. Annotated proteomes from 208 genomes were filtered and NB-ARC, NLR, LRR-RLK and LRR-RLP genes were identified. LRR-RLK genes were further classified into 20 subgroups according to their alignment to the Arabidopsis thaliana subgroups. Details of the pipeline are described in the methods section.

### Slide 2
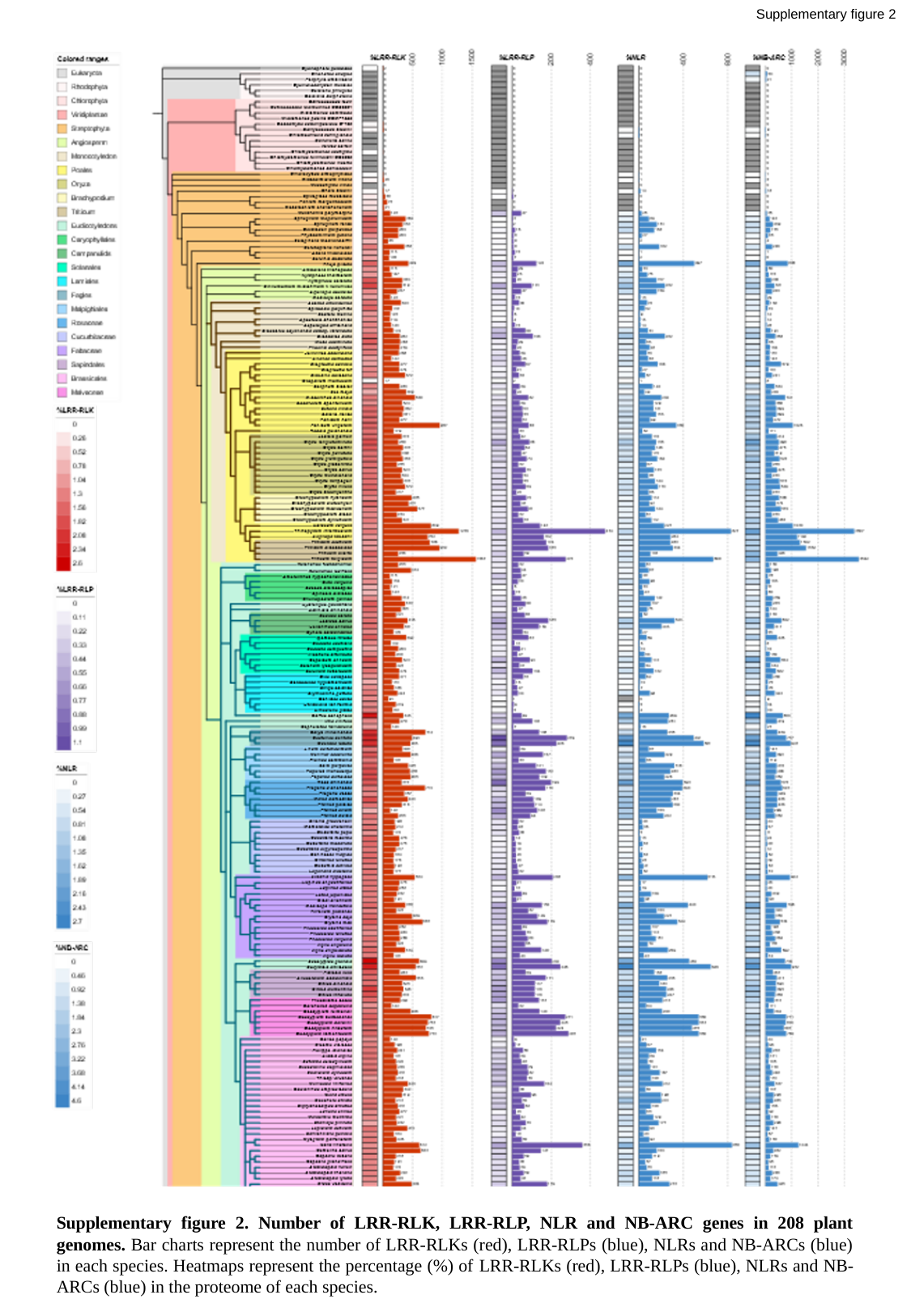

Supplementary figure 2
Supplementary figure 2. Number of LRR-RLK, LRR-RLP, NLR and NB-ARC genes in 208 plant genomes. Bar charts represent the number of LRR-RLKs (red), LRR-RLPs (blue), NLRs and NB-ARCs (blue) in each species. Heatmaps represent the percentage (%) of LRR-RLKs (red), LRR-RLPs (blue), NLRs and NB-ARCs (blue) in the proteome of each species.

### Slide 3
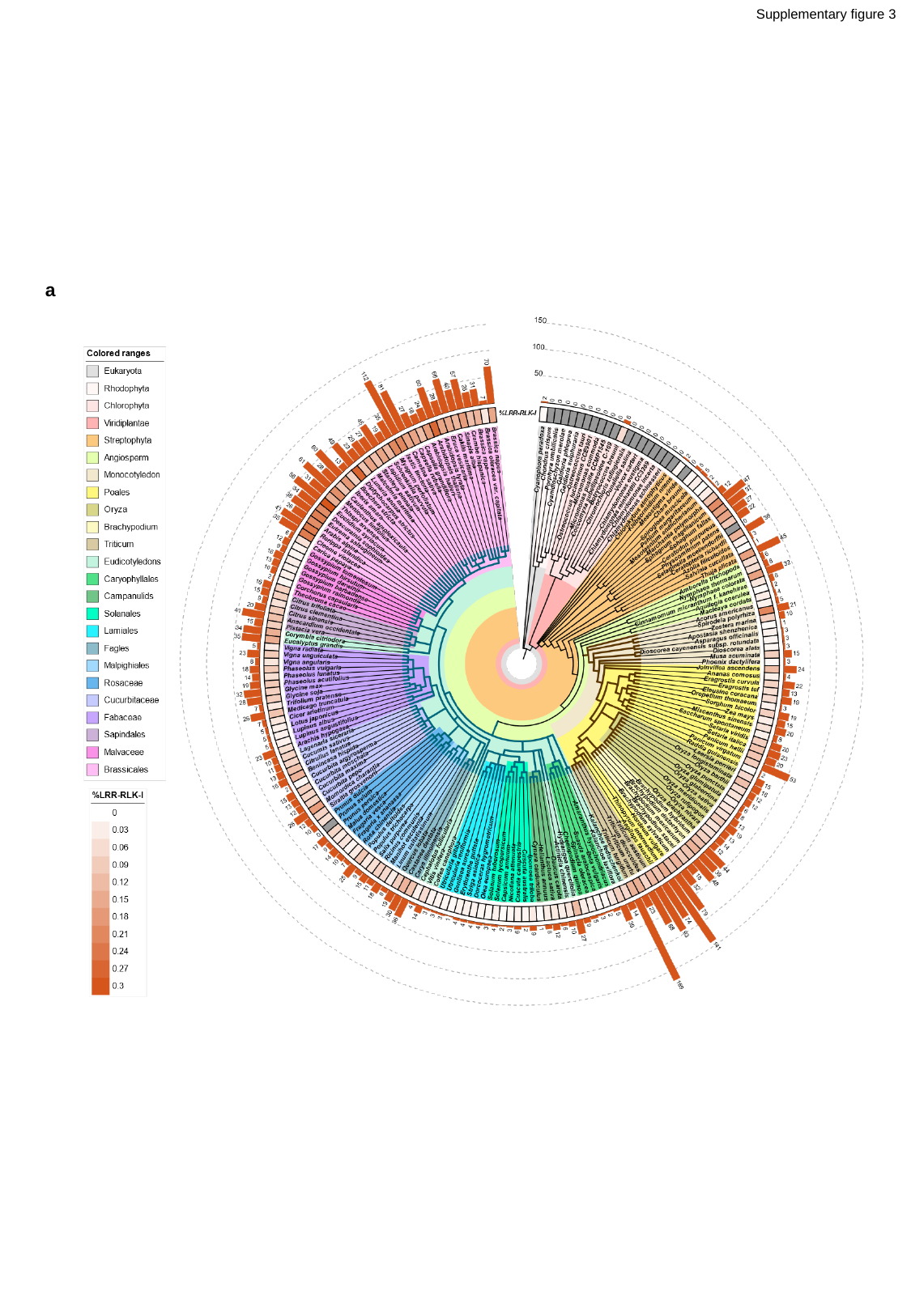

Supplementary figure 3
a

### Slide 4
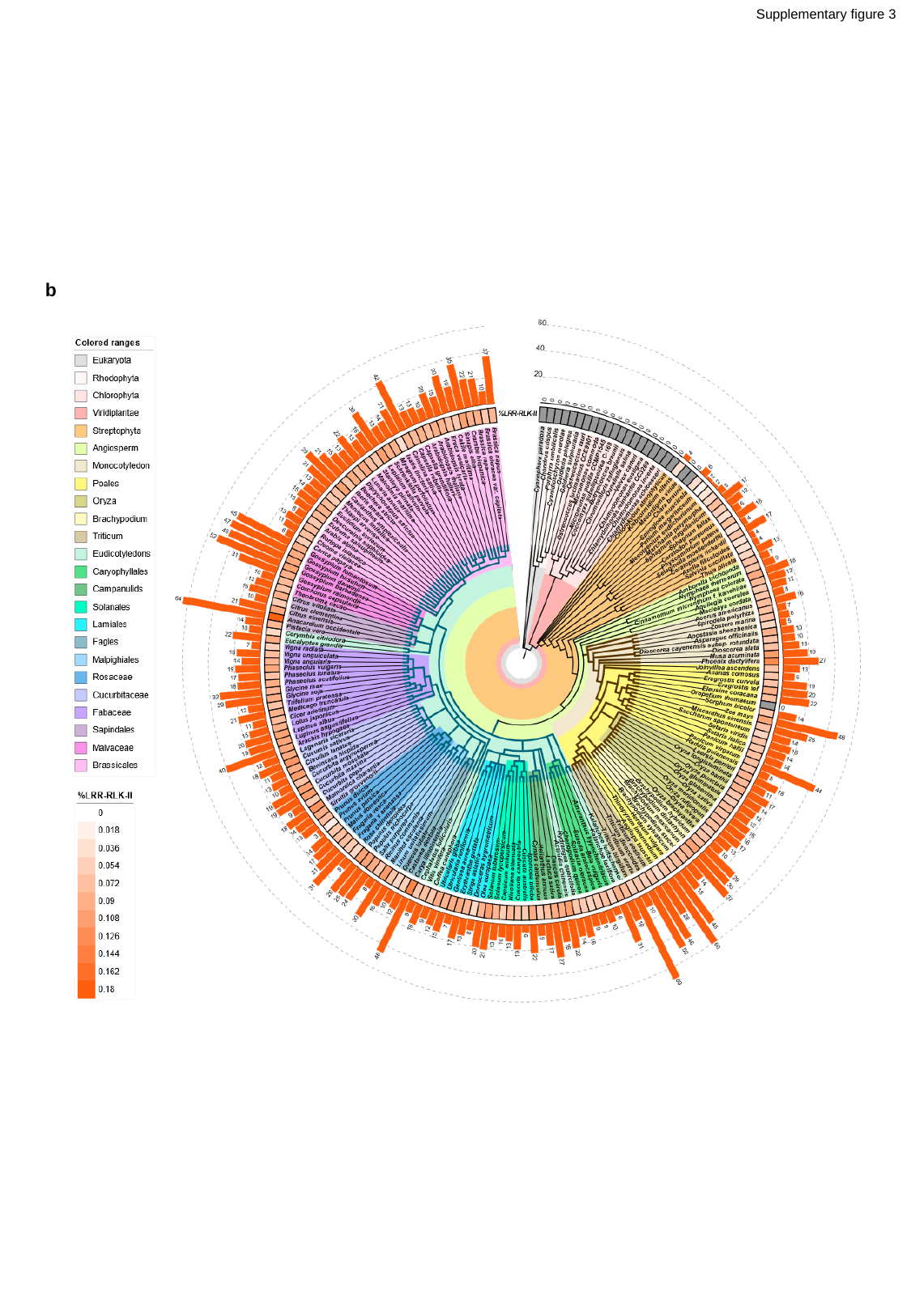

Supplementary figure 3
b

### Slide 5
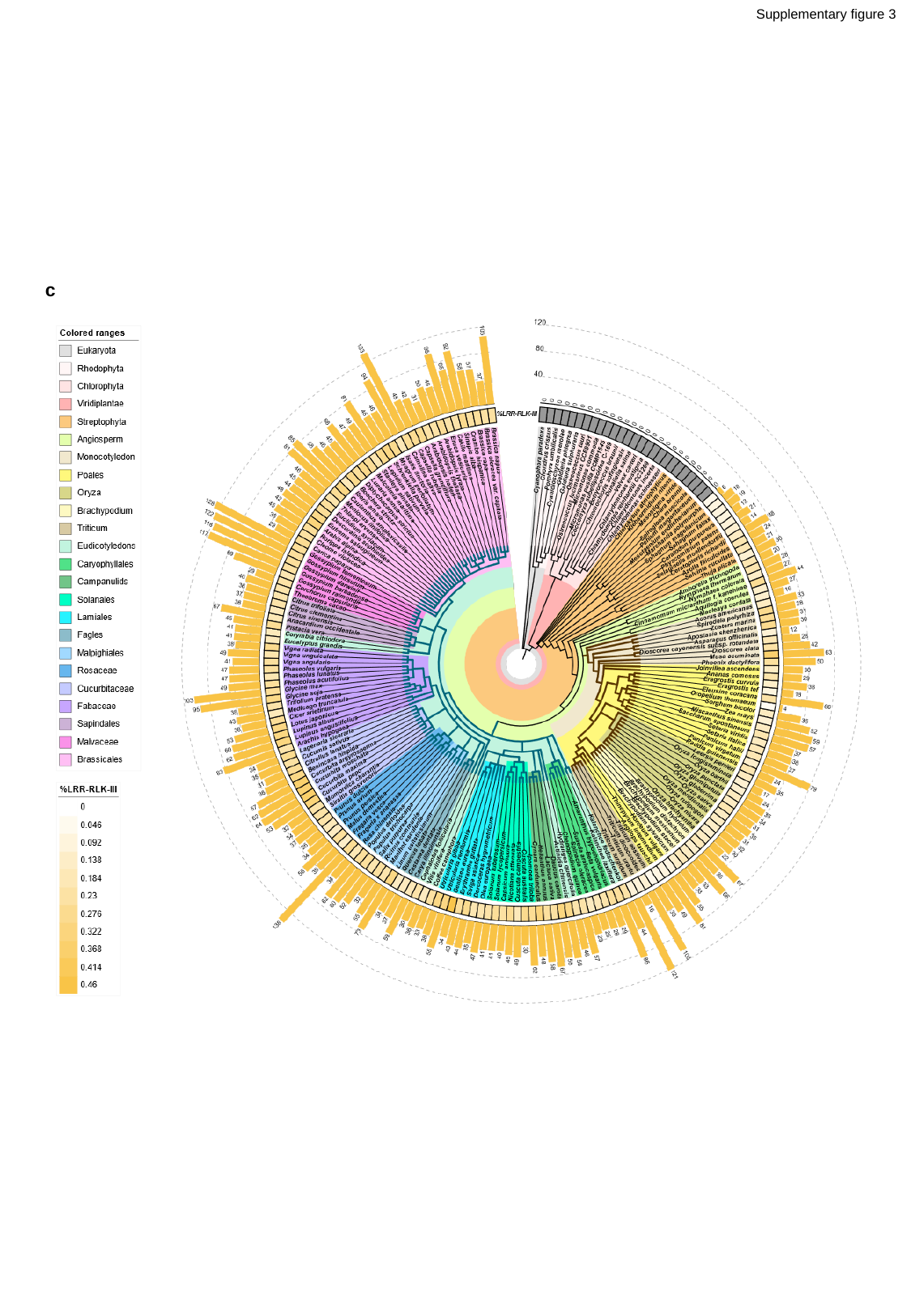

Supplementary figure 3
c

### Slide 6
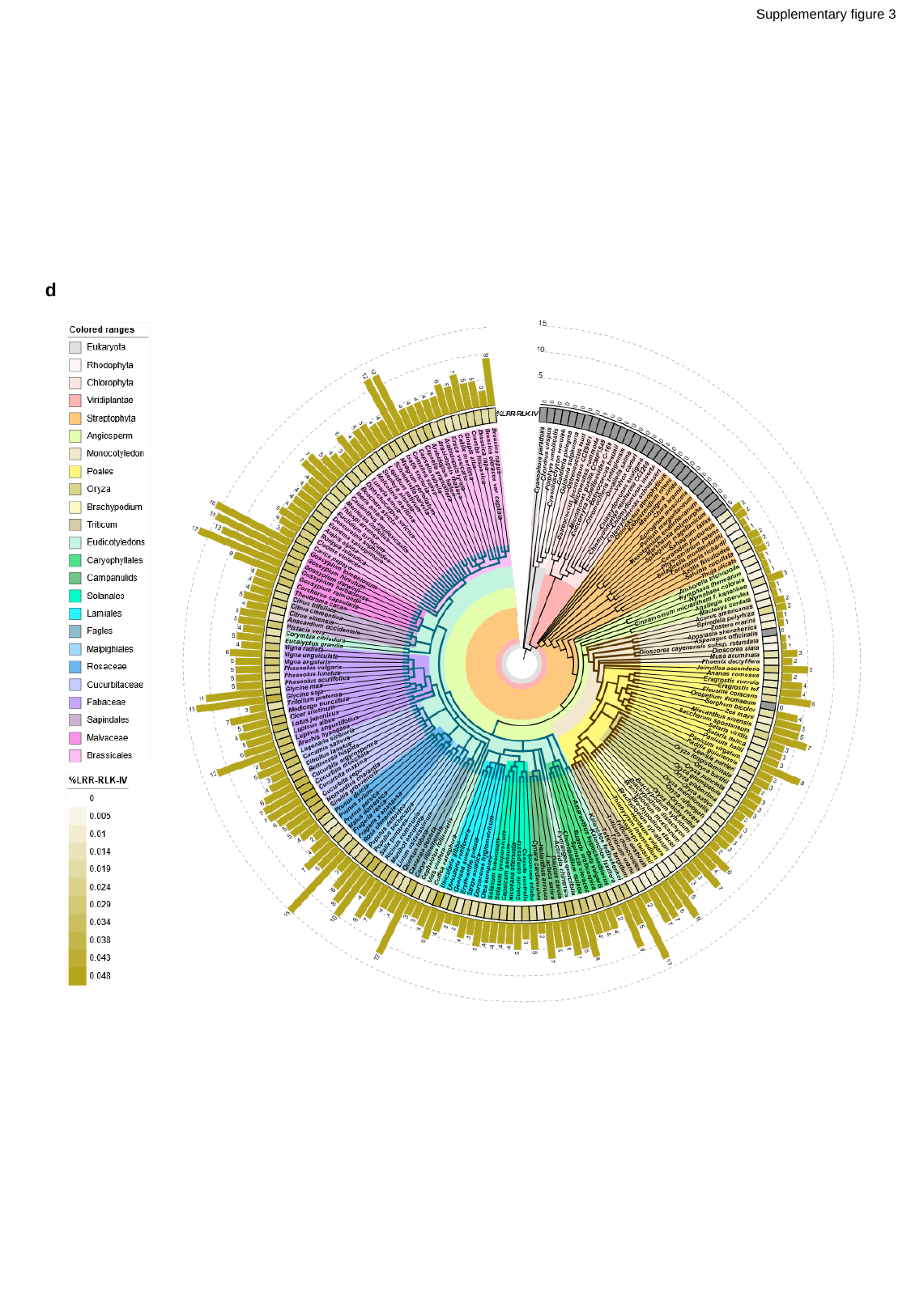

Supplementary figure 3
d

### Slide 7
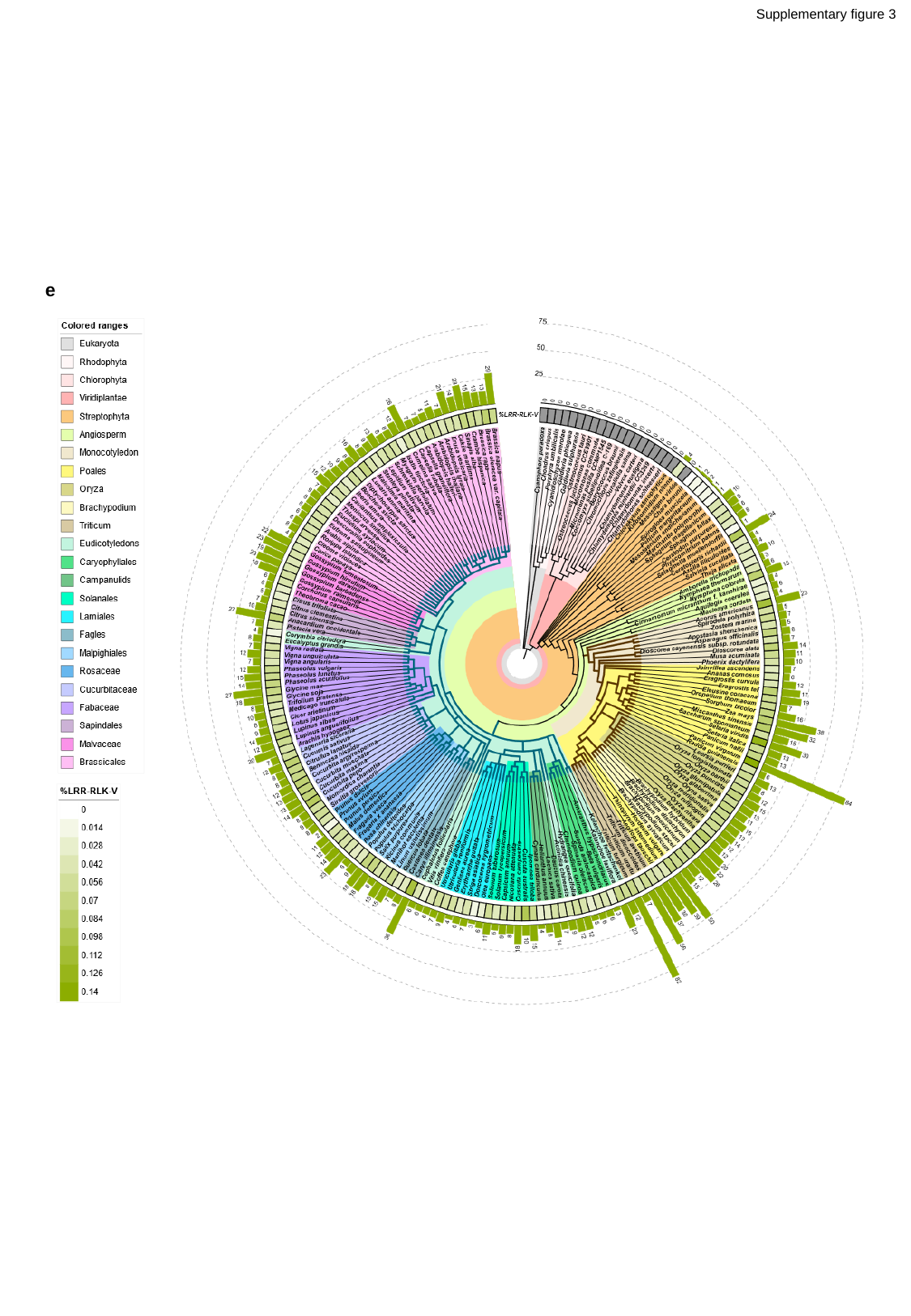

Supplementary figure 3
e

### Slide 8
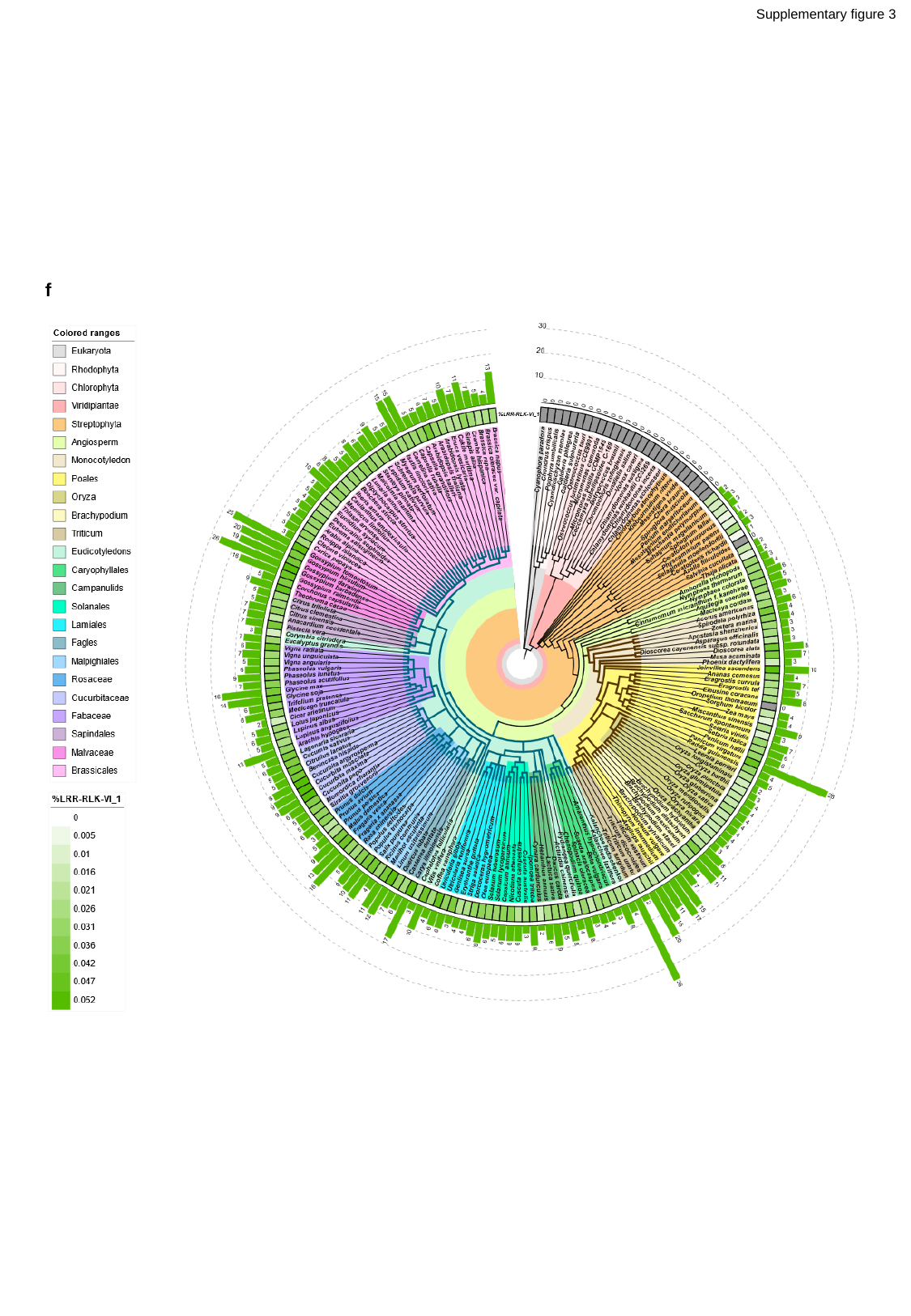

Supplementary figure 3
f

### Slide 9
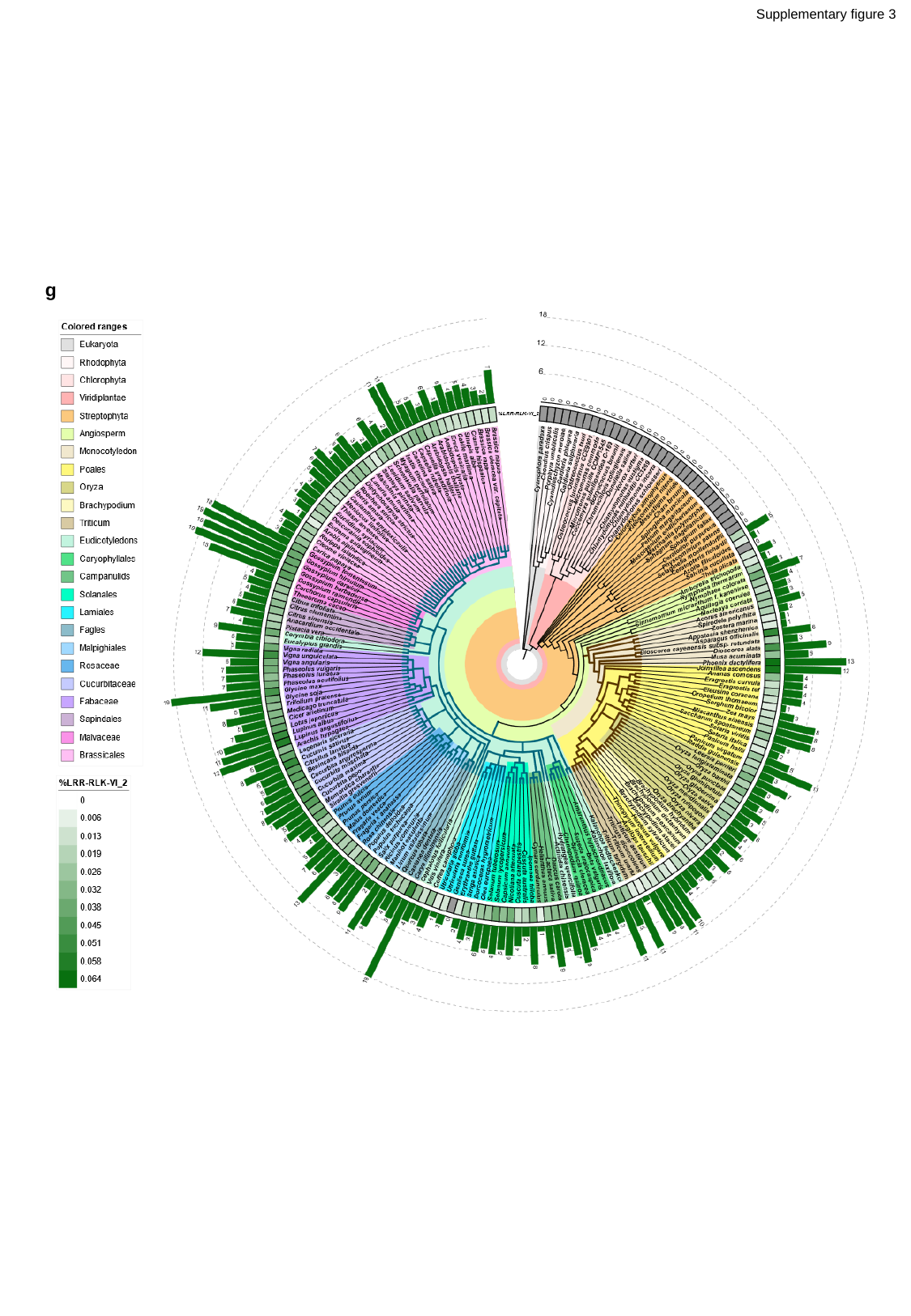

Supplementary figure 3
g

### Slide 10
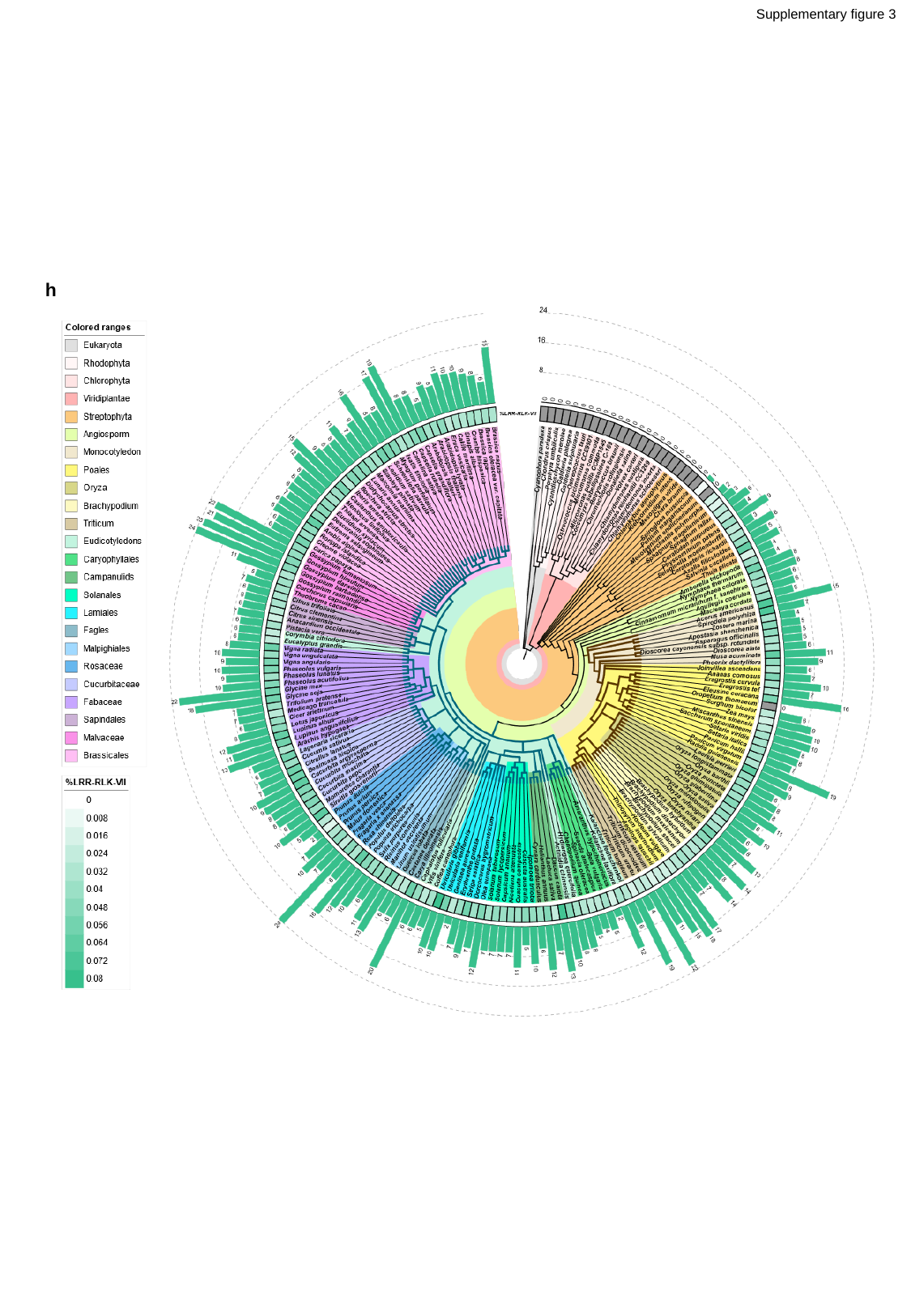

Supplementary figure 3
h

### Slide 11
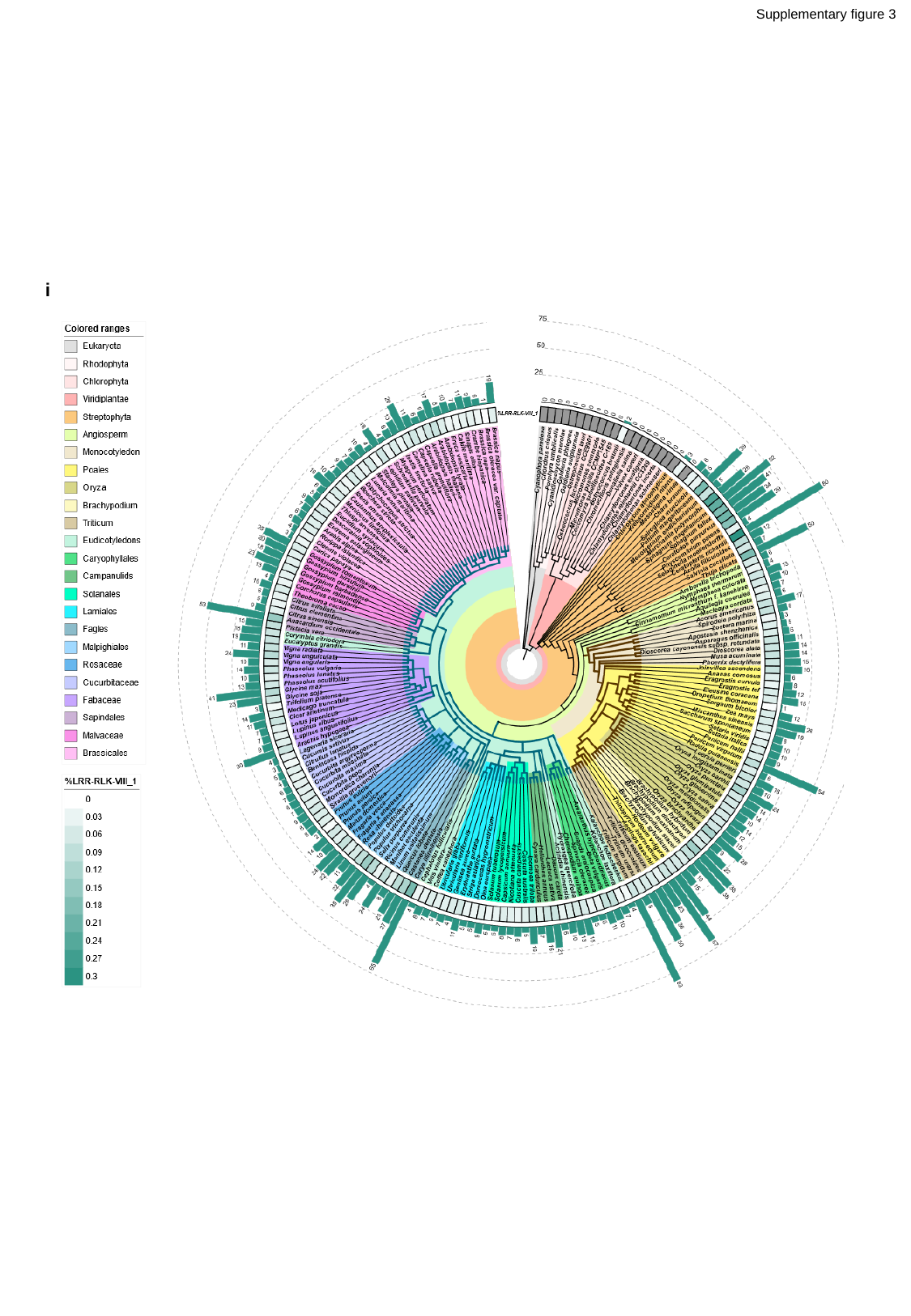

Supplementary figure 3
i

### Slide 12
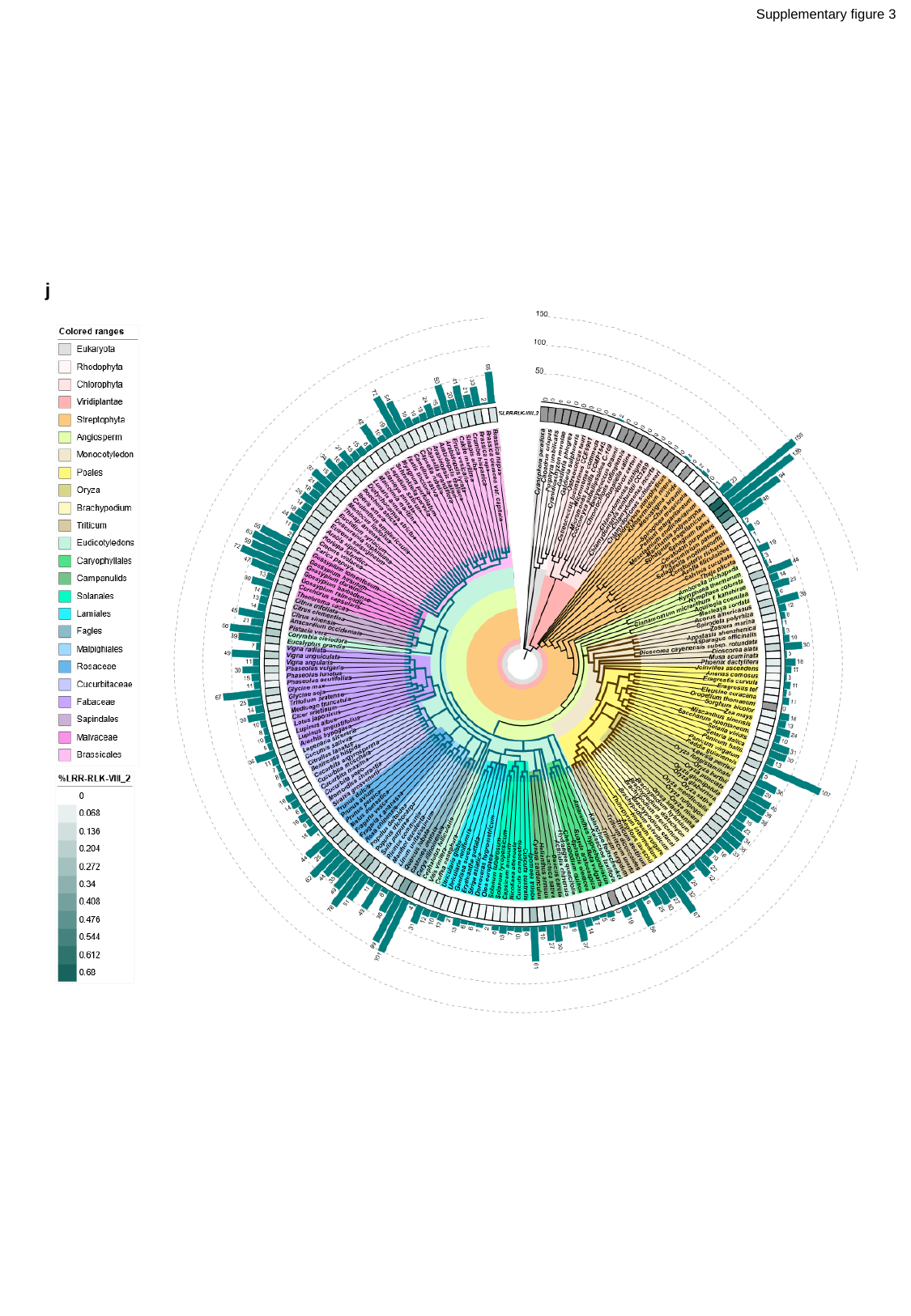

Supplementary figure 3
j

### Slide 13
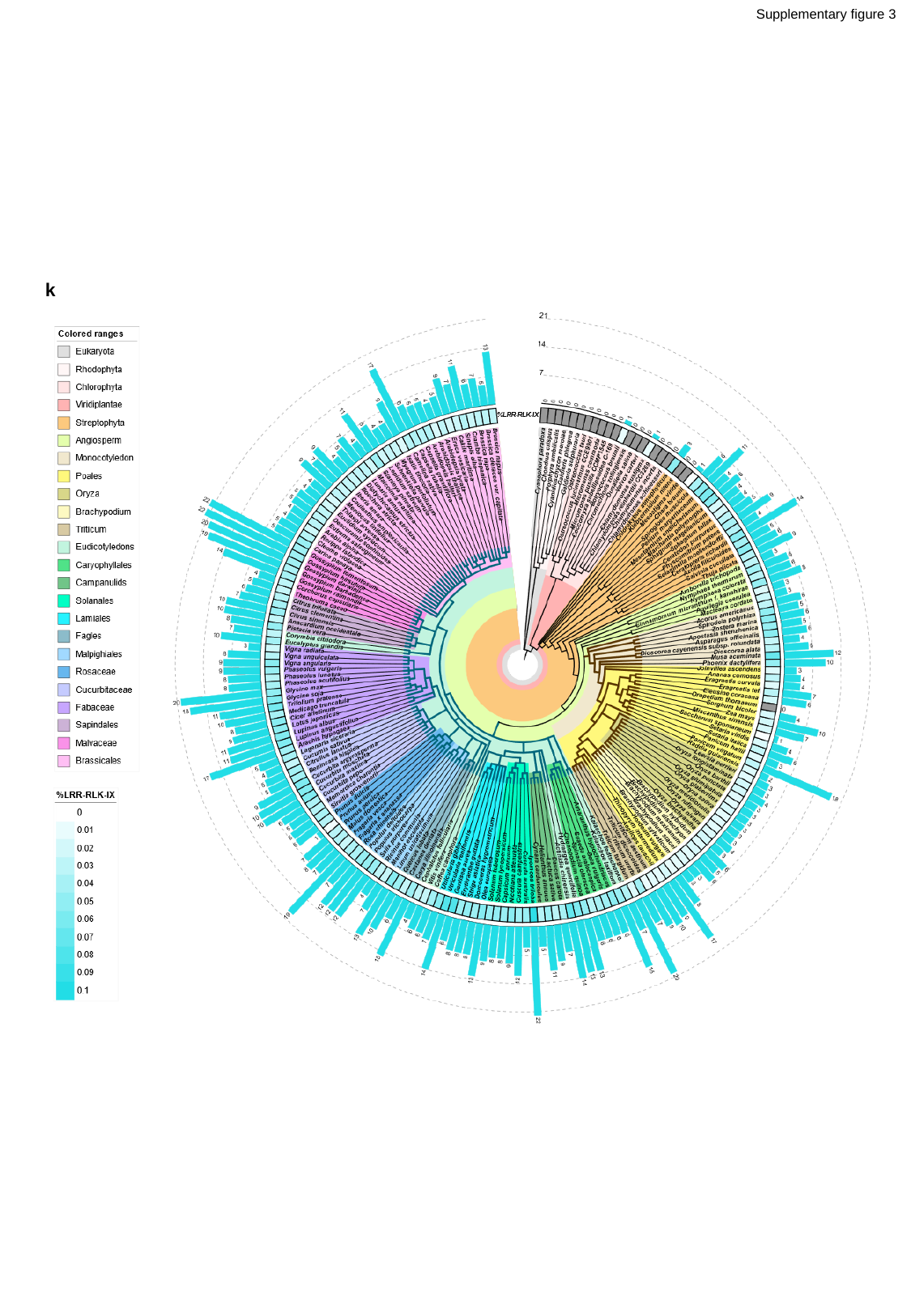

Supplementary figure 3
k

### Slide 14
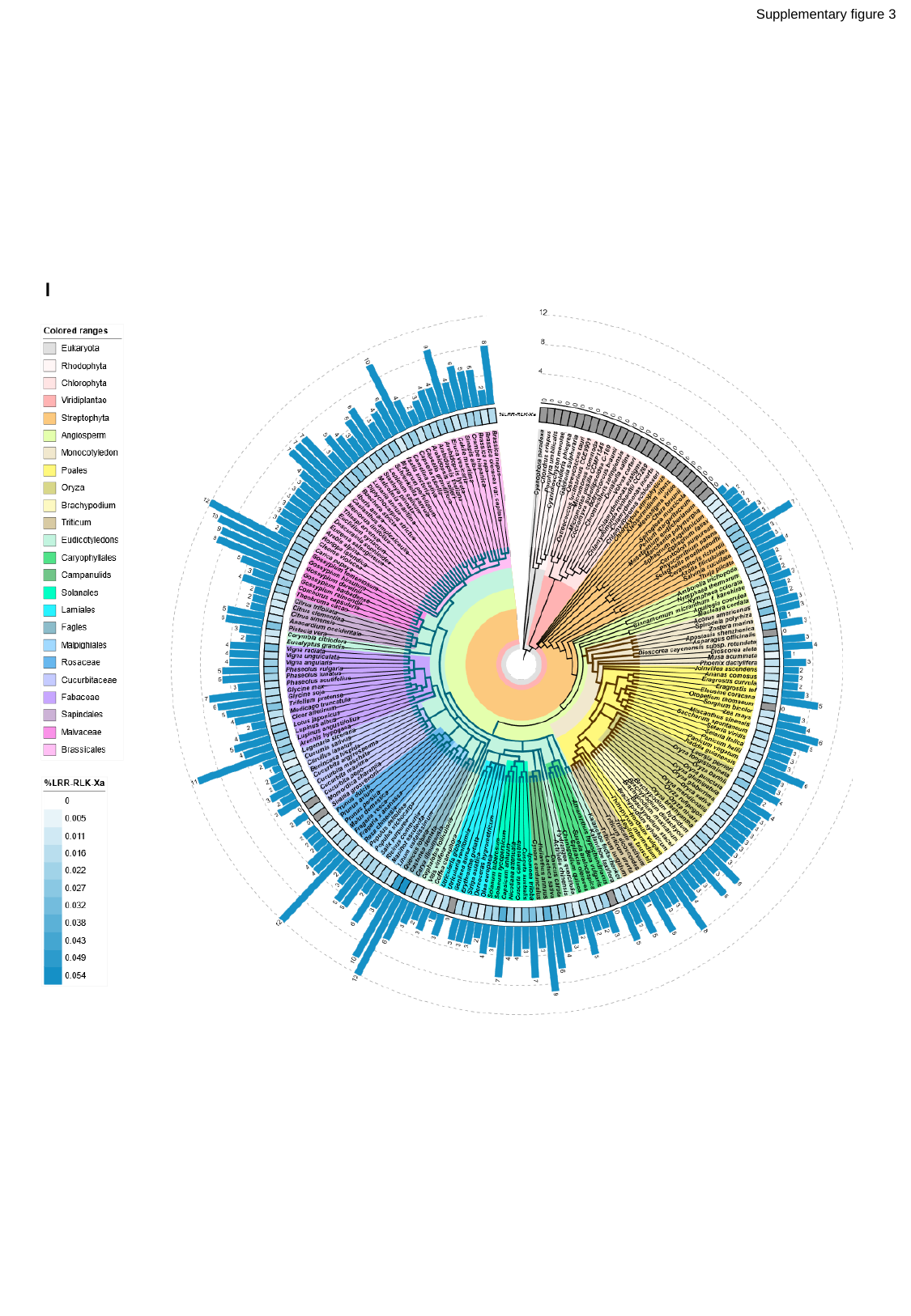

Supplementary figure 3
l

### Slide 15
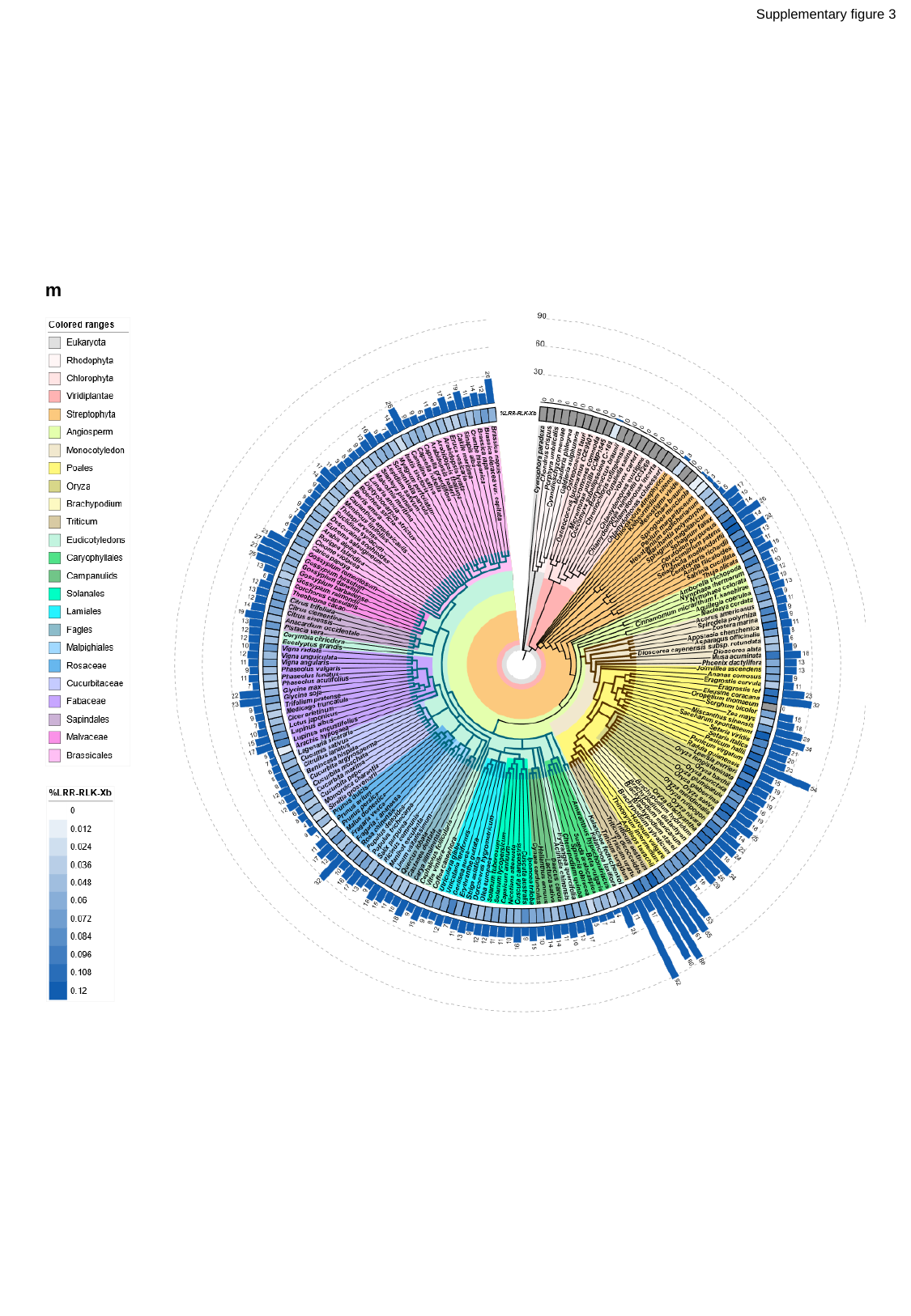

Supplementary figure 3
m

### Slide 16
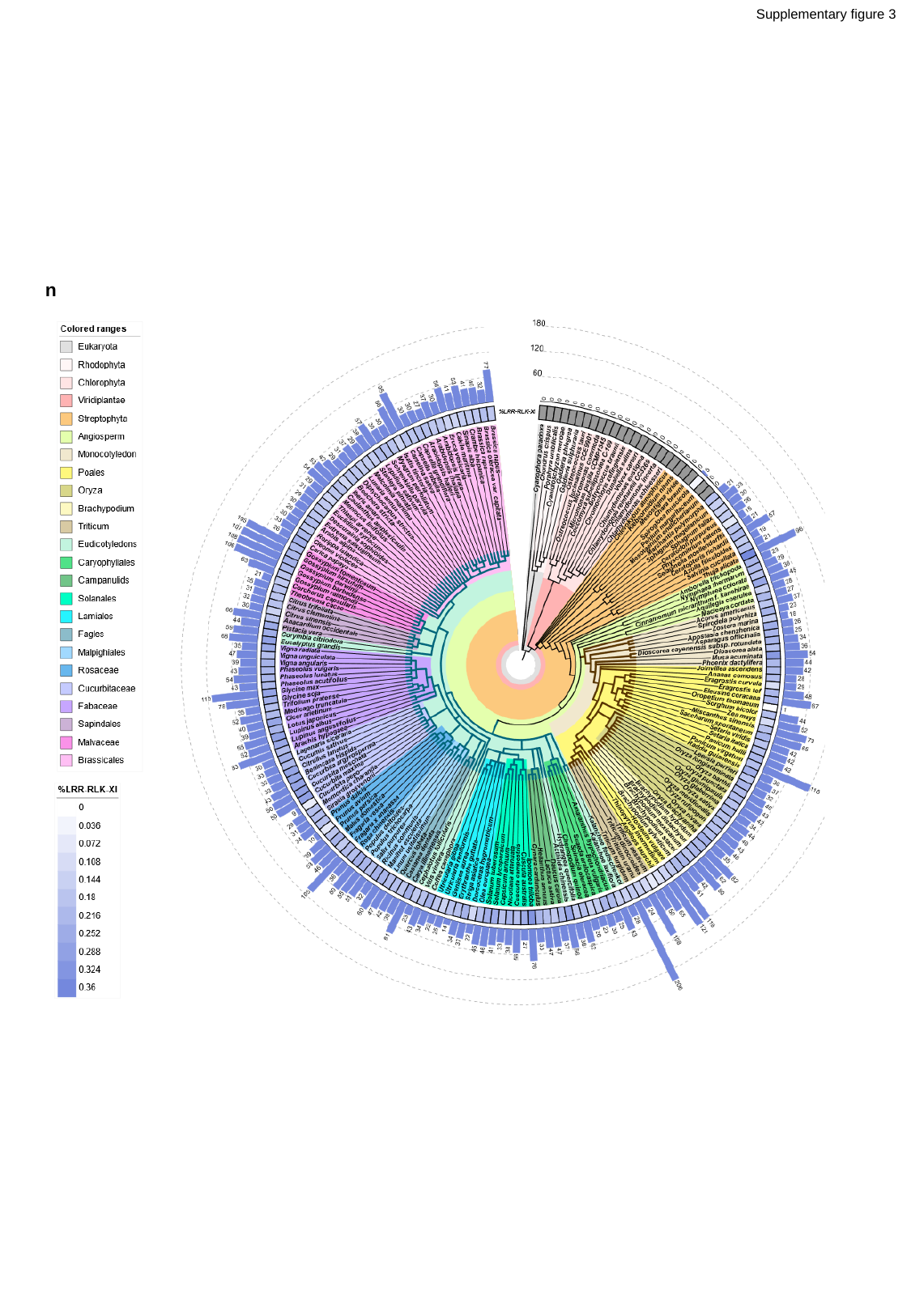

Supplementary figure 3
n

### Slide 17
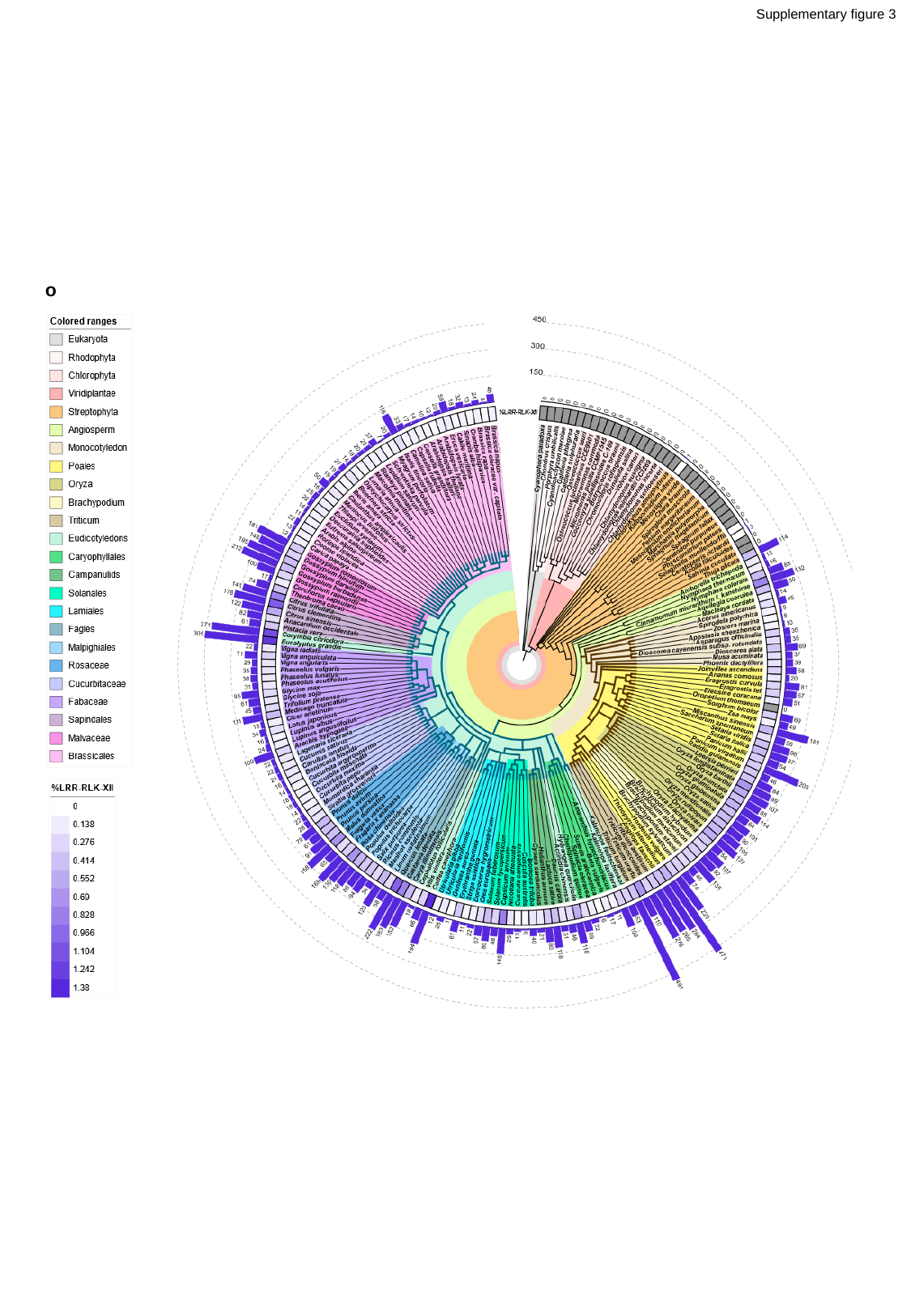

Supplementary figure 3
o

### Slide 18
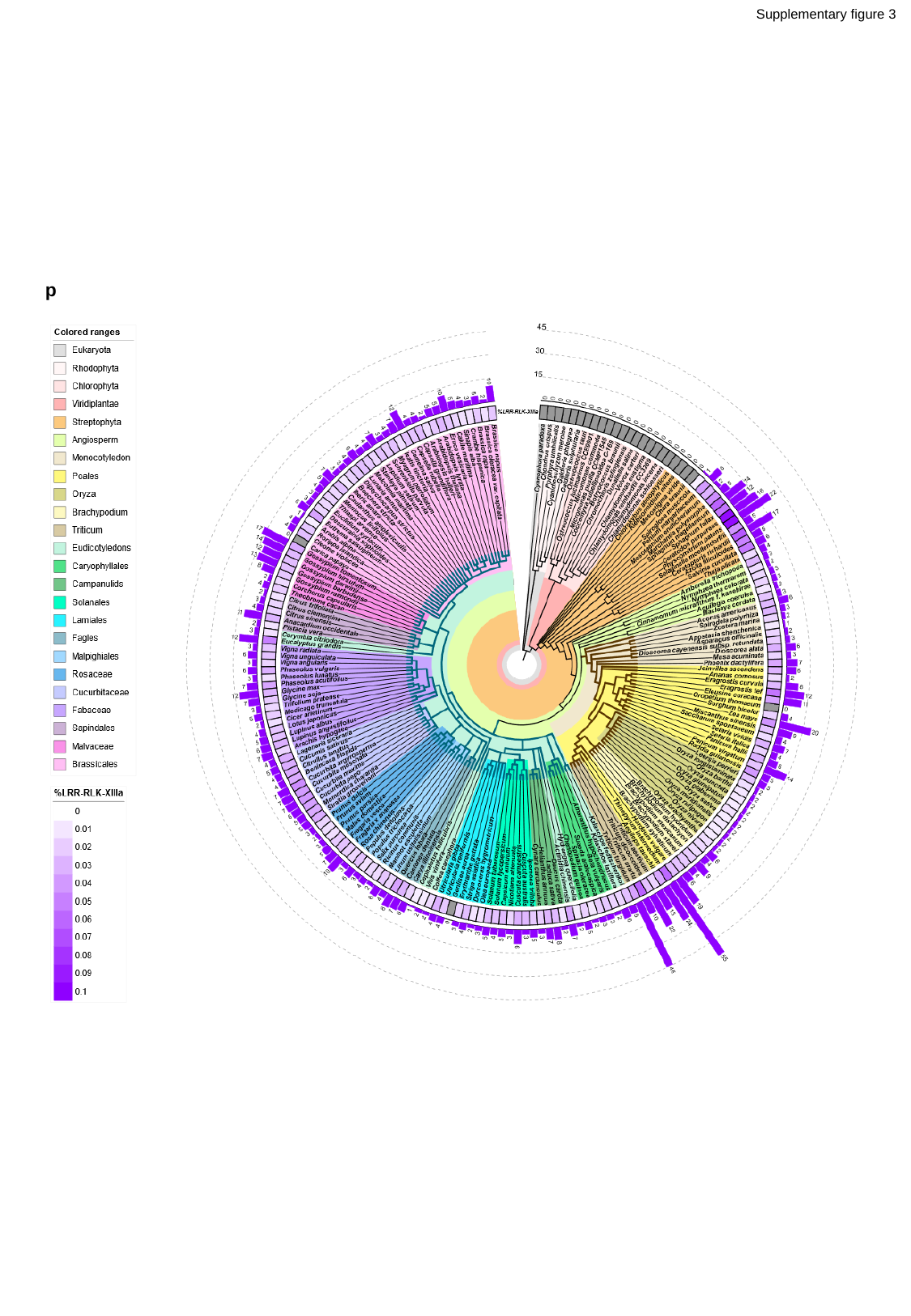

Supplementary figure 3
p

### Slide 19
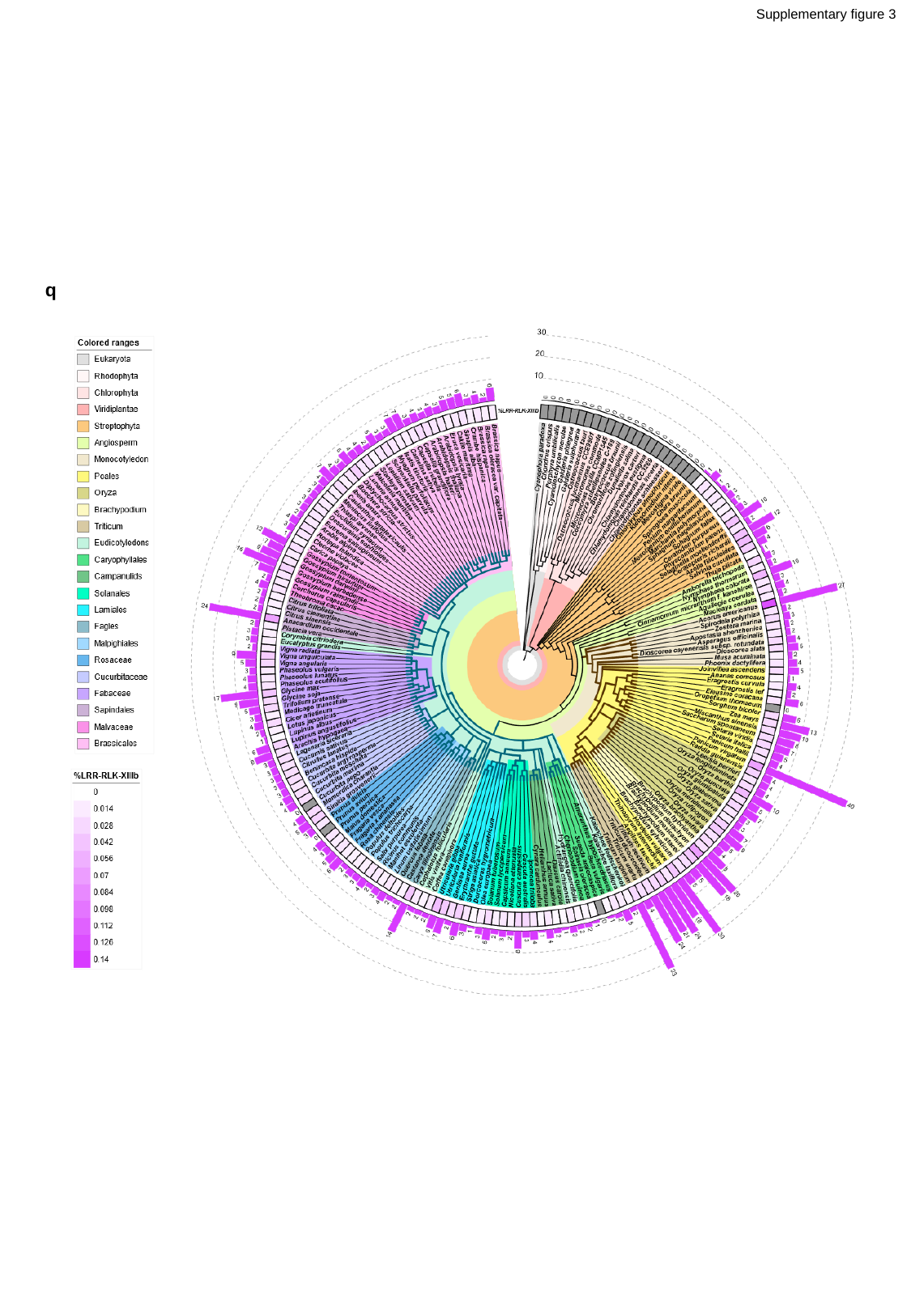

Supplementary figure 3
q

### Slide 20
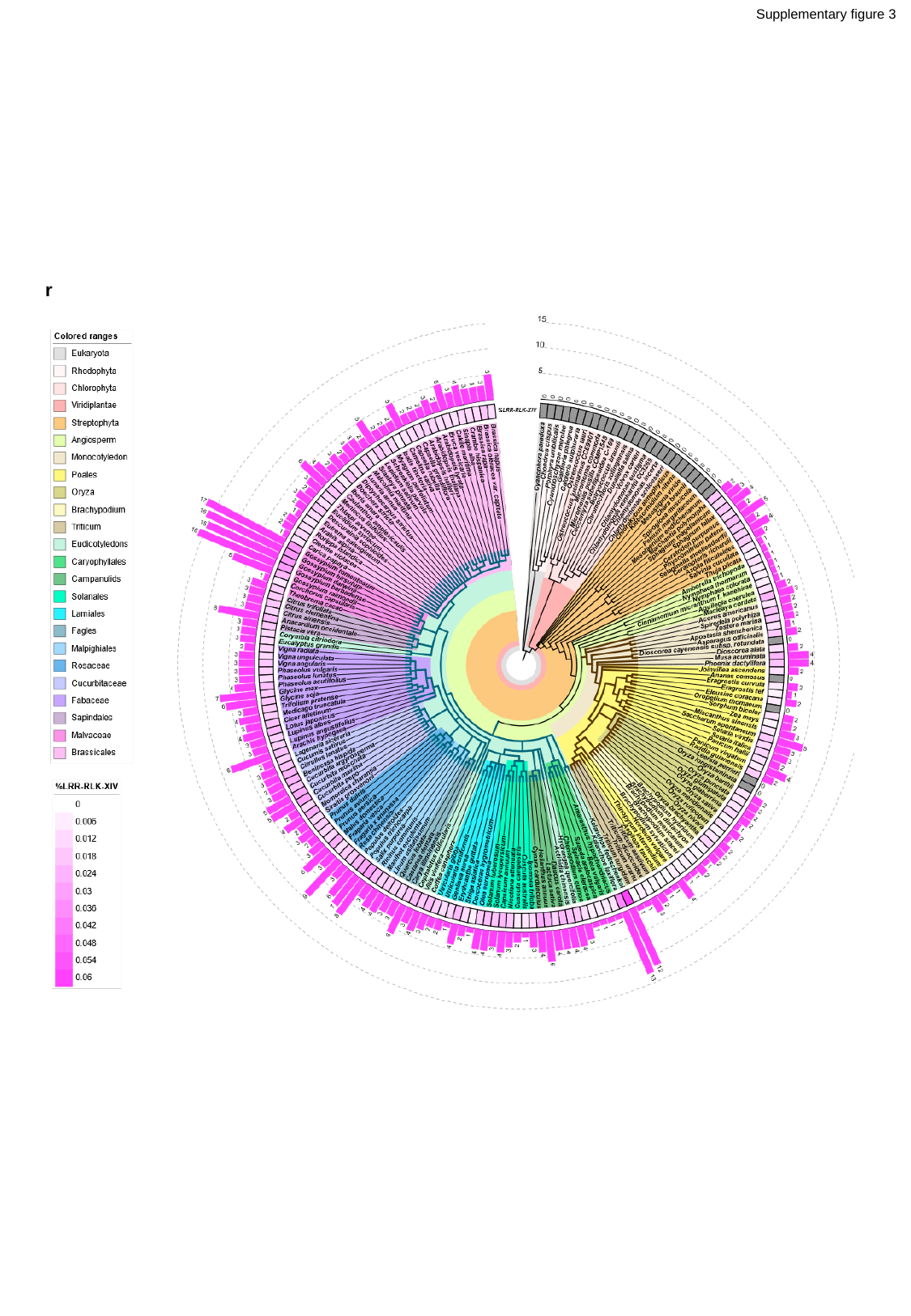

Supplementary figure 3
r

### Slide 21
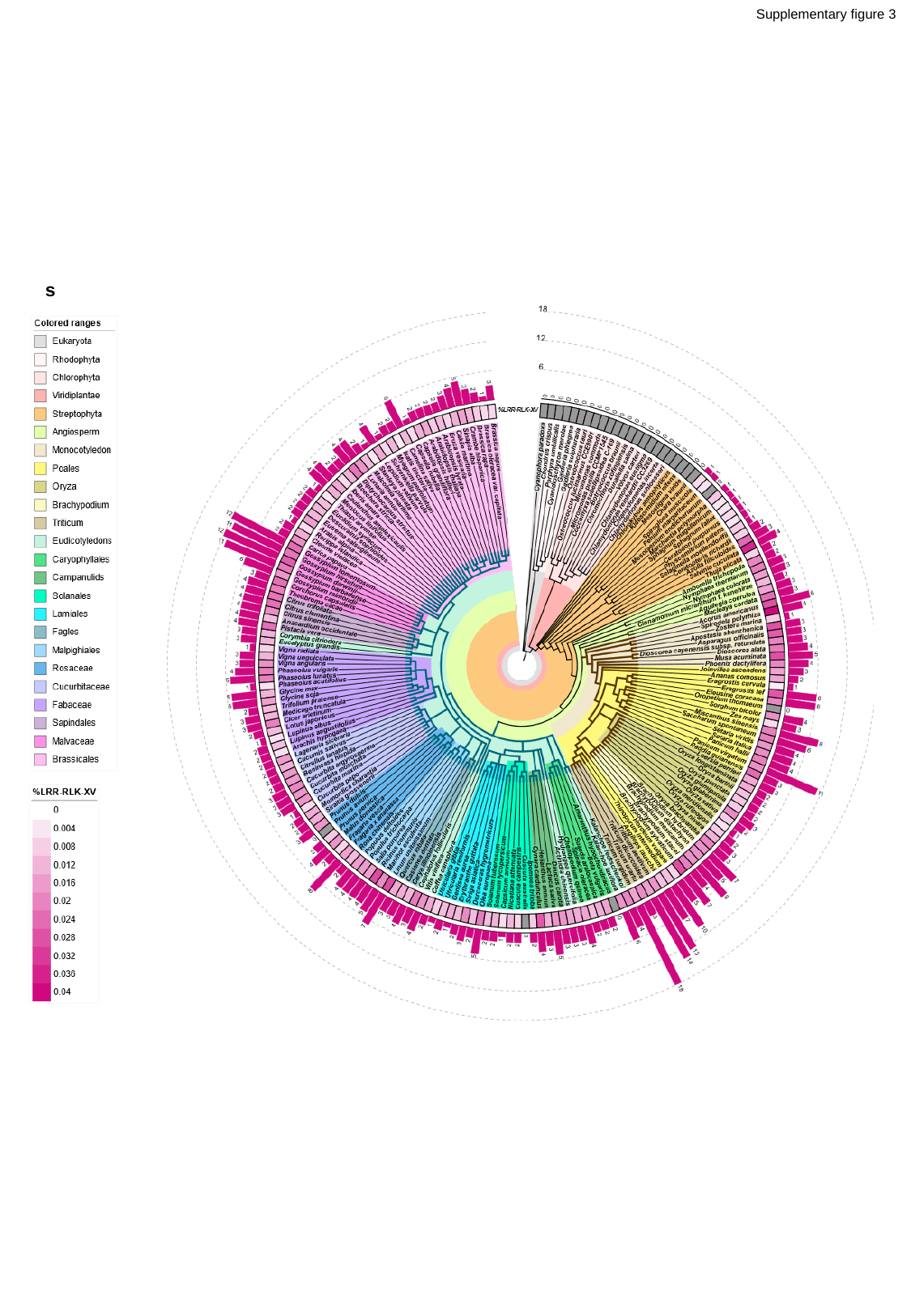

Supplementary figure 3
s

### Slide 22
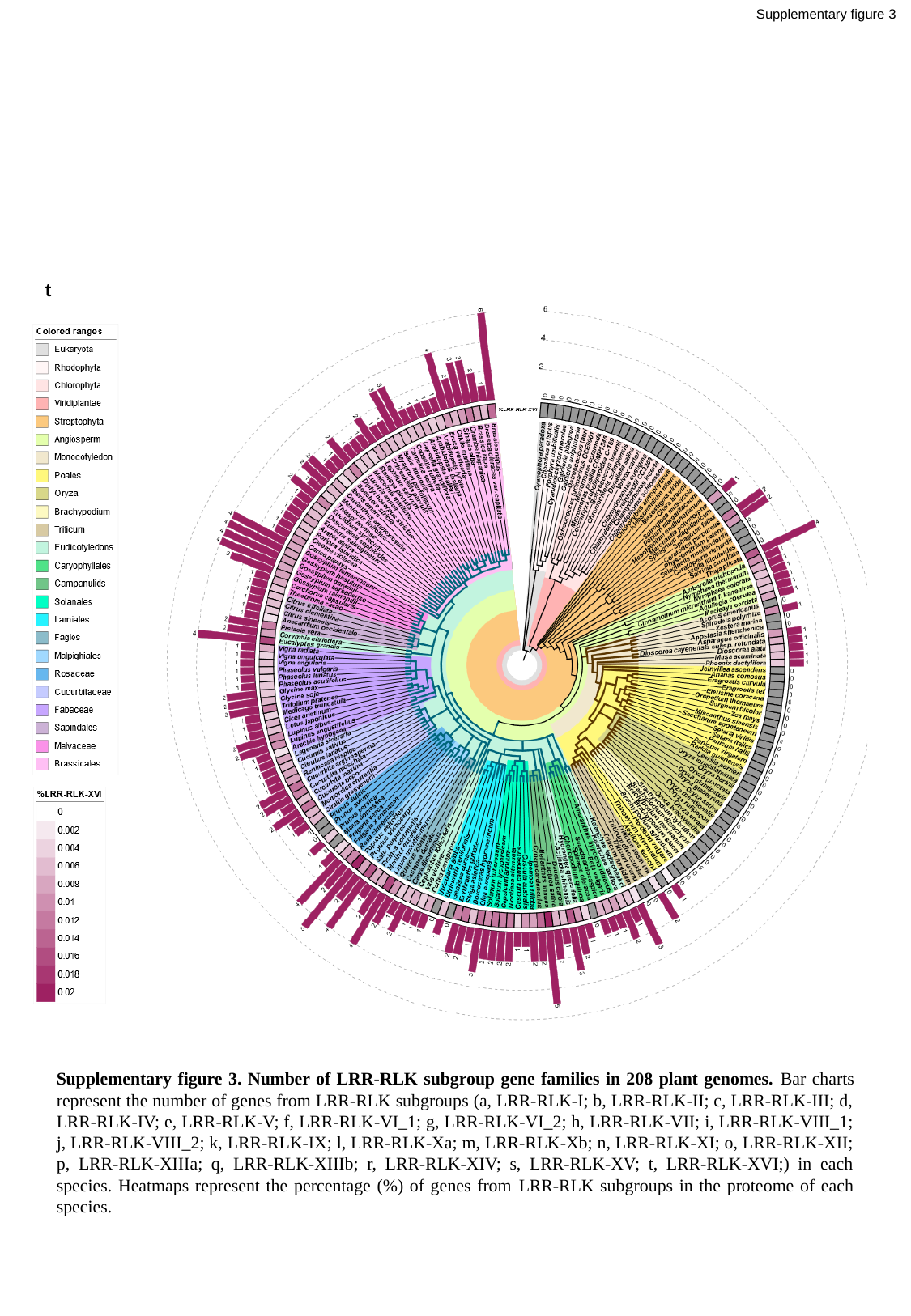

Supplementary figure 3
t
Supplementary figure 3. Number of LRR-RLK subgroup gene families in 208 plant genomes. Bar charts represent the number of genes from LRR-RLK subgroups (a, LRR-RLK-I; b, LRR-RLK-II; c, LRR-RLK-III; d, LRR-RLK-IV; e, LRR-RLK-V; f, LRR-RLK-VI_1; g, LRR-RLK-VI_2; h, LRR-RLK-VII; i, LRR-RLK-VIII_1; j, LRR-RLK-VIII_2; k, LRR-RLK-IX; l, LRR-RLK-Xa; m, LRR-RLK-Xb; n, LRR-RLK-XI; o, LRR-RLK-XII; p, LRR-RLK-XIIIa; q, LRR-RLK-XIIIb; r, LRR-RLK-XIV; s, LRR-RLK-XV; t, LRR-RLK-XVI;) in each species. Heatmaps represent the percentage (%) of genes from LRR-RLK subgroups in the proteome of each species.

### Slide 23
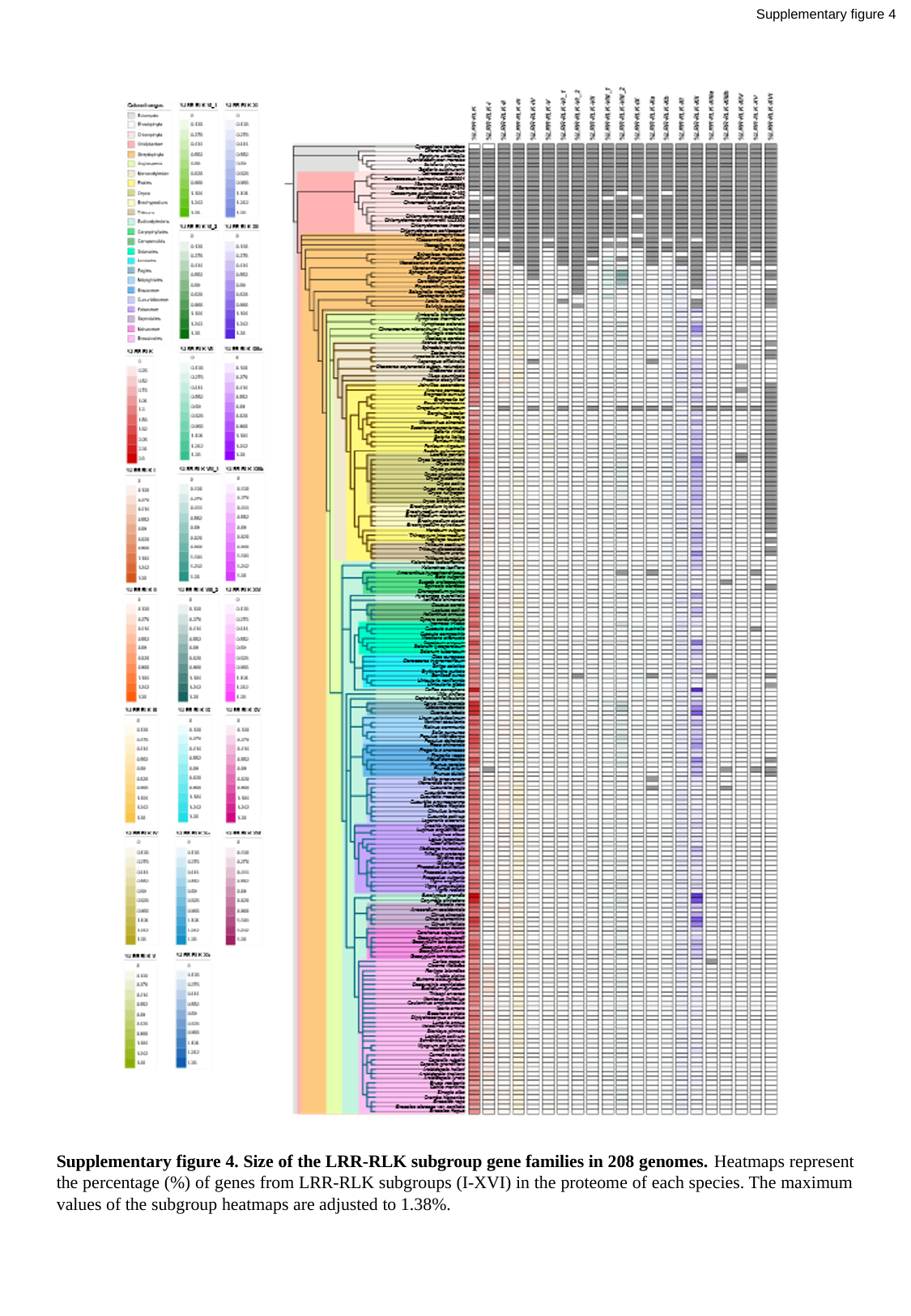

Supplementary figure 4
Supplementary figure 4. Size of the LRR-RLK subgroup gene families in 208 genomes. Heatmaps represent the percentage (%) of genes from LRR-RLK subgroups (I-XVI) in the proteome of each species. The maximum values of the subgroup heatmaps are adjusted to 1.38%.

### Slide 24
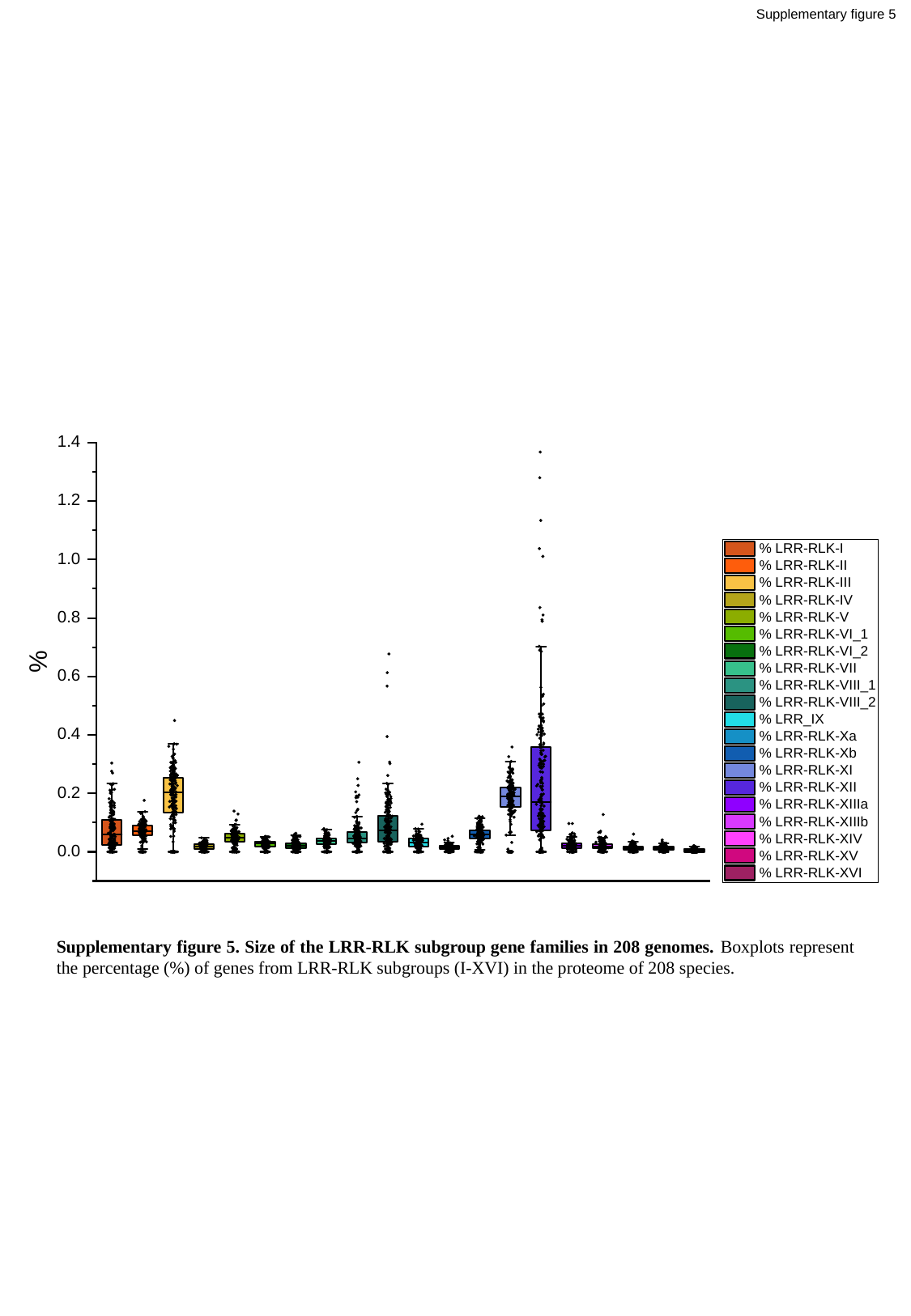

Supplementary figure 5
Supplementary figure 5. Size of the LRR-RLK subgroup gene families in 208 genomes. Boxplots represent the percentage (%) of genes from LRR-RLK subgroups (I-XVI) in the proteome of 208 species.

### Slide 25
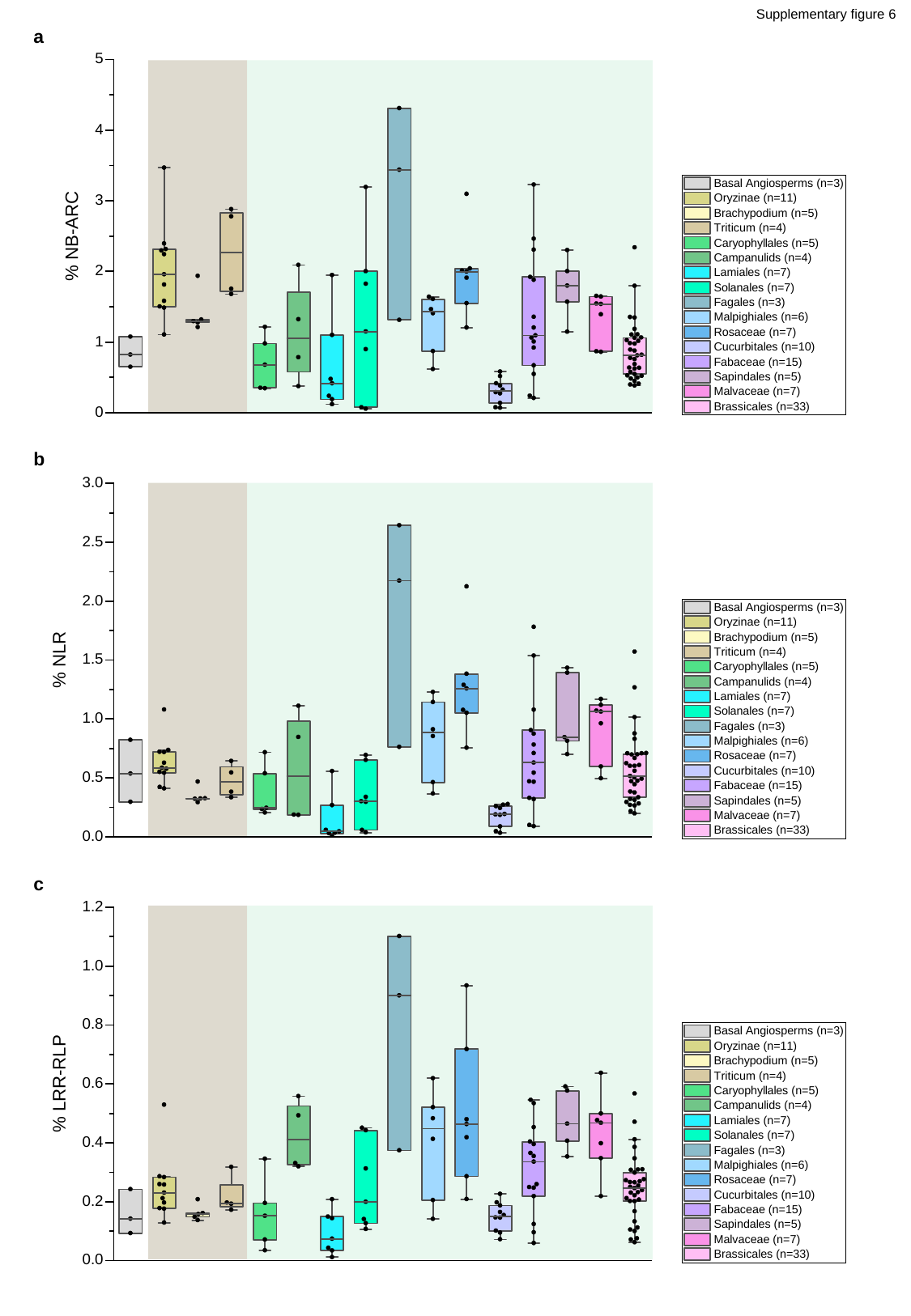

Supplementary figure 6
a
b
c

### Slide 26
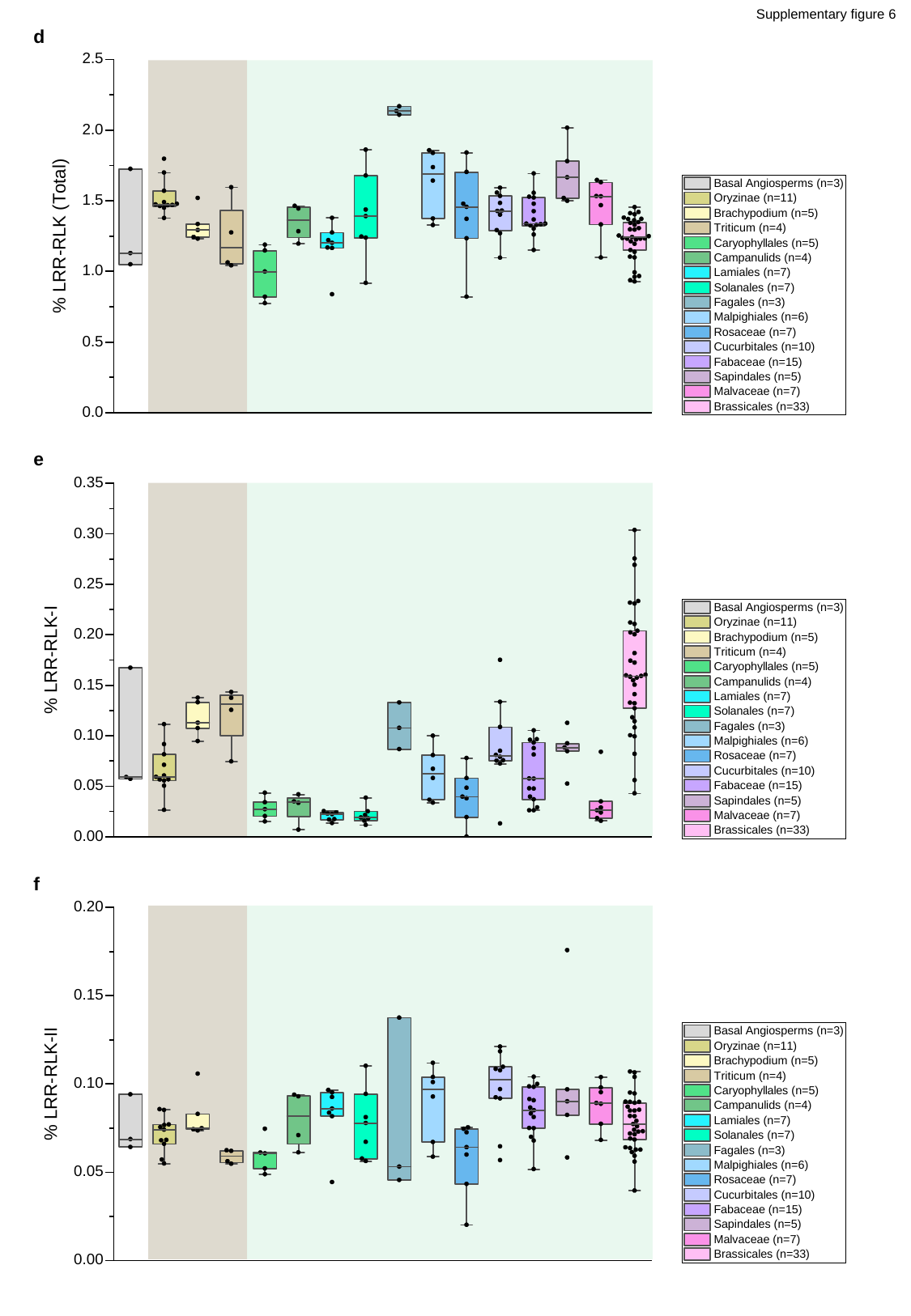

Supplementary figure 6
d
e
f

### Slide 27
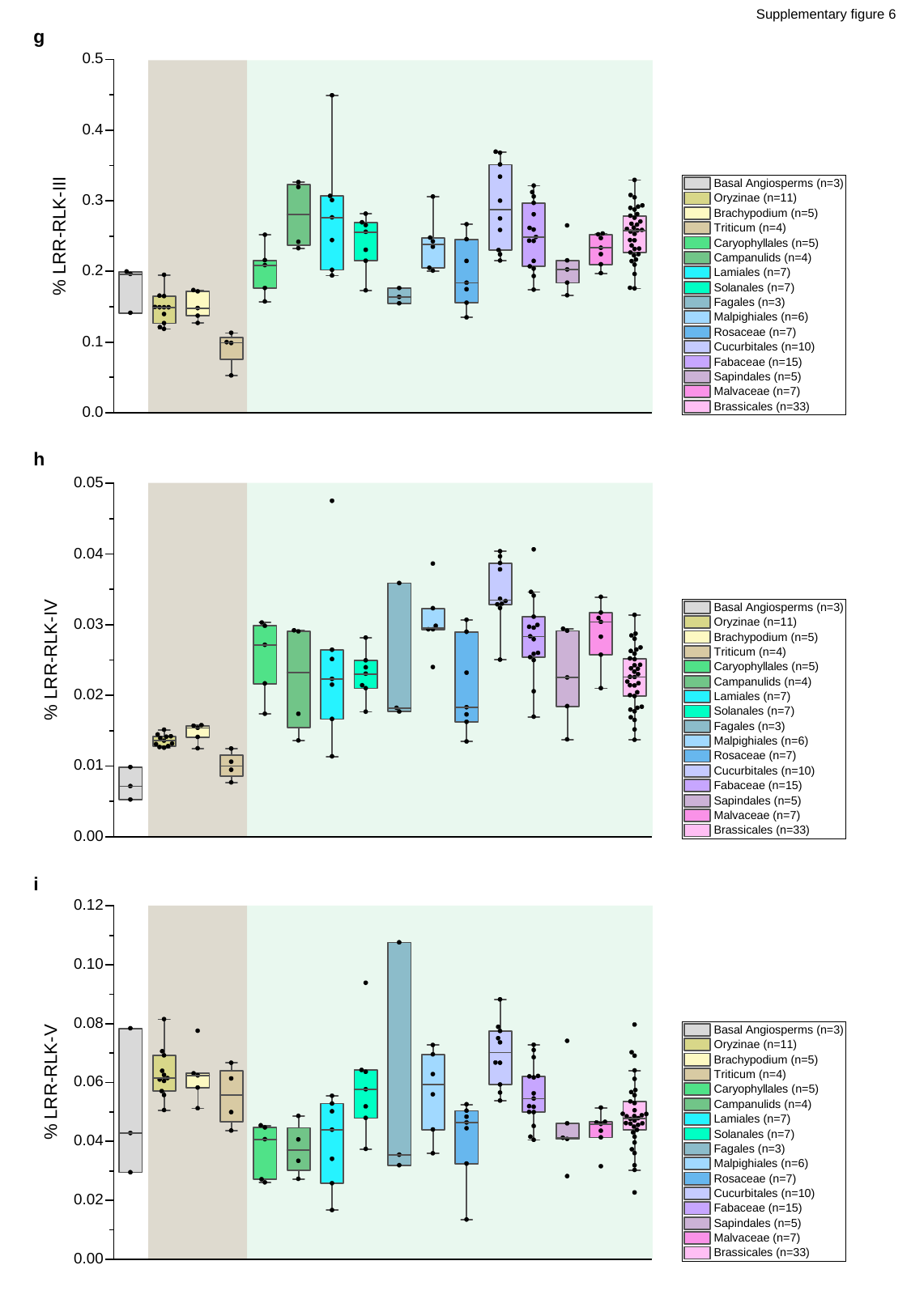

Supplementary figure 6
g
h
i

### Slide 28
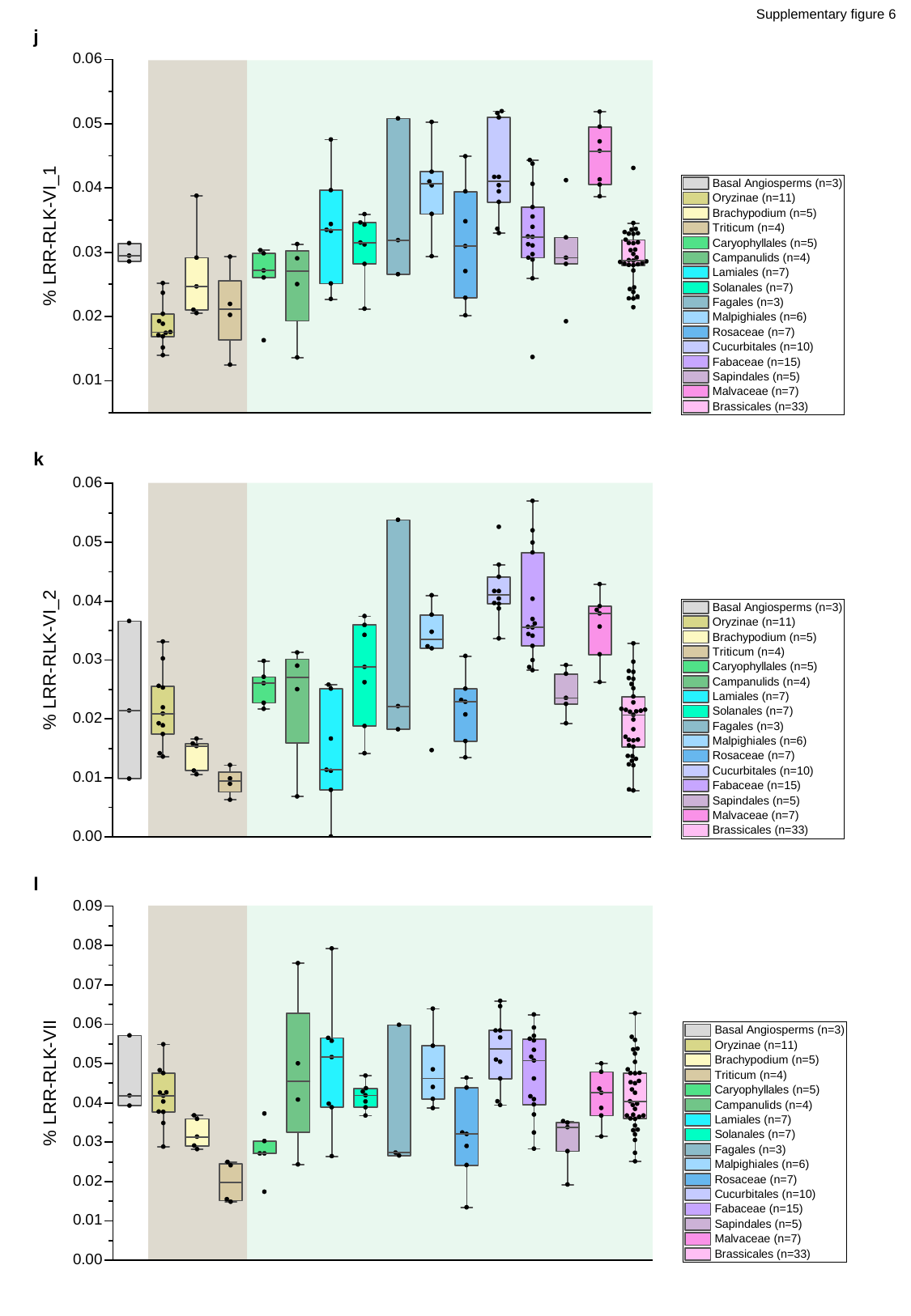

Supplementary figure 6
j
k
l

### Slide 29
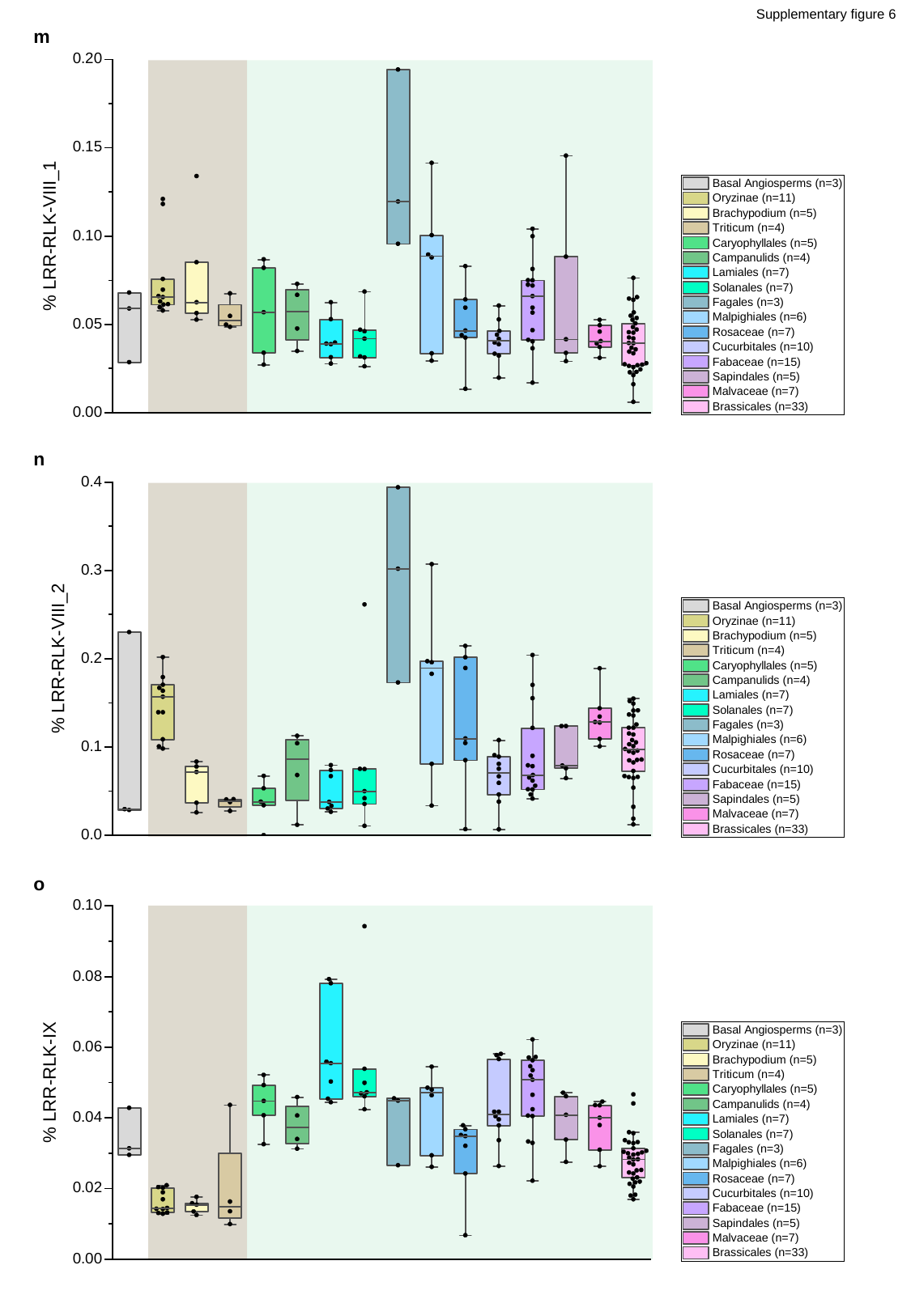

Supplementary figure 6
m
n
o

### Slide 30
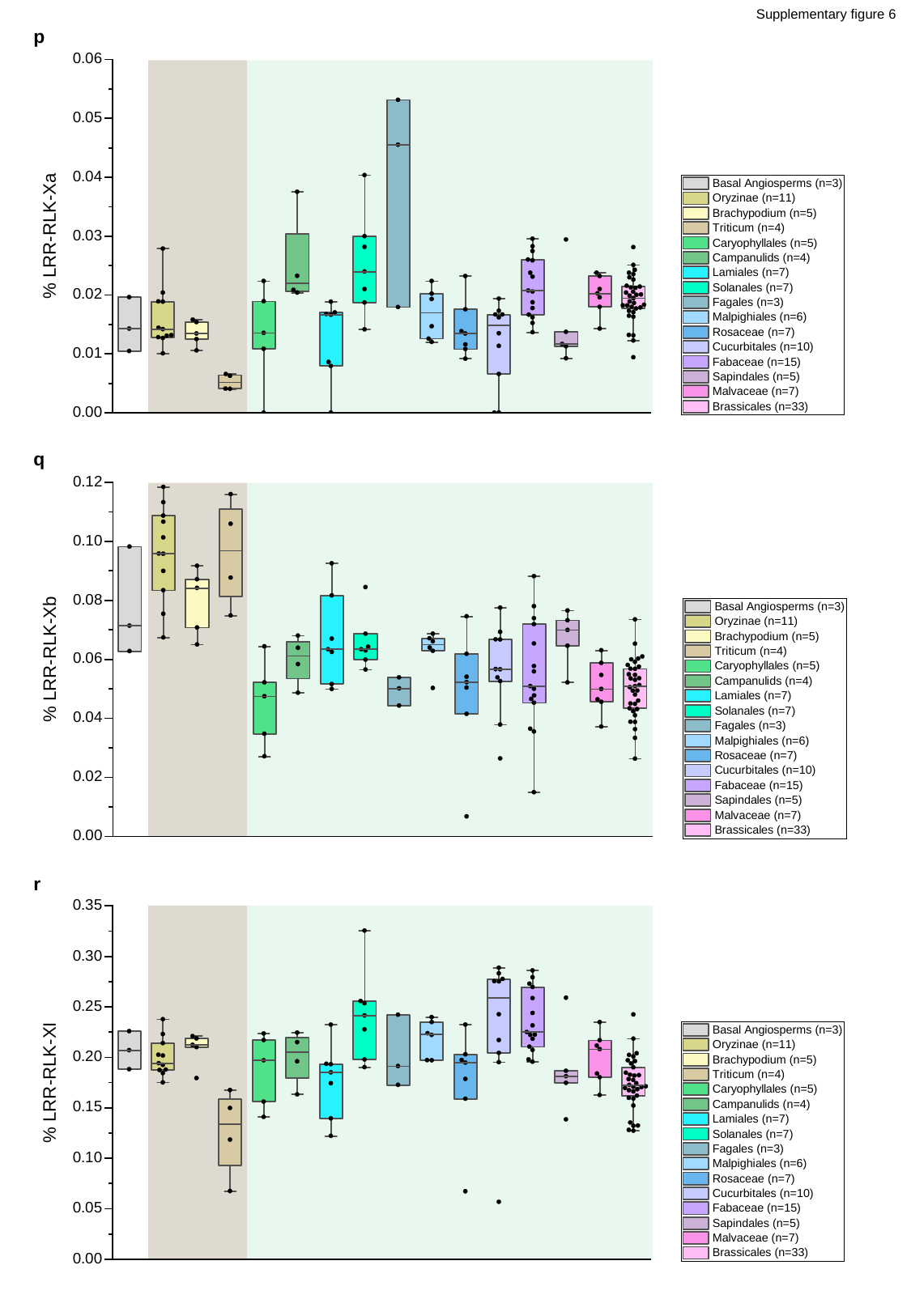

Supplementary figure 6
p
q
r

### Slide 31
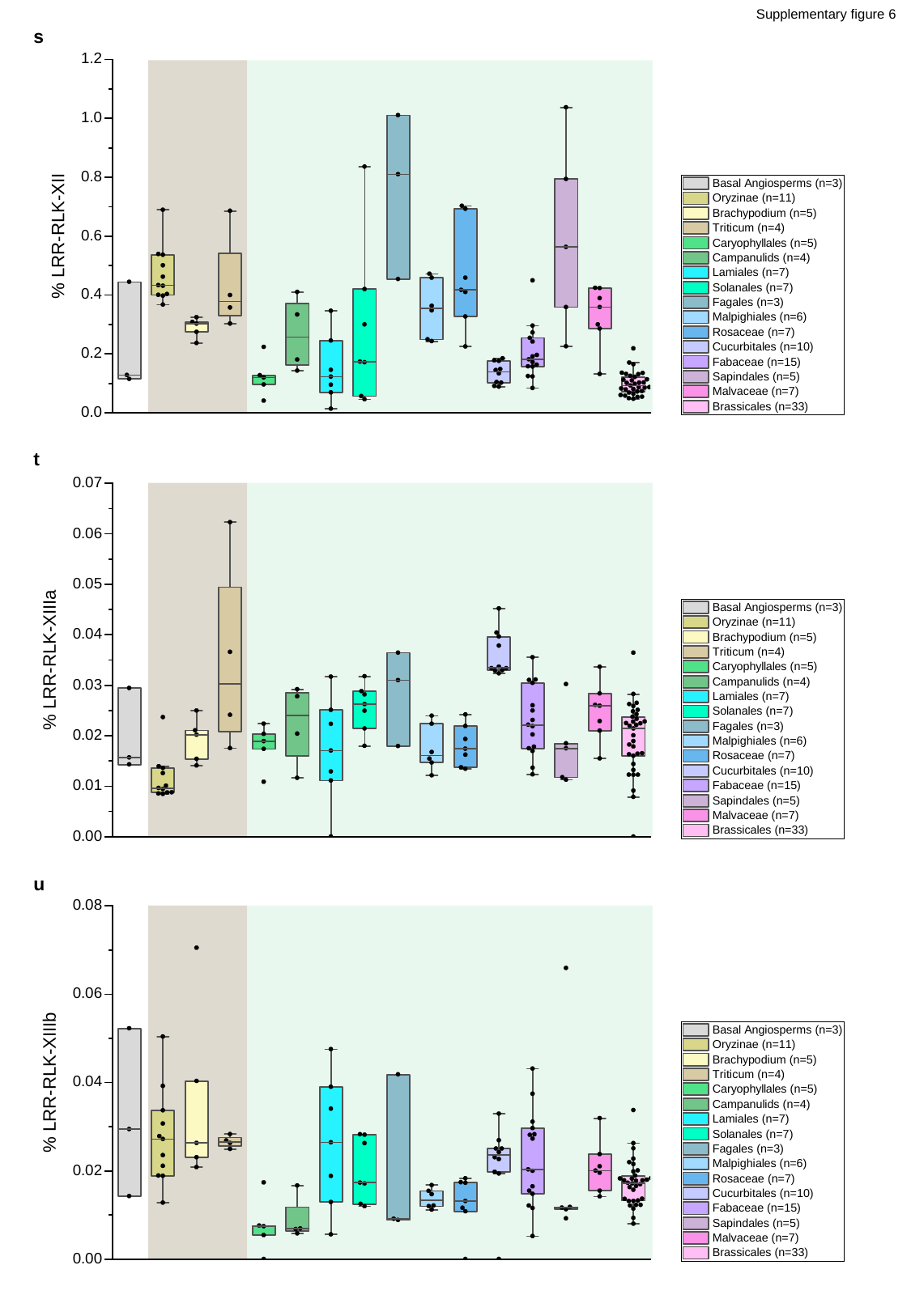

Supplementary figure 6
s
t
u

### Slide 32
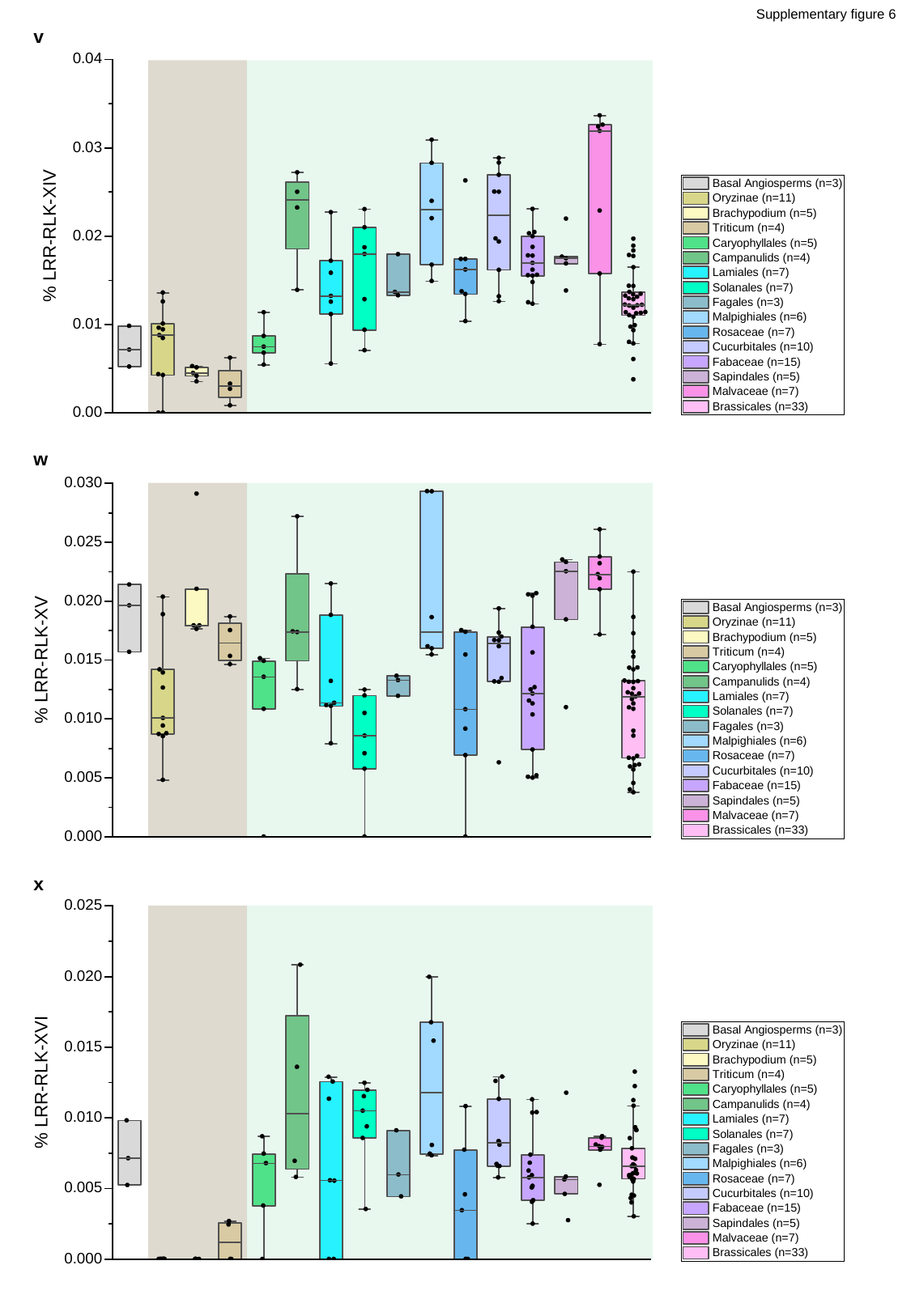

Supplementary figure 6
v
w
x

### Slide 33
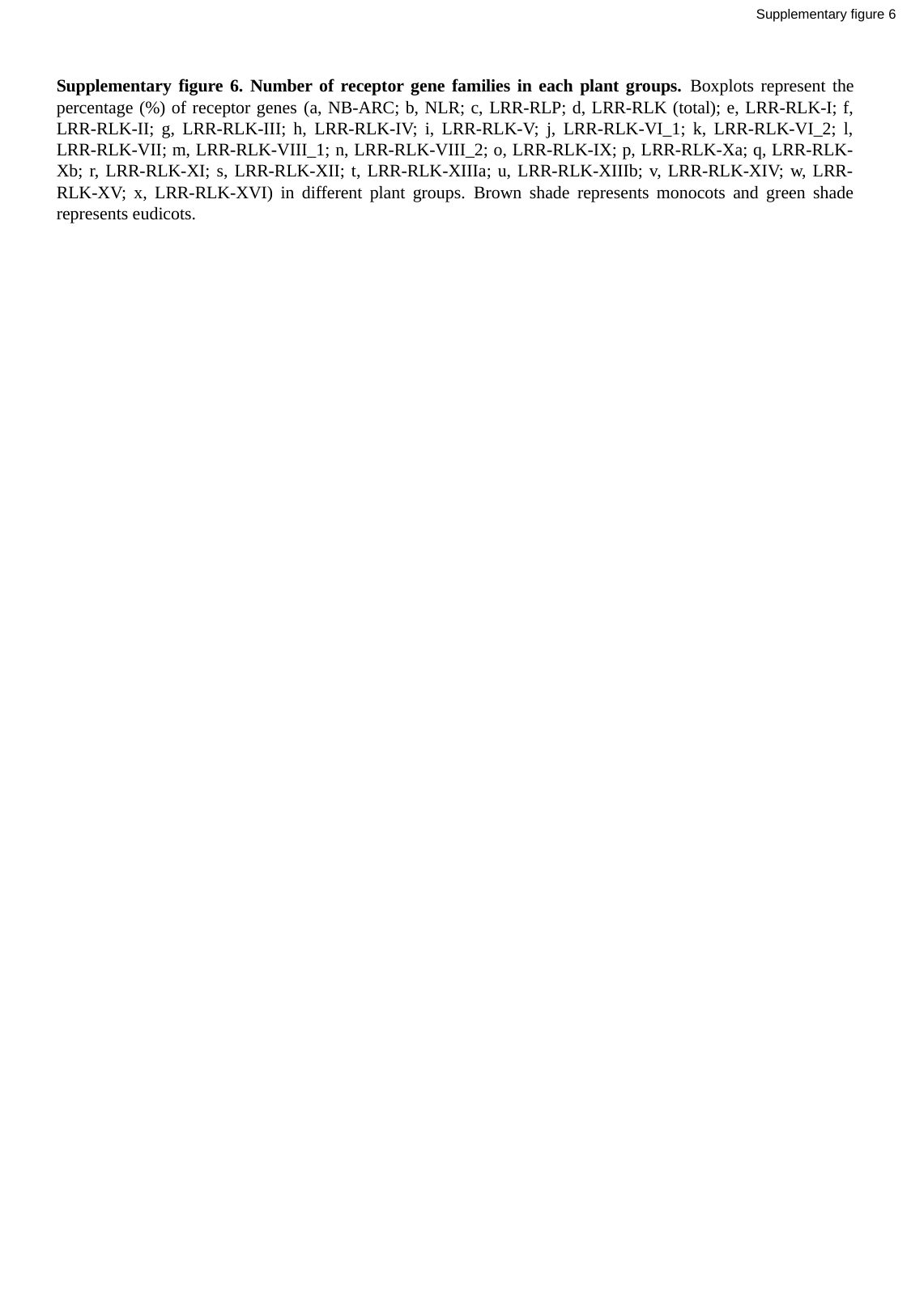

Supplementary figure 6
Supplementary figure 6. Number of receptor gene families in each plant groups. Boxplots represent the percentage (%) of receptor genes (a, NB-ARC; b, NLR; c, LRR-RLP; d, LRR-RLK (total); e, LRR-RLK-I; f, LRR-RLK-II; g, LRR-RLK-III; h, LRR-RLK-IV; i, LRR-RLK-V; j, LRR-RLK-VI_1; k, LRR-RLK-VI_2; l, LRR-RLK-VII; m, LRR-RLK-VIII_1; n, LRR-RLK-VIII_2; o, LRR-RLK-IX; p, LRR-RLK-Xa; q, LRR-RLK-Xb; r, LRR-RLK-XI; s, LRR-RLK-XII; t, LRR-RLK-XIIIa; u, LRR-RLK-XIIIb; v, LRR-RLK-XIV; w, LRR-RLK-XV; x, LRR-RLK-XVI) in different plant groups. Brown shade represents monocots and green shade represents eudicots.

### Slide 34
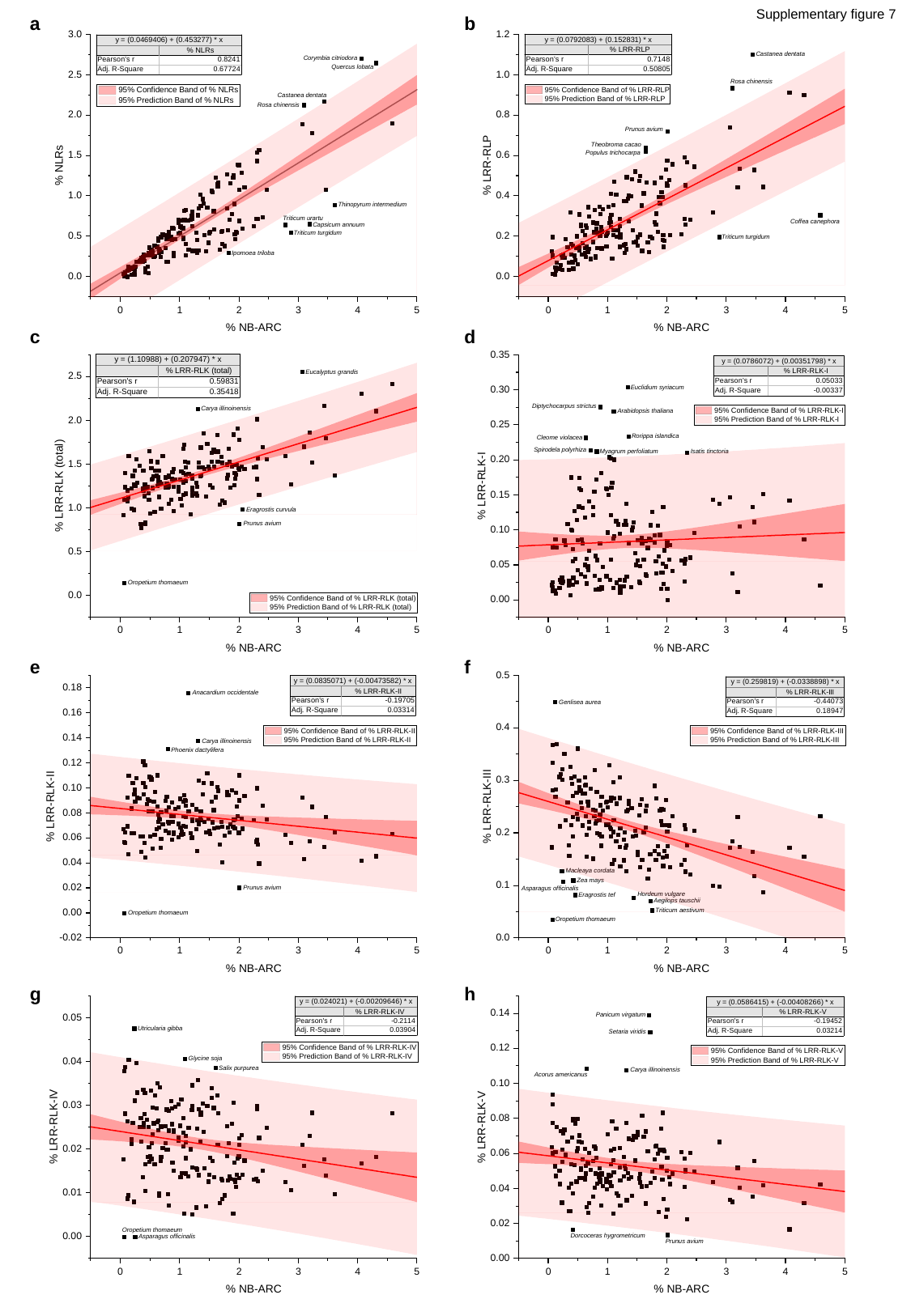

Supplementary figure 7
a
b
c
d
e
f
g
h

### Slide 35
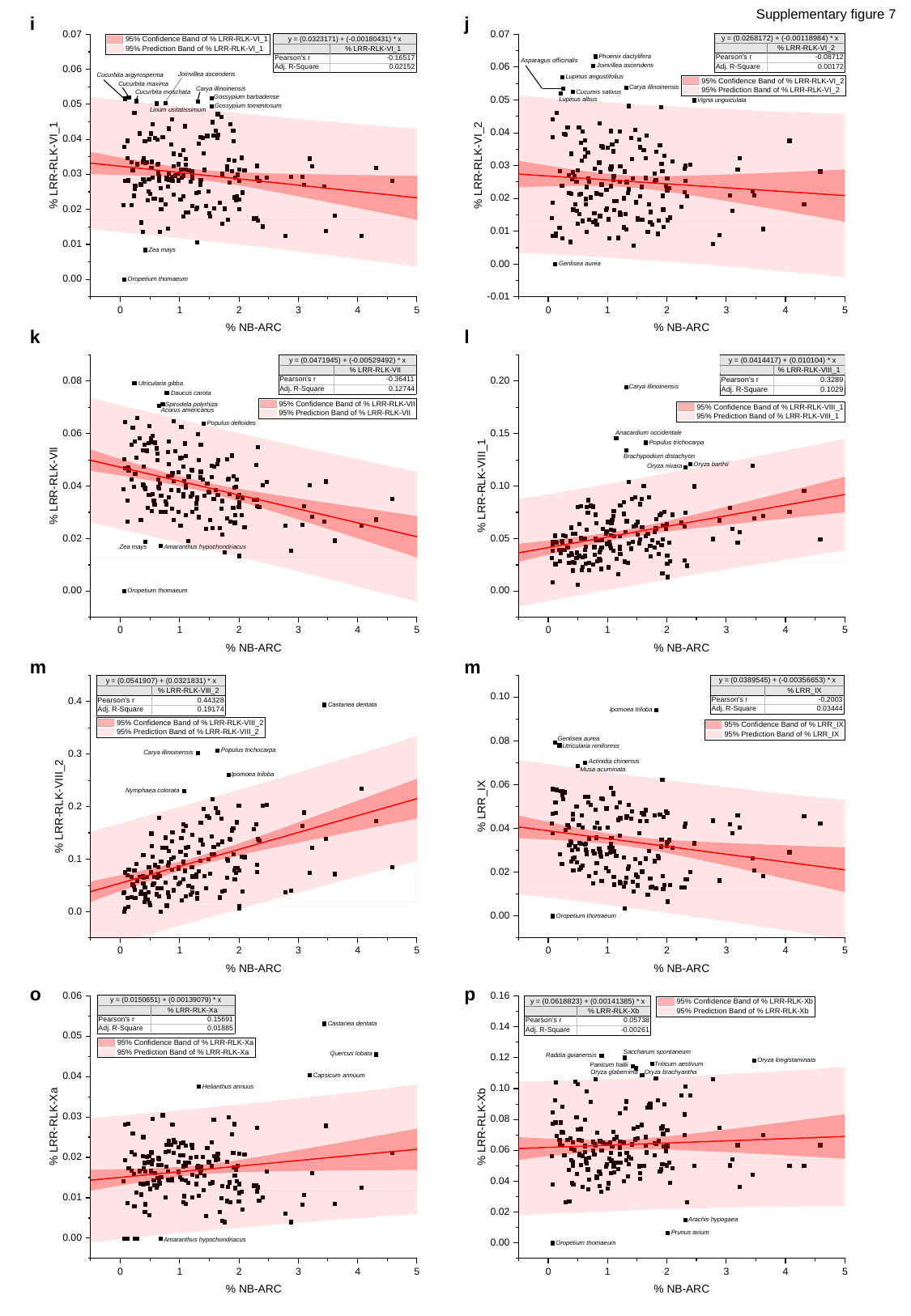

Supplementary figure 7
i
j
k
l
m
m
o
p

### Slide 36
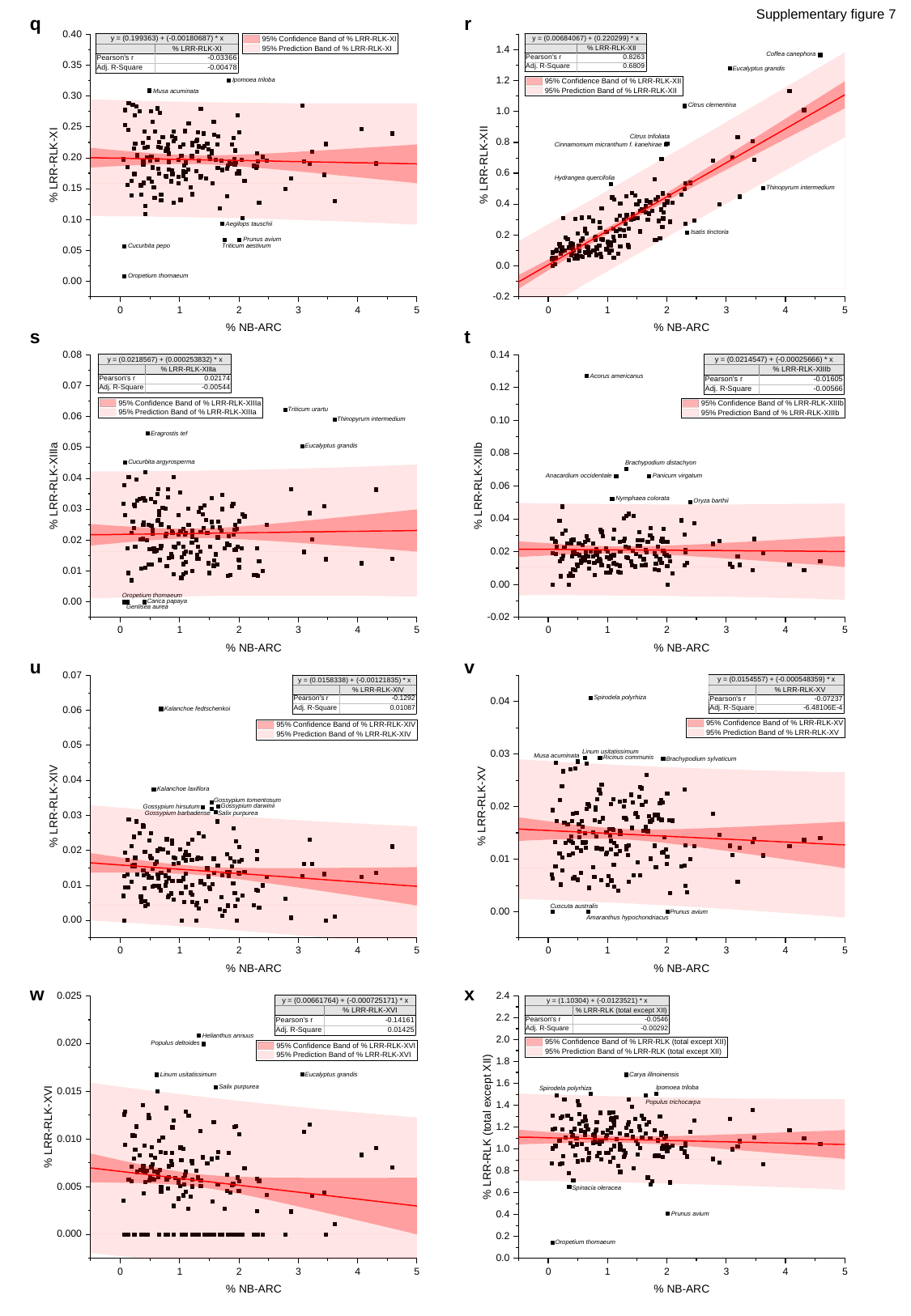

Supplementary figure 7
q
r
s
t
u
v
w
x

### Slide 37
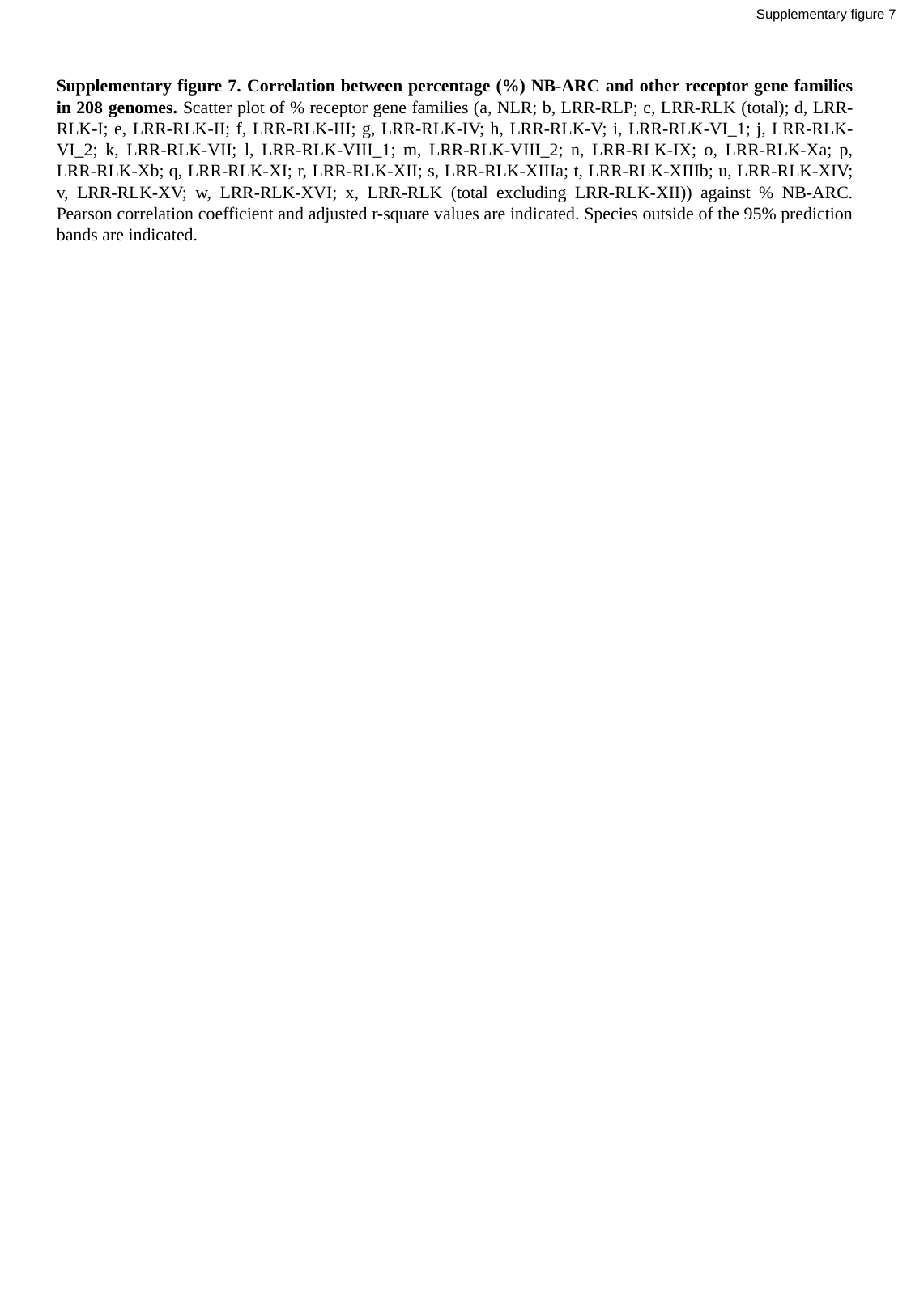

Supplementary figure 7
Supplementary figure 7. Correlation between percentage (%) NB-ARC and other receptor gene families in 208 genomes. Scatter plot of % receptor gene families (a, NLR; b, LRR-RLP; c, LRR-RLK (total); d, LRR-RLK-I; e, LRR-RLK-II; f, LRR-RLK-III; g, LRR-RLK-IV; h, LRR-RLK-V; i, LRR-RLK-VI_1; j, LRR-RLK-VI_2; k, LRR-RLK-VII; l, LRR-RLK-VIII_1; m, LRR-RLK-VIII_2; n, LRR-RLK-IX; o, LRR-RLK-Xa; p, LRR-RLK-Xb; q, LRR-RLK-XI; r, LRR-RLK-XII; s, LRR-RLK-XIIIa; t, LRR-RLK-XIIIb; u, LRR-RLK-XIV; v, LRR-RLK-XV; w, LRR-RLK-XVI; x, LRR-RLK (total excluding LRR-RLK-XII)) against % NB-ARC. Pearson correlation coefficient and adjusted r-square values are indicated. Species outside of the 95% prediction bands are indicated.

### Slide 38
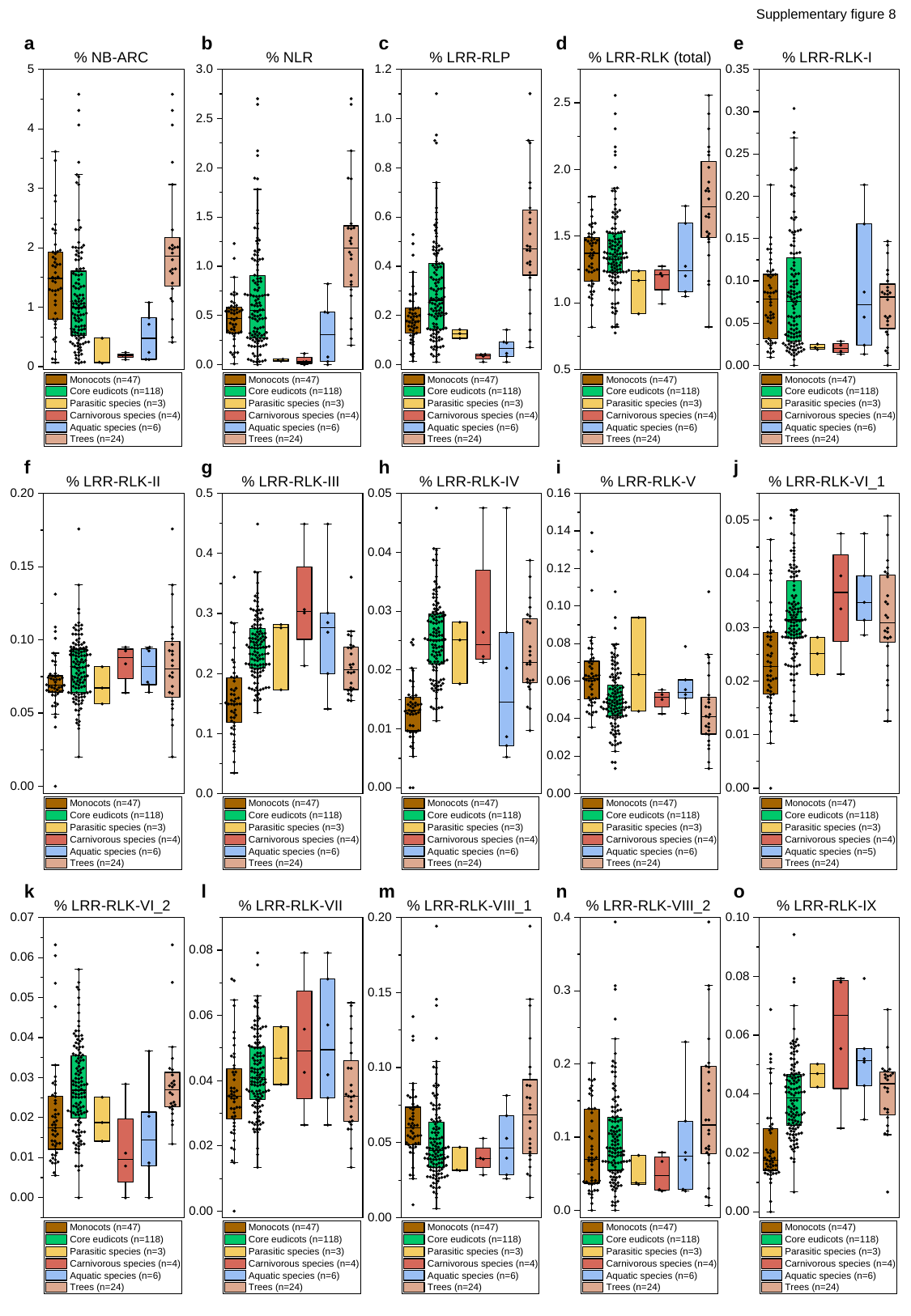

Supplementary figure 8
a
b
c
d
e
f
g
h
i
j
k
l
m
n
o

### Slide 39
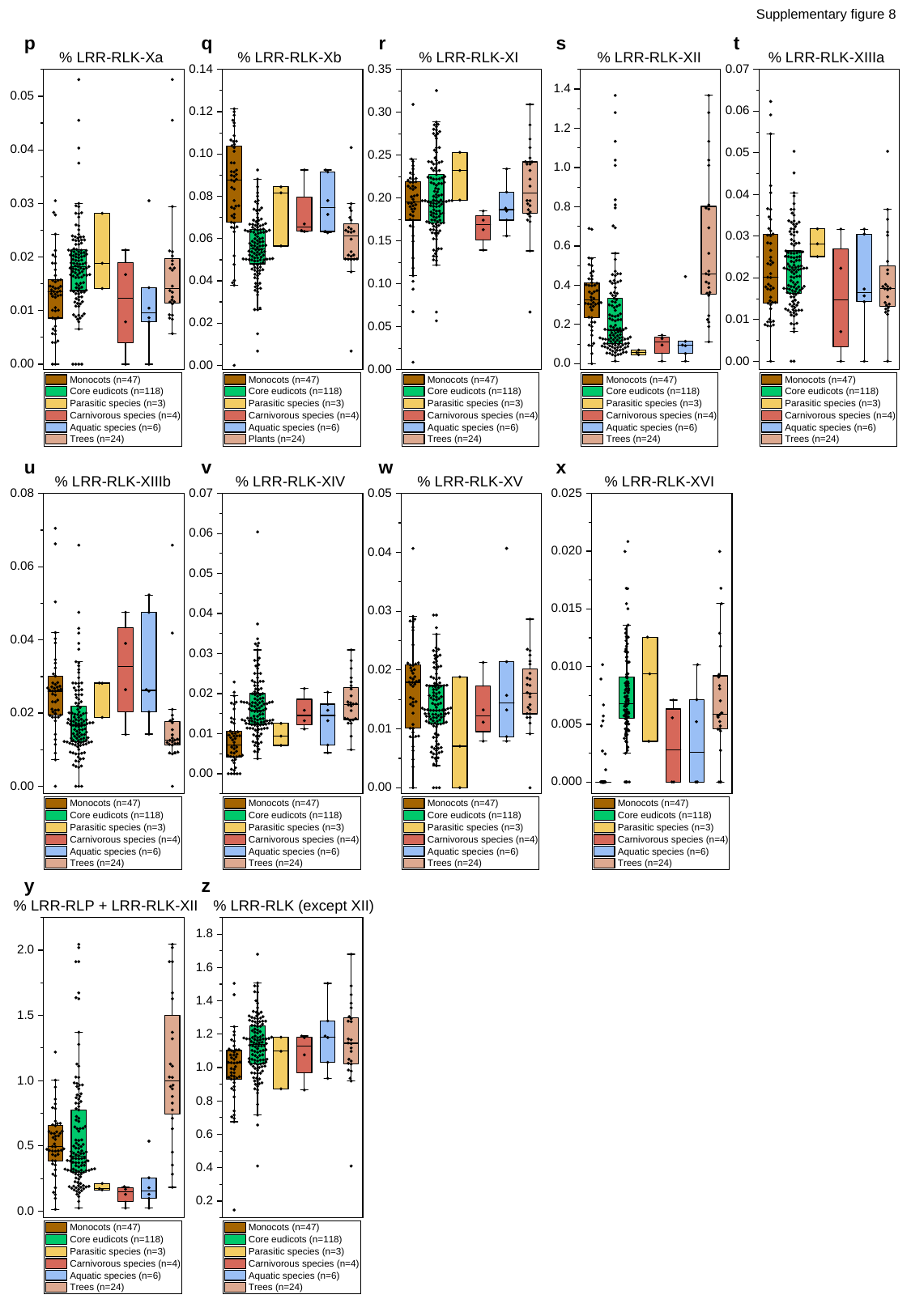

Supplementary figure 8
p
q
r
s
t
u
v
w
x
y
z

### Slide 40
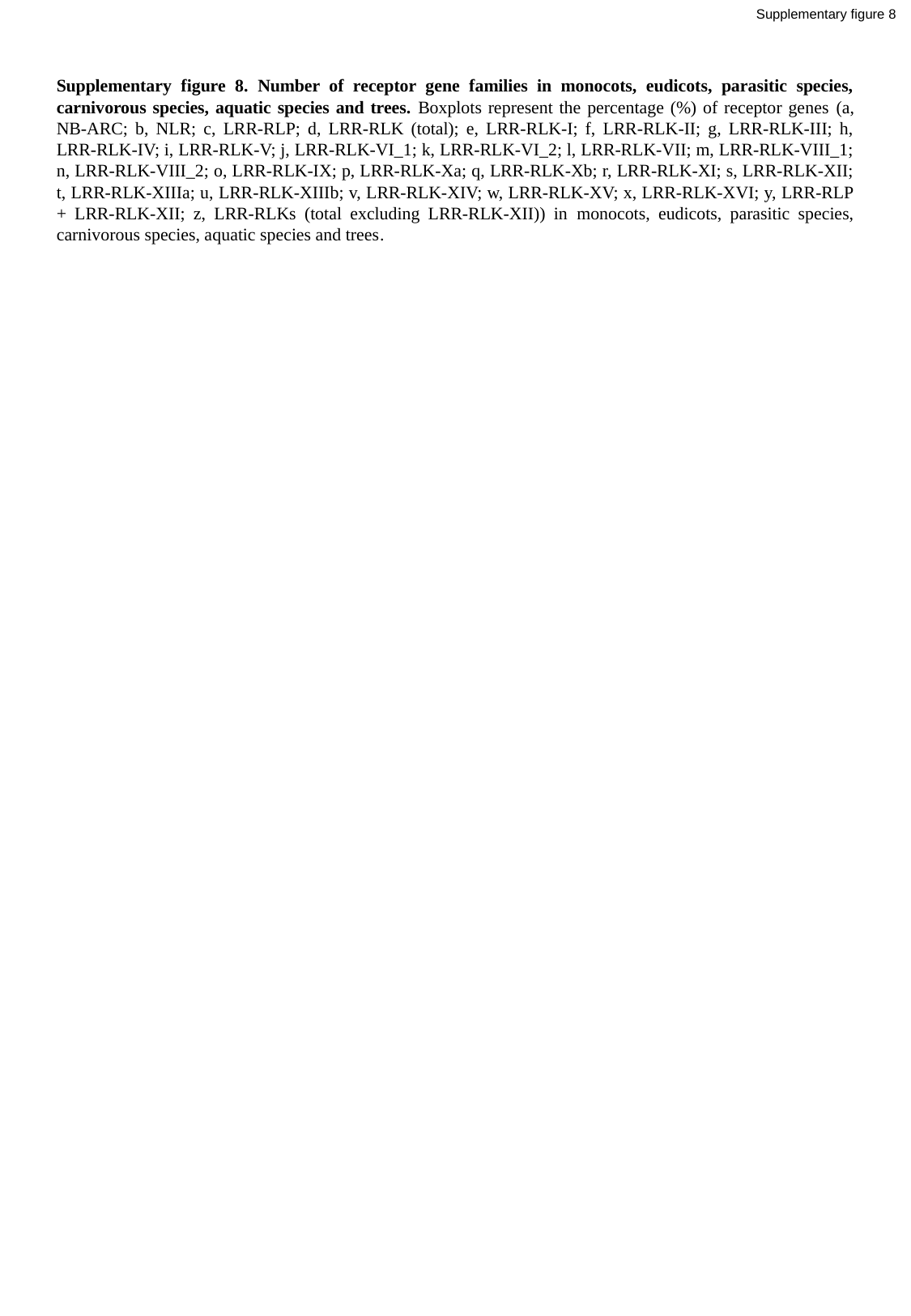

Supplementary figure 8
Supplementary figure 8. Number of receptor gene families in monocots, eudicots, parasitic species, carnivorous species, aquatic species and trees. Boxplots represent the percentage (%) of receptor genes (a, NB-ARC; b, NLR; c, LRR-RLP; d, LRR-RLK (total); e, LRR-RLK-I; f, LRR-RLK-II; g, LRR-RLK-III; h, LRR-RLK-IV; i, LRR-RLK-V; j, LRR-RLK-VI_1; k, LRR-RLK-VI_2; l, LRR-RLK-VII; m, LRR-RLK-VIII_1; n, LRR-RLK-VIII_2; o, LRR-RLK-IX; p, LRR-RLK-Xa; q, LRR-RLK-Xb; r, LRR-RLK-XI; s, LRR-RLK-XII; t, LRR-RLK-XIIIa; u, LRR-RLK-XIIIb; v, LRR-RLK-XIV; w, LRR-RLK-XV; x, LRR-RLK-XVI; y, LRR-RLP + LRR-RLK-XII; z, LRR-RLKs (total excluding LRR-RLK-XII)) in monocots, eudicots, parasitic species, carnivorous species, aquatic species and trees.

### Slide 41
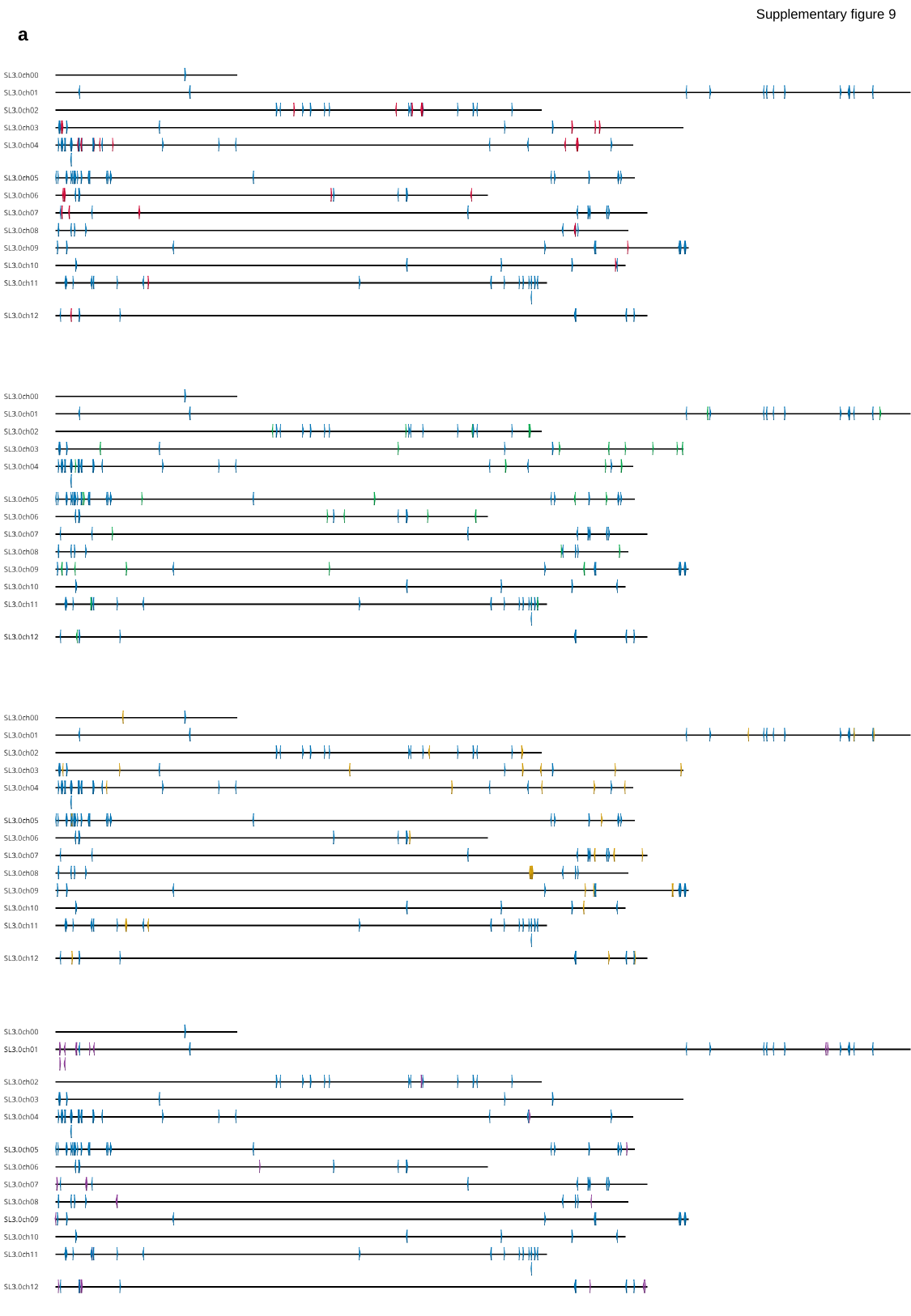

Supplementary figure 9
a

### Slide 42
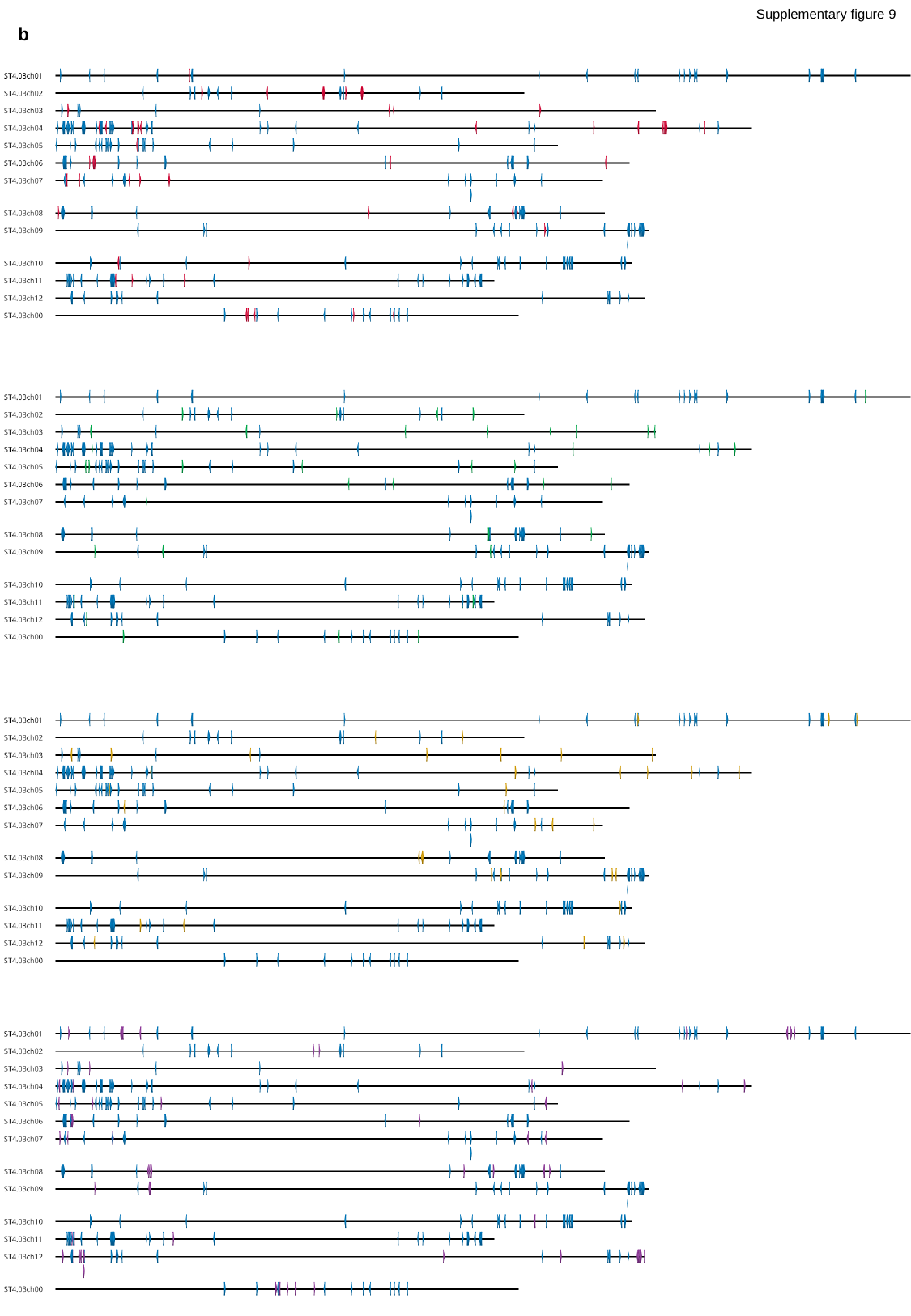

Supplementary figure 9
b

### Slide 43
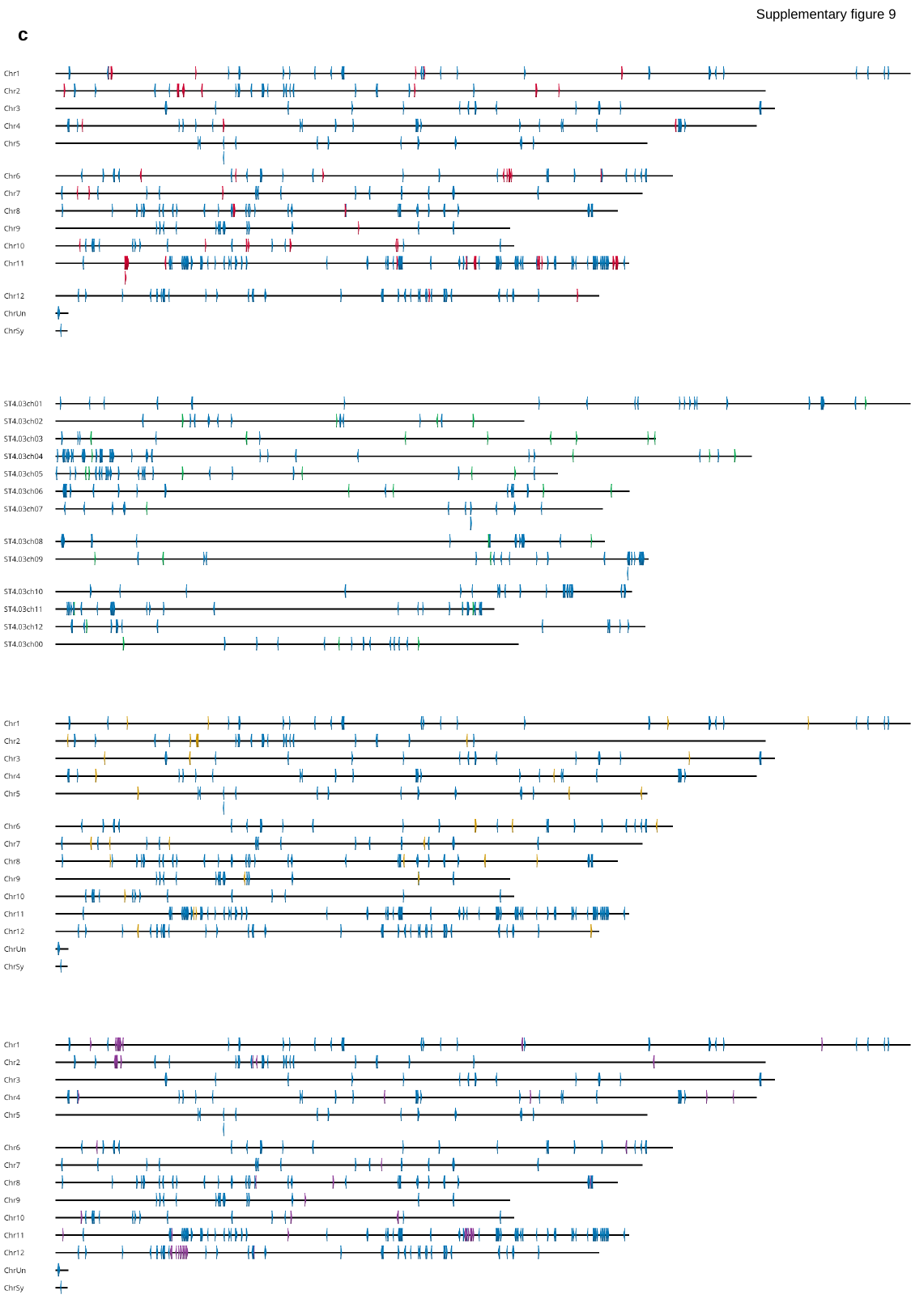

Supplementary figure 9
c

### Slide 44
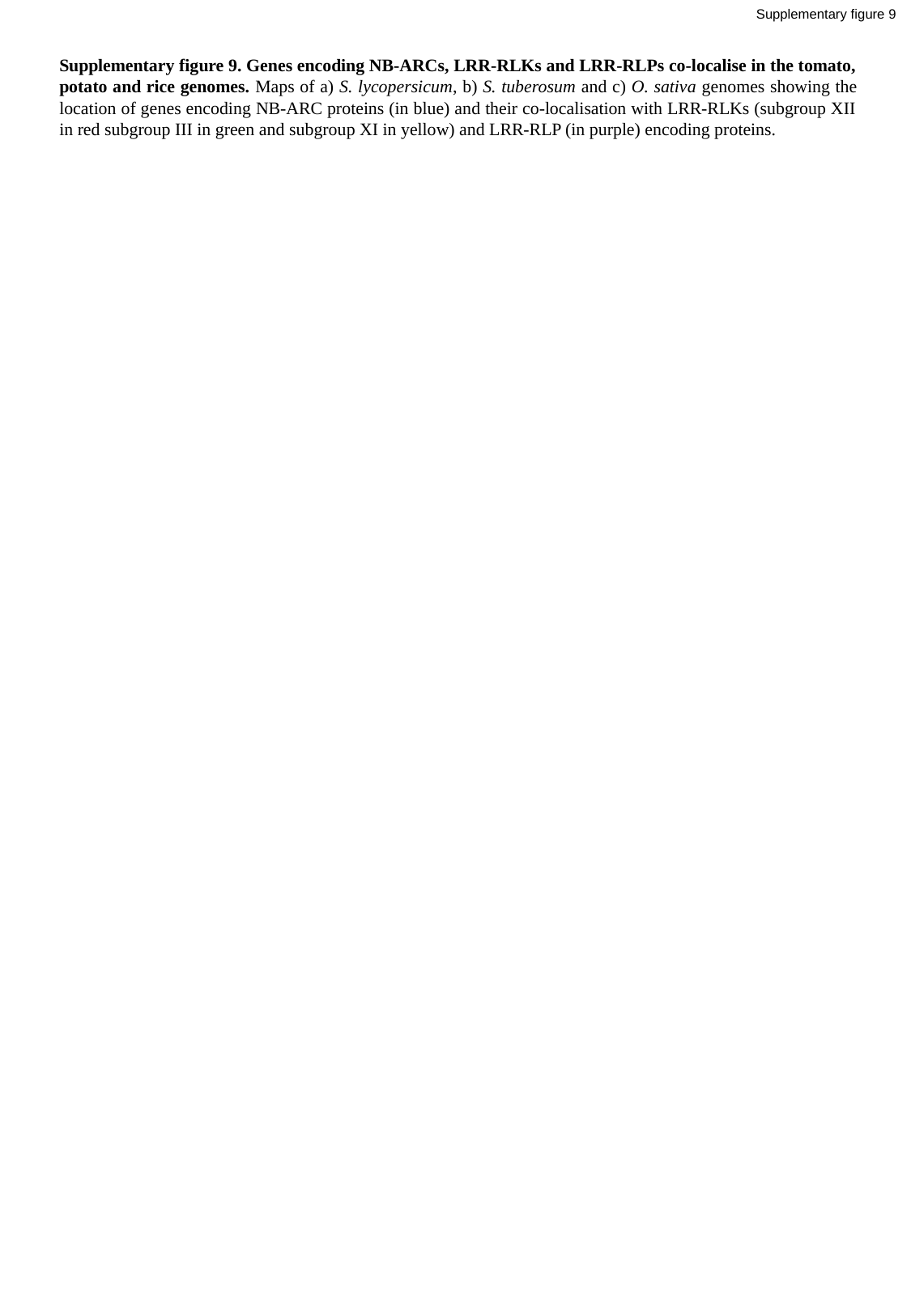

Supplementary figure 9
Supplementary figure 9. Genes encoding NB-ARCs, LRR-RLKs and LRR-RLPs co-localise in the tomato, potato and rice genomes. Maps of a) S. lycopersicum, b) S. tuberosum and c) O. sativa genomes showing the location of genes encoding NB-ARC proteins (in blue) and their co-localisation with LRR-RLKs (subgroup XII in red subgroup III in green and subgroup XI in yellow) and LRR-RLP (in purple) encoding proteins.
